# The cost of information acquisition by natural selection

**DOI:** 10.1101/2022.07.02.498577

**Authors:** Ryan Seamus McGee, Olivia Kosterlitz, Artem Kaznatcheev, Benjamin Kerr, Carl T. Bergstrom

## Abstract

Natural selection enriches genotypes that are well-adapted to their environment. Over successive generations, these changes to the frequencies of types accumulate information about the selective conditions. Thus, we can think of selection as an algorithm by which populations acquire information about their environment. Kimura (1961) pointed out that every bit of information that the population gains this way comes with a minimum cost in terms of unrealized fitness (substitution load). Due to the gradual nature of selection and ongoing mismatch of types with the environment, a population that is still gaining information about the environment has lower mean fitness than a counter-factual population that already has this information. This has been an influential insight, but here we find that experimental evolution of *Escherichia coli* with mutations in a RNA polymerase gene (*rpoB*) violates Kimura’s basic theory. To overcome the restrictive assumptions of Kimura’s substitution load and develop a more robust measure for the cost of selection, we turn to ideas from computational learning theory. We reframe the ‘learning problem’ faced by an evolving population as a population versus environment (PvE) game, which can be applied to settings beyond Kimura’s theory – such as stochastic environments, frequency-dependent selection, and arbitrary environmental change. We show that the learning theoretic concept of ‘regret’ measures relative lineage fitness and rigorously captures the efficiency of selection as a learning process. This lets us establish general bounds on the cost of information acquisition by natural selection. We empirically validate these bounds in our experimental system, showing that computational learning theory can account for the observations that violate Kimura’s theory. Finally, we note that natural selection is a highly effective learning process in that selection is an asymptotically optimal algorithm for the problem faced by evolving populations, and no other algorithm can consistently outperform selection in general. Our results highlight the centrality of information to natural selection and the value of computational learning theory as a perspective on evolutionary biology.

---

Living organisms are astonishingly complex and intricately adapted to their environments. To implement adaptations to particular environments, organisms require extensive information about the environmental conditions and how to function in response. Much of this information is carried by the genome. The core insight of the modern synthesis is that this adaptive genetic information is acquired by the process of natural selection acting on genetic variation.

Natural selection shifts the frequencies of types in a population such that types that are well-suited for the environment become more common, and less fit types are eliminated. Over successive generations, the effects of selection leave an imprint of the selective conditions on the population’s composition, and the relative frequencies of types provide an increasing amount of information about the population’s environmental history. In this way, we might view selection as a learning process through which the population acquires adaptive information. Population genetics offers a large body of theory describing how genetic variance changes in the process of evolution, but we lack correspondingly rich theory for how adaptive information changes as a consequence of these same dynamics.

Motoo Kimura was among the first to consider natural selection as an information acquisition process in a formal sense. Kimura (1961) proposed a simple relationship between information gain by natural selection and the long-term growth of a population: As selection updates the composition of the population to better match the environment, information about the environment is acquired and the mean fitness of the population increases. But selection is gradual and mean fitness remains suboptimal until a fully optimal composition is reached. Kimura noted that the amount of information gained in a simple allele substitution is proportional to the loss of potential fitness incurred by the incremental nature of selection. This result led Kimura to suppose that there is a cost of selection that limits how much information can be gained by this process.

Kimura’s result points to a fundamental relationship between two essential quantities in evolution—fitness and information— but this connection has seen limited development since (Frank 2009, 2012, Donaldson-Matasci et al. 2010, Rivoire and Leibler 2011, Adami 2012, Hledík et al. 2021). Here we test, extend, and reinterpret this theory to illuminate a rigorous and meaningful relationship between information and fitness. First, we formalize selection as an information acquisition process and review Kimura’s result relating information gain and substitution load. We then test this basic result experimentally, which prompts us to introduce a more general definition of mismatch load for heterogeneous and time-varying environments. We thereby validate a *lower* bound on the cost of selection: the minimum load that a population must incur in order to gain a given quantity of information. We then adopt the frame of natural selection as a learning process in order to clarify and formalize the cost of selection in learning theoretic terms. This allows us to characterize the relative efficiency of selection as an information acquisition process by establishing general *upper* bounds on the cost of selection that hold for any environment, including frequency-dependent and adversarial settings. Our results describe a tight relationship between information gain and lineage fitness.

## Natural selection as an information acquisition process

Learning is the process of iteratively updating a hypothesis in light of new evidence. Consider a population that consists of a number of asexual replicators of varying types (e.g., genotypes). Each alternative type represents a strategy for how to function and survive, and the distribution of type frequencies ***p***^*t*^ can be seen as the population’s hypothesis about which type is most suited for the environment at time *t*. Natural selection acts as a learning process that updates this hypothesis. The fitness *W*_*i*_ of the *i*th type provides evidence about its suitability for the current state of the environment, and selection changes the frequency *p*_*i*_ of each type accordingly following the well-known replicator dynamics (Appendix A.1.1):

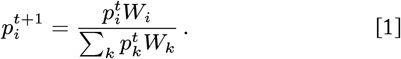

The process by which selection updates a population’s type distribution is formally analogous to Bayesian learning (Harper 2009b, Shalizi 2009, Campbell 2016, Watson and Szathmáry 2016, Czégel et al. 2020) (Appendix C.1). Selection increases the frequency of types with high relative fitness in the same way that Bayes’ rule increases the weight of alternatives that give high relative likelihood to the observed evidence. The population’s hypothesis regarding the fit of types to the environment is refined over time as new information about which types have obtained high fitness is encoded into the frequency distribution.

We can measure how much a population learns by measuring how much its hypothesis changes.The Kullback-Leibler divergence between the population’s initial type distribution ***p***^0^ and its updated distribution ***p***^*T*^ quantifies the amount of **information gain** *I*^*T*^ that selection provides over a learning period of duration *T* (Figure 1a):

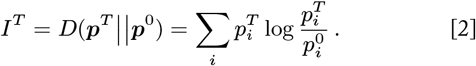

While other processes such as drift, mutation, and migration may change type frequencies, we focus here on changes due to selection. Selection is unique among these processes in that the type frequency changes it enacts are, by definition, explicitly dependent on the environment. Selection establishes an adaptive matching between types and environments such that observing a particular type in a given population reduces uncertainty about the kind of environment it occupies (Shea 2007, Bergstrom and Rosvall 2011). In this sense, frequency changes attributable to selection directly contribute to an increase in adaptive information about the environment, whereas frequency changes attributable to drift or other processes do not (McGee and Bergstrom 2022) (Appendix B.2.3).

**Fig. 1.**
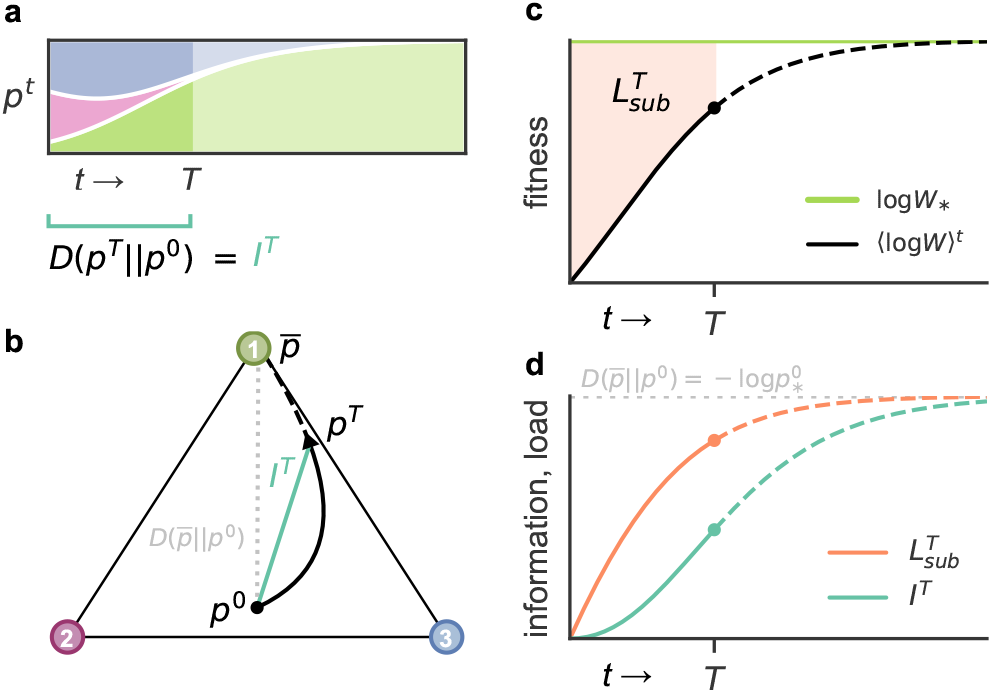
Selection accumulates information gain and substitution load. Natural selection changes the frequencies of types in a population according to their relative fitnesses. **(a)** Change in the population’s type frequency distribution over time is depicted by a Muller plot, where the height of each colored band represents the frequency of the corresponding type at a given time (type colors are indicated in (b)). The population’s information gain *I*^*T*^ at time *T* is measured by the KL divergence of the population’s state ***p***^*T*^ from its initial state ***p***^0^. **(b)** The black trajectory through the simplex tracks the composition of the population over time as selection moves the population from its initial type frequency distribution ***p***^0^ toward fixation of the optimal Type 1 (green vertex), which is an evolutionarily stable state (ESS) 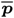. The teal line segment connecting the population’s initial state with its trajectory through the simplex provides graphical intuition about how information gain changes over time (note that KL divergence is not a true distance metric). The gray dotted line represents the initial potential information, which is defined as the KL divergence of the ESS from the population’s initial state. **(c)** As selection proceeds in this fixed environment, the mean fitness of the population increases (black line) as suboptimal types decrease in frequency. As the population approaches fixation of Type 1, the mean fitness converges on the optimal fitness of this type (green line). The substitution load at time *T* measures the cumulative depression in population mean fitness below the optimal fitness level up to that time, which corresponds to the area of the orange shaded region. **(d)** The accumulation of information gain (teal line) and substitution load (orange line) are plotted over the course of selection. Substitution load and information gain converge on the same value as the population approaches fixation (Proposition 1), and substitution load always exceeds information gain (Proposition 2).

As a population evolves with respect to a particular environment, selection moves the population toward an **evolutionarily stable state** (ESS) 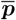 that maximizes expected fitness for the current conditions (Maynard Smith 1982, Eshel and Feldman 1984, Hammerstein 1996). The divergence 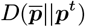 of the ESS from the population’s current type distribution defines the **potential information** of the system: the information gain that is still available from the population’s current state (Figure 1b). As selection proceeds, information is gained (Figure 1d) and the potential information continually decreases (Appendix B.2.2). In fact, the route that selection takes follows an information gradient, where each update shifts the type distribution in the direction that maximizes information gain relative to the population’s current state (Harper 2009a, Harper and Fryer 2015). When the population reaches an equilibrium composition, it has gained all of the information that can be gained about the current environment. Therefore natural selection can be understood at its most fundamental as an information acquisition process, both in its outcomes and in its underlying dynamics.

### Fitness loss associated with information gain

Populations do not gain information for free. Just as we learn from our mistakes, populations learn from the shortcomings of poorly adapted types. Differential fitnesses provide the evidence that drives the population’s learning, but selection does not immediately pivot to the highly fit types. Instead, it incorporates this information by gradually adjusting allele frequencies over time. A population that is still gaining information about the environment has lower mean fitness than a population that already has this information.

Haldane (1937, 1957) and Kimura (1960) introduced the concept of substitution load to describe the relative cost of using selection to fix beneficial alleles in a population. Haldane and Kimura considered the simple case of an allele substitution in an infinite haploid population with multiple segregating types. The optimal type with the highest fitness increases in frequency toward fixation, but until the substitution is complete the presence of less-fit types holds the mean fitness of the population below the maximum. The **substitution load** 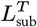 measures the cumulative difference between the maximum growth rate of an ideal monomorphic population of the optimal type and the actual population’s average growth rate over *T* generations of selection (Figure 1c, Appendix D.1.1):

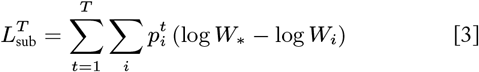

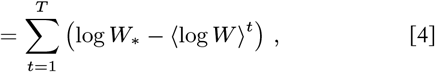

where *W*_∗_ and *W*_*i*_ denote the fitnesses of the optimal and *i*th types, respectively. The log fitness log*W*_*i*_ corresponds to the growth rate of the *i*th type, and 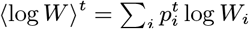 denotes the population’s average growth rate (Appendix A.1.1). Substitution load gives the total fold loss of potential growth that a population suffers by evolving its composition incrementally relative to a population that grows optimally all along.

Kimura (1961) noticed an interesting connection: the total information gain accumulated in a complete allele substitution is equal to the total substitution load incurred during that process. Irrespective of the strength of selection and the speed of the substitution process, the information gain and substitution load ultimately converge on the same value, which depends only on the initial frequency of the optimal type 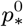 (Figure 1d, Appendix D.1.1).

#### Proposition 1.

*(Kimura 1961)*

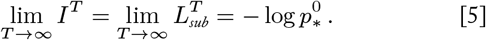

In addition, we prove here that substitution load always exceeds information gain throughout the course of selection, with equality in the limit of fixation **(Figure 1d)**

#### Proposition 2.

*(Appendix D.4.3)*

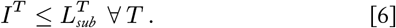

This implies that in order to gain a given amount of information, there is a minimum load that must be paid to achieve the type substitutions necessary to encode that information. In other words, the process of selection acquiring one bit of information, which represents a twofold reduction in uncertainty about the environment, requires at least a two-fold depression in mean fitness along the way. A population can either grow optimally (i.e., have no less-fit types) or acquire information, but not both.

### Measuring load in realistic environments

Kimura’s theory predicts a relationship between two primary quantities in evolutionary biology—fitness and information. If this relationship is fundamental to natural selection, then we should expect it to be readily observed in empirical populations, but these predictions have yet to be validated directly. While Kimura’s theory is clear and self-consistent, it was developed under a number of simplifying assumptions. It is important to confront such theory with the real world to test its applicability to natural populations and to reveal unexpected ways in which the theory may be revised to accommodate the complexities of real systems.

We set out to test the predictions of Kimura’s theory using an experimental evolution system that closely adheres to the assumptions made in the theory. We used four strains of *Escherichia coli* with distinct mutations in the RNA polymerase *rpoB* gene that confer differing growth rates in a standard growth medium. Each strain was transformed with a plasmid carrying a constitutive fluorescent protein marker for strain identification and cell enumeration. We ran selection experiments with various combinations of strains starting from different initial frequencies (Materials & Methods). Each population was maintained in exponential growth phase for 36 hours (Appendix E.2.1), which was sufficient time for all populations to approach fixation. Information gain was calculated at each time point as the KL-divergence between the population’s current and initial allele frequency distributions. Load was quantified as the cumulative difference between the population’s average growth rate and the growth rate of the optimal allele in each interval.

Results from one of these selection experiments are highlighted in Figure 2. As expected, the optimal allele increases in frequency and approaches fixation (Figure 2b). However, the observed relationship between information and load deviates from the theoretical predictions: the substitution load does not always exceed information gain or converge on information gain at fixation (Figure 2d). Why not? While Kimura’s definition of substitution load assumes a static environment with constant fitnesses, growth rates fluctuated stochastically in our experiments despite maintaining cultures in exponential growth phase (Figure 2c). Given that this assumption cannot be satisfied even in a highly controlled laboratory population, applying this fixed definition of substitution load to natural systems is problematic.

**Fig. 2.**
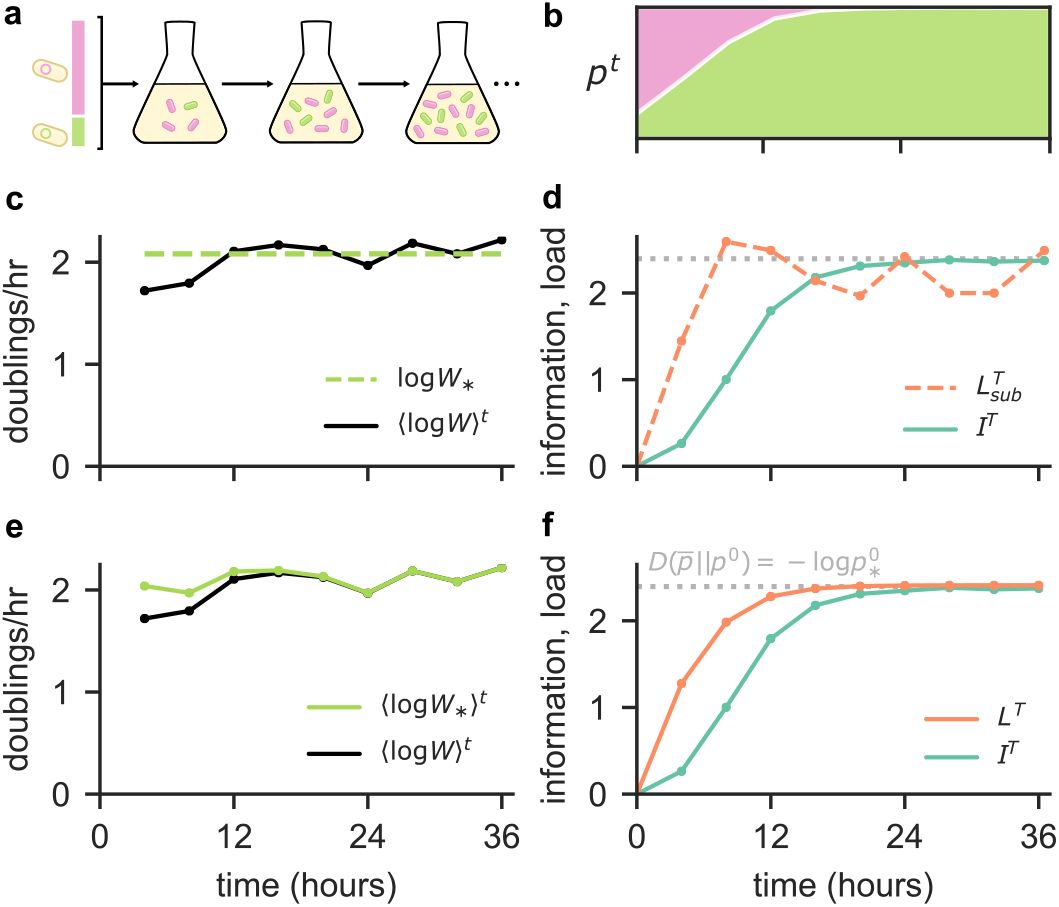
Information gain and load measured in a selection experiment. Results from a single representative selection experiment are presented in detail. **(a)** This selection competition involved a GFP-labeled WT strain and a less-fit RFP-labeled M1 strain with initial frequencies of approximately 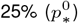 and 75%, respectively. **(b)** Changes in the frequencies of these types over time are depicted by a Muller plot, where the height of the green and pink bands give the frequency of the WT and M1 strains, respectively. The WT strain has the optimal growth rate and selection moves the population toward fixation of this type. **(c)** The mean growth rate of the population (doublings per hour) is plotted over the course of the experiment (black line). Each point represents the estimated average population growth rate over the preceding 4 hour interval. The growth rate of the optimal WT strain was also measured at 4 hour intervals, and the average of these measurements is used as a fixed estimate of this type’s characteristic growth rate (green dashed line). **(d)** Information gain (teal line) and *substitution* load (dashed orange line) measurements are plotted over time. Here substitution load is computed as the difference between the population growth rate (black line in (c)) and the fixed characteristic growth rate of the optimal type (dashed green line in (c)), as per the classical definition. Under this definition, the observed load does not conform to the predictions of Proposition 1 or Proposition 2. **(e)** In addition to the the estimated mean population growth rates over time (black line, same as in (c)), we also plot the estimated growth rates of the optimal WT strain for each interval (green line). While growth rates fluctuate, the growth rate of the optimal allele exceeds the mean population growth rate in every interval, and the population growth rate converges on the optimal rate as it nears fixation of the WT strain. **(f)** Information gain (teal line) and *mismatch* load (orange line) measurements are plotted over time. Using the extended definition of mismatch load, which accommodates time-varying fitnesses, the observed load and information adhere to the the predictions of Proposition 1 and Proposition 2.

These observations led us to extend this concept of load to realistic contexts where conditions and fitnesses change over time. Suppose there is a set of distinct environmental conditions, which may be characterized by abiotic properties, type frequencies, population densities, or other factors (Appendix A.2). Let the probability that a given organism experiences the *j*th condition be denoted by *x*_*j*_, and let the fitness of an organism of the *i*th type in the *j*th condition be given by *W*_*ij*_. Then scenarios with fluctuating conditions can be represented with a probability distribution ***x***^*t*^ that may change arbitrarily over time. For this more general context, we define the **mismatch load** (Appendix D.1.2):

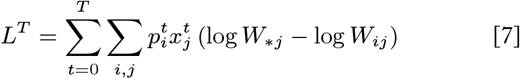

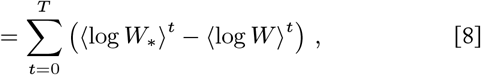

where *W*_∗*j*_ is the fitness of whichever type is optimal in condition *j* (i.e., *W*_∗*j*_= max_*i*_*W*_*ij*_). Mismatch load measures the cumulative loss of potential fitness due to the mismatch of types and environmental conditions. Kimura’s substitution load (Equation 4) is a special case of our mismatch load for an allele substitution in an environment with only a single, unchanging condition.

Although we did our best to create a static and identical environment in our experiments, the environmental conditions (e.g., inoculum size, nutrient concentration, temperature) nevertheless inadvertently change slightly from one interval to the next. This causes growth rates to fluctuate, but all cells in each well-mixed batch culture experience the same conditions, and the identity of the optimal type does not change. This still constitutes an allele substitution scenario, albeit a more noisy one. The theoretical relationship between information gain and load (Proposition 1, Proposition 2) is predicted to hold in such a case, but for our more general mismatch load instead of Kimura’s more particular substitution load (Appendix D.4).

Using the more appropriate definition of mismatch load, our empirical results conform remarkably well to the predictions of theory: information gain is always exceeded by load, and these quantities approach equality as the populations approach fixation (Figure 2f). This relationship is observed in all of our selection competitions regardless of the selection strength or the initial frequency of the optimal allele (Figure 3, Supplemental Figure E6).

**Fig. 3.**
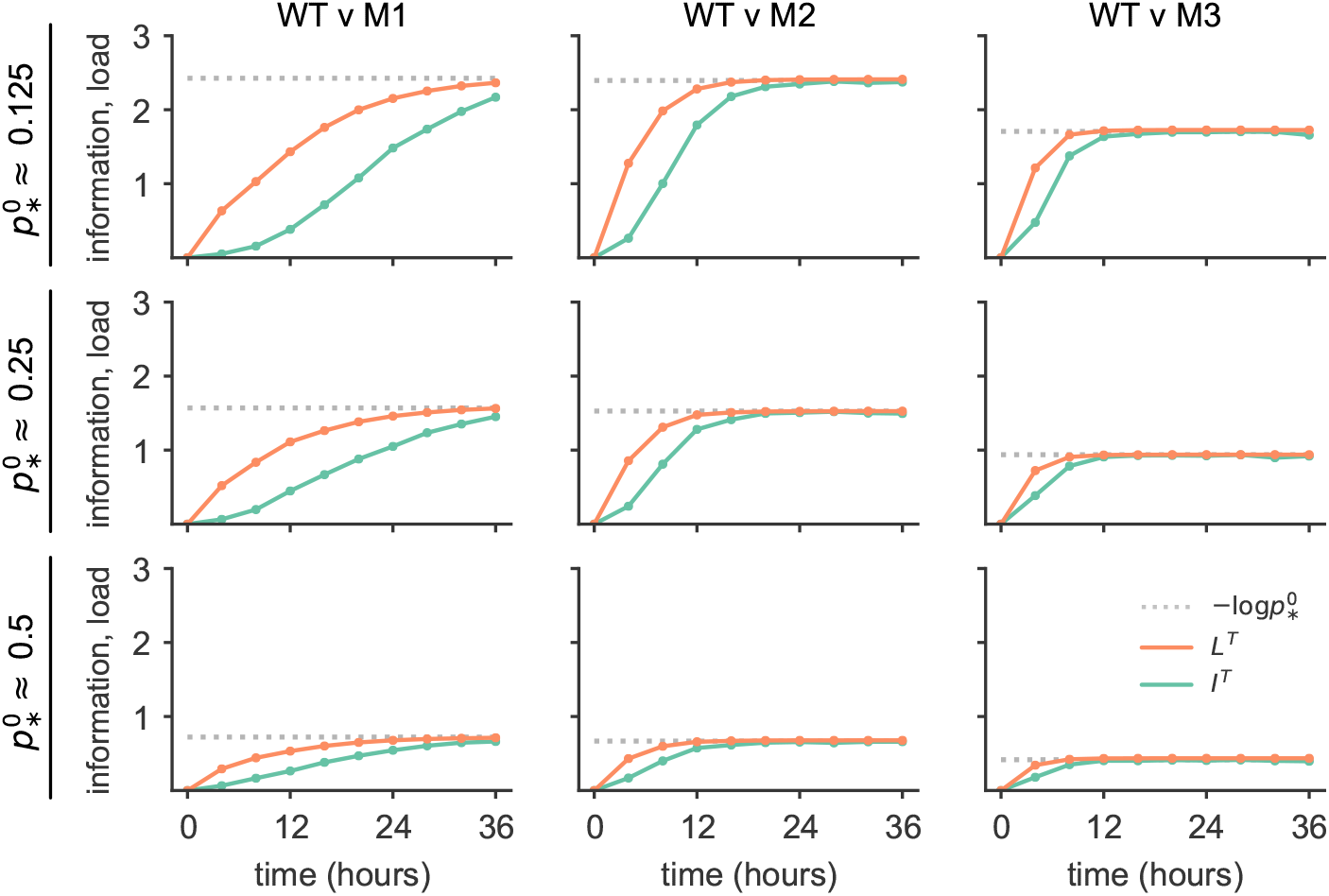
The relationship between load and information predicted by theory is reliably observed in selection experiments. Results from 9 selection competition experiments that involved different combinations of types (column headings) and that initialized batch culture populations with different frequencies of the optimal type 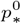 (row headings) are shown. The WT strain had the optimal growth rate and approached fixation in all competitions. Theory predicts that information gain and load converge on the same value, equal to the negative logarithm of the initial frequency of the optimal type (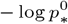, Proposition 1). The convergence value depends on the initial frequencies of each population, but this convergence is observed in all contexts. Notably, neither the total load nor total information gain depend on the strength of selection, which varies from one group of types to another. Mismatch load is expected to exceed information gain at all times in this setting (Proposition 2), and this relationship is observed in all competitions.

Theory predicts that information gain is a lower bound on the load incurred by selection. Our experimental populations suffer the requisite minimum load necessary to encode that information, but no more. These experimental results suggest that this relationship between information gain and load, suitably generalized, is not simply an artifact of abstractions in population genetics theory, but rather a general property of selection acting on real populations.

### The cost of selection as a learning process

While, substitution load was originally conceived of and studied as a “cost of selection” (Haldane 1937, 1957, Crow 1958, 1970, Kimura 1961, 1968), many have since argued that it is counterintuitive to view selection acting on beneficial variation as costly, since populations that evolve are surely better off than those that do not (Van Valen 1963, Brues 1964, Maynard Smith 1968, Sved 1968, O’Donald 1969, Moran 1970). However, the “cost of selection” that Haldane (1957) and Kimura (1961) refer to is not the disadvantage of a population that undergoes selection relative to one that does *not* evolve, but rather the disadvantage of a population needing to evolve relative to one that is *already* optimal. As Felsenstein (1971) points out, “it is appropriate to speak of a cost of selection, since the cost comes from the fact that natural selection is less efficient than divine intervention.”

That said, it is true that the generalization of substitution load— mismatch load—does not quantify precisely this notion of cost in all cases. Fundamentally, mismatch load measures the loss of potential fitness due to ongoing mismatch between types and environmental conditions. When there is only one environmental state (as in Haldane and Kimura’s substitution models) or when the same type is optimal in all conditions (as in our experiments), an optimal population is monomorphic for the most fit type and experiences no mismatch or load. In these cases, all mismatch is transient and attributable to the process of selection being incomplete, and mismatch load quantifies the cost of selection’s gradual nature as desired.

However, when the environment includes multiple conditions that favor different types, a type distribution that maximizes expected fitness over all conditions may still experience some load due to mismatch resulting from stochastic associations of types and conditions (i.e., mismatch load cannot be reduced to zero). In this case, not all of the cumulative load is attributable to the gradual nature of selection, and mismatch load does not accurately reflect the cost of the process itself. Instead, it would be more appropriate to quantify the cost of selection as the amount of *excess* load that accrues during evolution relative to the baseline load that is expected of an adapted population.

We do this below by extending and reinterpreting the cost of selection through the lens of learning theory.

#### A learning theoretic model of evolution

Analyzing the relative cost of learning processes is the purview of computational learning theory, which has seen recent adoption for the study of evolution (Valiant 2009, Chastain et al. 2014, Kaznatcheev 2020). Here we cast evolution by natural selection in a learning theoretic framework in order to clarify the meaning of mismatch load and develop a rigorous measure for the cost of selection as a learning process.

The general learning problem faced by an evolving population can be modeled as repeated play of a game between two players: the population and the environment (Figure 4, Appendix C.2). A play of this **population versus environment** (PvE) **game** in round *t* (i.e., generation *t*) consists of the population ‘choosing’ a distribution of types and the environment ‘choosing’ a distribution of environmental conditions to be experienced by the population. The distribution of types ***p***^*t*^ and the distribution of conditions ***x***^*t*^ represent the strategies of the population player and the environment player, respectively, in round *t* of the game. A particular learning problem instance can be defined by specifying an *n*-by-*m* game matrix **G** that gives the log fitness (i.e., growth rate) for each of the *n* types in each of the *m* environmental conditions (i.e., *G*_*ij*_= log*W*_*ij*_) and by specifying the process by which the environment updates its strategy *x*^*t*^ over time (Figure 4a). A wide range of biological contexts can be modeled with appropriate choices of **G** and ***x***^*t*^.

**Fig. 4.**
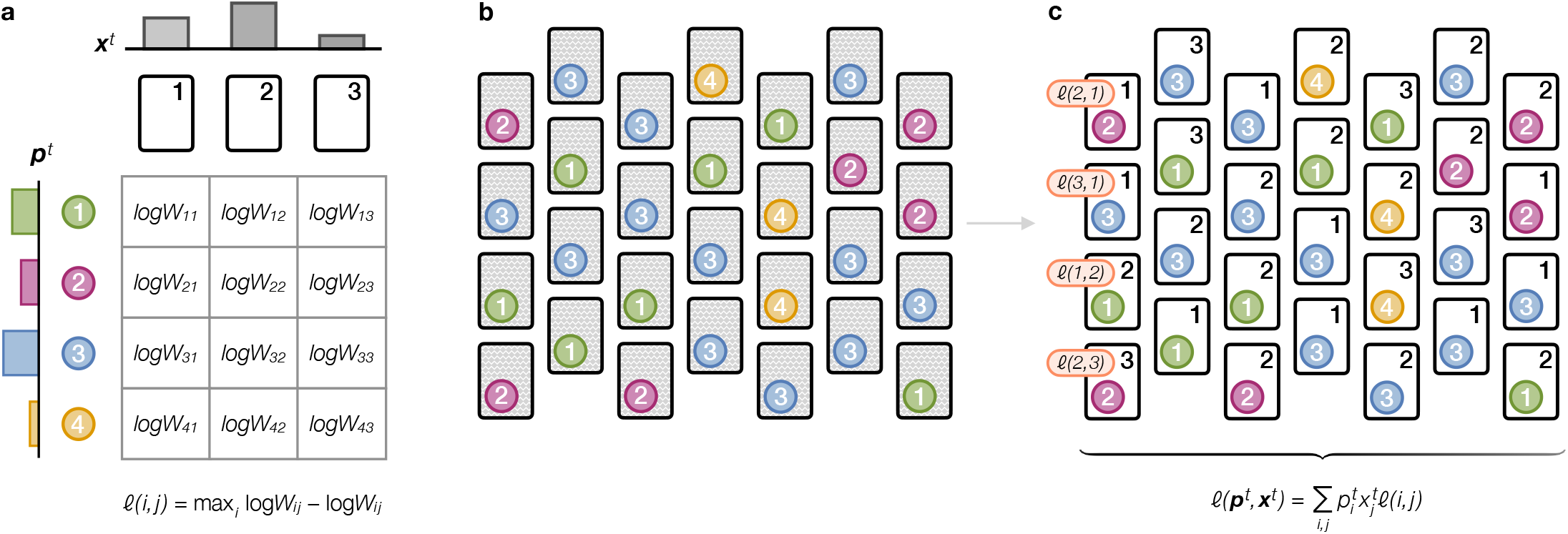
The Population versus Environment game. The learning problem faced by an evolving population can be modeled as a repeated game against the environment, which is illustrated here using the metaphor of a tabletop game. **(a)** The game is defined by the set of types available to the population player (tokens, rows), the set of conditions available to the environment player (cards, columns), and a game matrix G that specifies the payoff that each type receives when played against each environmental condition. Here the payoffs received by individual tokens are defined as the log fitness (growth rate) log*W*_*ij*_ of each type *i* in each condition *j*. **(b)** In each round *t*, the environment player puts out an array of environmental conditions (cards) with condition frequencies given by the environmental strategy *x*^*t*^. The population player selects a collection of types to play against the environment according to their type frequency strategy ***p***^*t*^. Each individual type is assigned to an independent environmental condition (one token placed on each condition card), but the pairing of types and conditions is random and out of both players’ control (condition cards are shuffled and remain face down until the type tokens are arbitrarily placed). **(c)** The environmental conditions that individuals have been paired with are then revealed. An individual with type *i* that encountered condition *j* receives a *loss l*(*i, j*) = max_*i*_*G*_*ij*_− *G*_*ij*_= max_*i*_ log*W*_*ij*_− log*W*_*ij*_ (a subset of individual loss terms are denoted in orange). The population player’s score in round *t* is the expected (mean) loss *ℓ*(***p***^*t*^, ***x***^*t*^) of all of the individuals (tokens) they played across the distribution of conditions (cards) played by the environment. The population player’s goal is to update their strategy ***p***^*t*^ so as to minimize their cumulative expected loss over many rounds of the game.

Each individual in the population experiences a micro-environment characterized by an independent condition drawn from the environment’s distribution of conditions *x*^*t*^ (Figure 4b). The association of particular conditions to specific individuals is assumed to be random and out of both players’ control. The loss of potential long-term fitness *ℓ*(*i, j*) that an individual of type *i* incurs in condition *j* is defined as the difference in log fitness (i.e., growth rate) between the optimal type for the *j*th condition and the *i*th type in the same condition

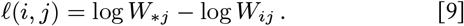

After each round *t*, the population player ‘observes’ the expected loss of each *i*th type across the distribution of conditions *x*^*t*^ (Figure 4c)

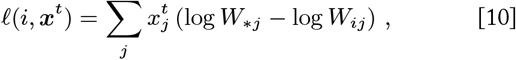

which provides evidence about the relative suitability of each type for the current environment. The population player can then use this type loss information to update the type frequencies that define its strategy ***p***^*t*^ according to some learning process.

The population incurs the expected fitness loss of its strategy in each round *t* (Figure 4c)

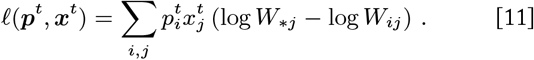

The **cumulative loss** of potential fitness accrued after *T* rounds of the population’s learning process is equivalent to the mismatch load *L*^*T*^ (Equation 8, Appendix D.1.2)

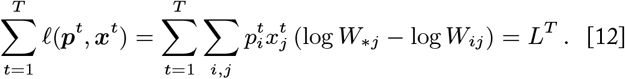

This connection reinforces the concept of mismatch load (and substitution load) as the cumulative loss of potential fitness due to mismatch between types and environmental conditions.

#### Regret measures the cost of learning

The ‘goal’ of the game for the population player is to minimize its regret about unfulfilled fitness when looking back after many generations. In learning theory, **regret** is defined as the difference between the cumulative loss that the population experiences as it updates its strategy ***p***^*t*^ over time and the loss it could have achieved had it played an optimal fixed strategy from the beginning. Regret is the true cost of a learning process: the excess loss one experiences in having to learn an effective strategy as opposed to knowing an optimal solution all along.

In general, the optimal strategy for the population depends on the particular sequence of environments it experiences and is determined in hindsight with knowledge of this sequence. There are multiple ways to define an optimal strategy with respect to a given environmental sequence. Here we consider a notion of optimality that follows from the dynamics of natural selection (another more general notion of optimality will be considered below). An evolutionarily stable state (ESS) 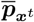 is a type distribution that locally maximizes expected fitness for a particular environment *x*^*t*^. In each update, selection always moves the population in the direction of an ESS for the environment at that time. If the ESS accessible to the population remains constant over a sequence of environments, then selection will continually approach and settle on this **stationary evolutionarily stable state** 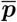 (i.e., 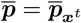 for all *t* ∈ [0, *T*]; Appendix C.3). A stationary ESS is the optimal strategy accessible to the population in the sense that it locally maximizes cumulative expected fitness over the corresponding sequence of environments. The ESS that the population approaches may not be globally optimal, but it is the best strategy that can be reached using replicator dynamics from the population’s initial state.

The population will accrue mismatch load as it gradually converges on the optimum. The population’s regret 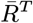 with respect to the stationary ESS 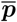 is the amount by which its load *L*^*T*^ exceeds the load 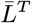 that it would have experienced had it used the optimal strategy 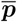 all along over the same sequence of environments.

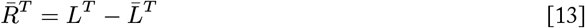

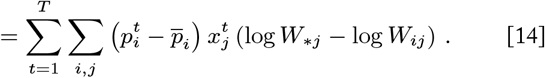

Here, the optimal strategy could be non-monomorphic and have a persistent load 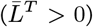 due to a stochastic environment. Contrast this with substitution load, which is defined in a setting where one type always has the highest fitness, and thus the optimal strategy that fixes that type incurs no load (i.e., 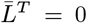 for all *T*). Therefore, Kimura’s substitution load is a special case of regret when there is just one universally optimal type.

For a biological population, regret is equal to the cumulative difference in expected growth between a hypothetical population using the optimal strategy 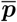 and the evolving population using ***p***^*t*^

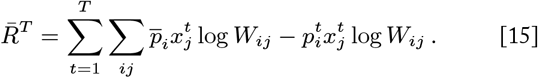

If we consider a lineage to be the collection of individuals derived from an initial population, this quantity can be equivalently expressed as the negative log ratio of the population’s total lineage size Γ^*T*^ to the size of the optimal lineage 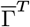 (Appendix D.2.2)

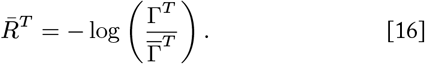

Just as relative fitness can refer to the short-term reproductive output of a type relative to the maximum output of other types, we can interpret the ratio 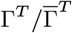 as the **relative lineage fitness**: the cumulative growth of the evolving lineage relative to the lineage that maximizes cumulative growth for the given sequence of environments. Therefore, the population’s ‘goal’ in the game can be interpreted as seeking to learn a type distribution that approximates the optimal strategy in order to maximize its relative lineage fitness and thus minimize its regret.

#### Natural selection as a no-regret learning process

In general, it would be intuitive for the population player to adapt their strategy by increasing the weight of types that have observed low fitness loss in the past while maintaining diversity to hedge against the future. Computer scientists and economists have shown that a simple, yet powerful, learning algorithm known as **Multiplicative Weights Updating** (MWU) is an effective solution along these lines for settings where the environment can change arbitrarily over time (Freund and Schapire 1999, Cesa-Bianchi and Lugosi 2006, Arora et al. 2012). Effectively, MWU balances concentrating weight on types that have performed well (i.e., incurred low loss) in the past against spreading weight over types that may perform well in the future (Appendix C.2.3). A learning rate parameter modulates the emphasis given to reacting to loss versus maintaining spread in each update. When the learning rate can be tuned in response to the learning problem, MWU is guaranteed to learn a strategy that converges on the performance of the optimal strategy in hindsight, which ensures that the learner’s per-round regret will approach zero in the long run (Freund and Schapire 1999, Cesa-Bianchi and Lugosi 2006).

Recent work has shown that replicator dynamics are equivalent to a fitness-based implementation of MWU with a constant learning rate that equally balances minimizing fitness loss (maximizing growth) and maintaining diversity (Chastain et al. 2014, Mehta et al. 2015, Meir and Parkes 2015, Chastain 2017) (Appendix C.2.4). Because the learning rate implicit in replicator dynamics is not tuned, the no-regret guarantees from the analysis of MWU do not automatically carry over to selection.

So far we have shown that information gain is a tight lower bound on regret for selection in the special case of an allele substitution (where substitution load is a special case of regret). Now we will characterize the cost of selection more broadly by establishing bounds on load and regret that hold in more general conditions.

When the sequence of environments is characterized by a stationary optimal strategy 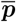, the total load for a population undergoing selection is bounded.

##### Proposition 3.

*(Appendix D.3.1)* *

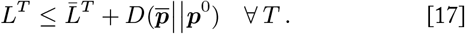

This bound guarantees that an evolving population’s load will be no greater than the load of the optimal strategy plus the divergence between the population’s initial type distribution ***p***^0^ and the optimal composition 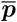. The divergence term represents how much learning the population has to do at the outset, which translates to excess load—regret—that accrues while the population is shifting its strategy toward the optimal. In fact, this divergence is an upper bound on regret with respect to a stationary ESS.

##### Theorem 1.

*(Appendix D.3.1) For any game matrix G and for any sequence of environmental conditions x*^0^, …, *x*^*T*^ *such that the population’s initial type distribution p*^0^ *is in the basin of attraction of an <u>evolutionary stable state</u>* 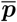 *that remains <u>stationary</u> for all t* ∈ [0, *T*], *the total regret* 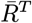 *with respect to* 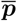 *of the trajectory of type distributions p*^0^, …, ***p***^*T*^ *generated by replicator dynamics is bounded from above by*

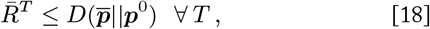

*with equality as* 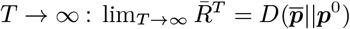.

In other words, the total cost of selection with respect to a particular ESS is equal to the amount of learning that the population must do to arrive there. The total regret is finite, which guarantees that the per-round regret of selection approaches zero in the long term (i.e., 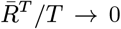 as *T* → ∞). Therefore, selection is a no-regret algorithm with respect to a fixed learning target, and selection is guaranteed to arrive at a type distribution that maximizes relative lineage fitness in this setting. This means that evolution by natural selection is an asymptotically optimal algorithm for solving the fundamental learning problem faced by evolving populations.

### The cost of information acquisition by natural selection

Populations that evolve by natural selection accumulate both information and fitness losses as a result of this learning process. Kimura noted a proportional relationship between information and load in the special case of an allele substitution (Proposition 1), and we observe that the accumulation of load outpaces that of information in such cases (Proposition 2, Figure 3). Having formalized and generalized the cost of selection in terms of regret, we can now investigate the cost of information acquisition by natural selection more generally.

The following result relates the information gained by selection in a single generation (as measured by the divergence *D*(***p***^*t*+1^||***p***^*t*^)) to the expected fitness loss due to type mismatch in the current and next generations.

#### Proposition 4.

*(Appendix D.4.1)*

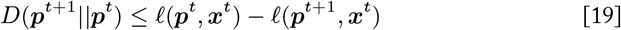

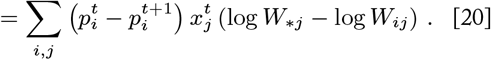

We can view the loss difference in Proposition 4 as the single-step regret that the population has about not already having the composition enjoyed by the next generation. This single-step regret is always greater than or equal to the information gained in that step. If the population were to have no regret relative to the next generation, then it would have nothing to gain—fitness nor information—in changing its composition. Thus the experience of regret must precede the acquisition of information.

The population’s cumulative information gain *I*^*T*^ after *T* generations is measured by the divergence *D*(***p***^*T*^ ||***p***^0^) of its evolved type distribution from its initial composition (Equation 2). The total amount of information that can be gained for a particular ESS 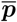 is given by the initial potential information 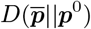. So long as the ESS remains stationary, the population will eventually converge on the ESS and acquire all of the potential information (i.e., 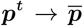 and 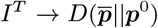 as *T* → ∞). We recognize the initial potential information as the same divergence that bounds the population’s regret with respect to a stationary ESS (Theorem 1). Therefore, we generalize Kimura’s finding that the cost of selection is proportional to the amount of information the population lacks out the outset:

#### Theorem 2.

*(Appendix D.4.2) For any game matrix G and for any sequence of environmental conditions x*^0^, …, *x*^*T*^ *such that the population’s initial type distribution p*^0^ *is in the basin of attraction of an <u>evolutionary stable state</u>* 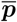 *that remains <u>stationary</u> for all t* ∈ [0, *T*], *the total information gain I*^*T*^ *and the total regret* 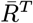 *with respect to* 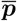 *of the trajectory of type distributions p*^0^, …, ***p***^*T*^ *generated by replicator dynamics both converge on the value of the initial potential information*

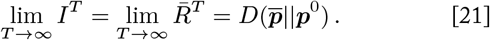

It follows from Theorem 2 that the cost of each bit of information is a two-fold reduction in relative lineage fitness in the long run. While a population that evolves by natural selection accumulates regret, all of the excess fitness loss it experiences is eventually translated into information encoded in its distribution of types. Selection is a highly efficient information acquisition process in this sense.

Given that regret is equal to subsitution load when the ESS 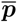 is monomorphic (such that 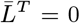 and 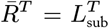), our experimental results (Figure 3) can be seen as validating the regret predictions given by Theorem 1 and Theorem 2 in that case. Even in our stochastically fluctuating real-world populations, natural selection converts all regret to information.

### Learning problems faced by evolving populations

The flexibility of this learning theoretic view of selection and its implications can be better understood by considering a few concrete cases.

#### Constant environments

First, let us revisit the simple case of a constant environment (i.e., *x*^*t*^ = *x*^*t*+1^ for all *t*). When the makeup of the environment does not change, a single type will be the best option in every round. Selection will continually approach fixation of this optimal type, which is a stationary ESS. We know from Theorem 1 that a population that evolves by natural selection will achieve vanishing per-round regret in learning this optimal strategy.

In a constant environment, achieving the goal of minimizing regret is equivalent to maximizing fitness in an absolute sense. Haldane and Kimura considered the simplest possible environment with only one unchanging condition (i.e., *m* = 1). In this case, selection increases the frequency of the type that has the highest fitness in this condition, which maximizes the mean fitness of the population as described by Fisher’s fundamental theorem (Fisher 1930). Similarly, if the environment consists of multiple conditions (i.e., *m >* 1) but the frequency of these conditions does not change, then selection will consistently favor the single type with the highest expected fitness over the constant distribution of conditions (Figure 5, left), which maximizes the geometric mean fitness in keeping with Gillespie’s geometric mean principle (Gillespie 1974).

**Fig. 5.**
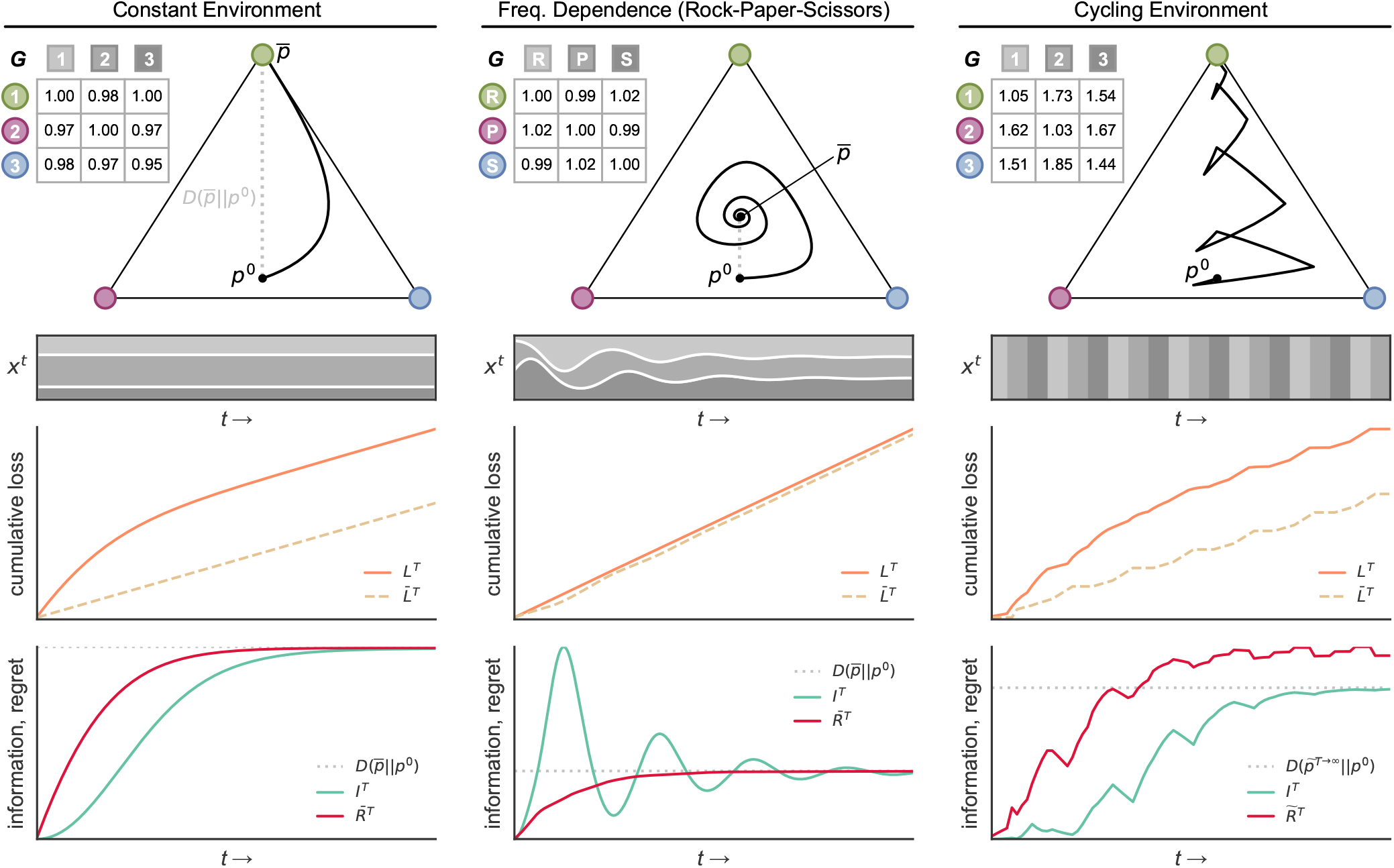
Load, regret, and information gain in different environmental contexts. Results from simulations of replicator dynamics in three different contexts (columns) are shown. In each column, the matrix **G** defines the log fitnesses (growth rates) for 3 types in each of 3 environmental conditions. Selection updates the type composition of the population along the black trajectory in the simplex. Gray dotted lines in the simplex depict the KL divergence 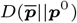 of a stationary ESS 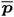 from the population’s initial state ***p***^0^, where applicable. Stacked frequency plots show the distribution of environmental conditions ***x***^*t*^ over time (i.e., a vertical slice at time *t* represents a distribution *x*^*t*^ over the environmental conditions shown in the game matrix). The mismatch load of the evolving population (*L*^*T*^, orange line) and of either the ESS composition (*L*^*T*^, dashed gold line, left and center columns) or the empirically optimal composition (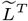, dashed gold line, right column) are plotted over time. The bottom-most plot in each column gives the information gain (teal line) and either the ESS regret (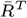, red line, left and center columns) or empirical regret (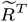, red line, right column) over the course of selection. **(Constant environment, left)** The environment is heterogeneous but the distribution of conditions is constant. Type 1 (green) has the highest expected fitness and increases in frequency in every generation. However, Type 1 incurs non-zero loss in Condition 2, so the optimal composition that is fixed for Type 1 accrues load over time (dashed gold line). Fixation of Type 1 is a stationary ESS, so regret is bounded by the initial potential information 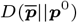 (Theorem 1). Information gain and regret converge on this value in the long-run (Theorem 2). **(Frequency-dependent selection, center)** Here the environmental conditions are defined by the types themselves, and the distribution of conditions is set to the type distribution in every generation (i.e., ***x***^*t*^ = ***p***^*t*^). This frequency dependence and choice of **G** define a “Rock-Papers-Scissors” scenario, where each type (‘R’, ‘P’, or ‘S’) is advantaged over one type and disadvantaged to the other type. Selection leads to a dampened oscillation in type frequencies that converges on a mixed stationary ESS. The existence of a stationary ESS implies that regret converges on the initial potential information 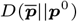 (Theorem 1). Information gain fluctuates before converging on the regret and initial potential information (Theorem 2). **(Cycling environment, right)** The environment consists of a single condition that switches cyclically over time. The ESS changes with each environmental shift, and selection moves the population in different directions in response to each condition. Type 1 has the highest expected fitness over the entire cycle and tends to increase in frequency. The empirically optimal strategy initially alternates in response to the environmental cycle before settling on the Type 1 fixation composition for the long-run. In this example, selection causes the population to converge toward the empirically optimal strategy, so the empirical regret is finite. The empirical regret exceeds the information gain at all times (Theorem 4).

#### Frequency dependence

Constant environments are only a small subset of what the learning theoretic view of evolution can express. In fact, there are no restrictions on the how the environment’s distribution of conditions can change over time. For example, the environment’s strategy in each round may depend on the population’s current or prior strategies, which allows for cases of frequency-dependent selection. For example, if the frequencies of conditions that individuals experience are set by the frequencies of types in the population (i.e., *m* = *n*, ***x***^*t*^ = ***p***^*t*^), then the make up of the “environment” is defined by the composition of the population itself, and fitnesses reflect interactions between types (the PvE game becomes an *effective evolutionary game* (Kaznatcheev 2017)). This context is of interest to evolutionary biology as it opens up new kinds of dynamics (e.g., social dilemmas) and is known to have a transformative effect on the adaptive capabilities of evolving populations (Kaznatcheev 2020).

The frequency-dependent setting reveals why the goal of the game is to maximize relative lineage fitness (i.e., minimize regret) rather than to maximize fitness outright. To illustrate this, consider a specific learning problem corresponding to the Prisoner’s Dilemma game described in Supplementary Figure D7 (note that this is a scenario where ***x***^*t*^ = ***p***^*t*^). In the Prisoner’s Dilemma the best outcome occurs if the population plays an all-cooperate strategy, which achieves the maximum possible population mean fitness. However, replicator dynamics lead toward a population of all defectors, which continually reduces the population’s mean fitness and clearly does not maximize fitness outright.

Recall that the only feedback the population player receives is the average fitness loss of each type in each round of the game. At each step, the population observes that no matter the environmental distribution the defector type always has lower expected loss (higher average fitness) than the cooperator. Based on this evidence, the defector type is seen to be better off in every round, and replicator dynamics increases its frequency accordingly. Arriving at an all-cooperate strategy would require the population player to use information that is not available to it (e.g., knowledge of **G** or knowledge of the environment’s process for updating *x*^*t*^) or to adopt a learning algorithm that is not generally robust across learning problems (e.g., up-weighting types with high relative loss). Thus when facing some environments, the population may be simply unable to maximize fitness in absolute terms by using a general online learning process.

In general, the attainable goal for the population is to maximize its cumulative fitness relative to the maximum possible fitness that could be achieved by a fixed strategy given the same sequence of environments. Evaluating the performance of a learning process with respect to the fixed sequence of environments in hindsight may seem strange, particularly when that sequence of environments was dependent on the trajectory of the population while the learning was underway. But reframing regret in terms of relative lineage fitness lends intuition to the fixed retrospective nature of this quantity.

For example, consider a large, well-mixed population of cells that evolves according to replicator dynamics in the Prisoner’s Dilemma scenario (Supplementary Figure D7). Suppose we identify two very small subpopulations: one that has the same initial type distribution as the overall population and one that is initially all defectors. In the first subpopulation, replicator dynamics increases the frequency of defectors and tracks the change in the overall population. The second subpopulation remains fixed for defectors, which turns out to be optimal. Given that these subpopulations are small relative to the overall population, their trajectories do not affect the makeup of the overall population that constitutes the “environment.” The two subpopulations therefore experience the same sequence of environments, and it is natural to ask how the cumulative growth of these two lineages compare given the conditions that play out. Relative lineage fitness quantifies this comparison. The more effective the learning process used by the first subpopulation, the greater its relative lineage fitness and the less regret it experiences. Just as the growth of a learning subpopulation can be compared to a fixed subpopulation in the same physical environment, the cost of a learning process is measured with respect to the fixed optimal strategy in hindsight, whether or not such an optimal population actually existed.

#### Arbitrary environmental change

In this section, we extend the concept of regret to the fully general setting where no assumptions are made about the environment. Regret measures the cumulative fitness loss of an evolving population with respect to a fixed optimal strategy for the same sequence of environments. So far, we have considered the fixed optimal strategy to be an ESS that is stationary over the observed sequence of environments. However, when the distribution of environmental conditions ***x***^*t*^ is free to fluctuate arbitrarily, shifts in the environment may cause the evolutionarily stable states to change. Whereas an environmental sequence with a stationary ESS represents a basic learning problem with a fixed solution, environments where the stable state switches from one interval to the next can be seen as a series of problems over which the population must integrate its learning.

If we make no restrictions on the sequence of environments, then we cannot determine a fixed optimal strategy based on the process that generates the environmental sequence or on the dynamical trajectory of the population, since these are free to change at any time. Instead, we consider the **empirically optimal strategy**:

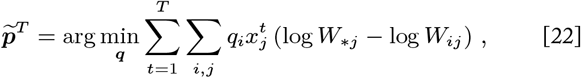

which is the fixed type distribution that would have minimized cumulative loss (Equation 12) for the particular observed sequence of environments in hindsight after *T* rounds.

In the learning problems that we have considered so far (i.e., constant environments and the Prisoner’s Dilemma) the empirically optimal strategy is always a stationary ESS as well (i.e., 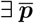 and 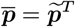 ∀*T*), but this is not always the case. For example, in the Snowdrift game scenario shown in Supplementary Figure D7 replicator dynamics carry the population to a polymorphic ESS, while the empirically optimal strategy is to play all-defect, which globally minimizes loss (maximizes fitness) for the given sequence of environments.

We refer to regret measured with respect to the empirically optimal strategy 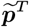 as **empirical regret**:

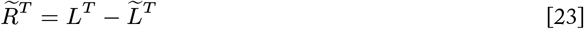

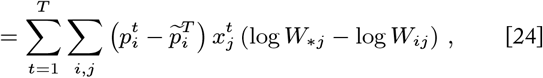

where 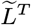 is the cumulative loss that would have been achieved using 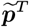 all along in the observed sequence of environments.

In general, the environment may fluctuate in such a way that the evolutionarily stable states change as time goes on. For example, in the Rock-Paper-Scissors and cyclical environment games shown in Figure 5, the strategy that is empirically optimal in hindsight changes in response to environmental shifts as the game progresses. The accumulation of excess load while learning strategies that turn out to be suboptimal later on increases the cost of the overall learning process. Selection is an asymptotically no-regret algorithm with respect to fixed learning targets, but the regret bound from Theorem 1 does not apply when the ESS accessible to the population is not stationary.

Nevertheless, results from analysis of the MWU algorithm allow us to bound the cumulative loss—mismatch load—for selection even in this general setting.

##### Proposition 5.

*(Freund and Schapire 1999)*

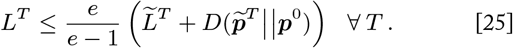

This bound is similar to the bound on load in the setting of a stationary ESS (Proposition 3), but is worse by a constant factor (where *e* denotes Euler’s number) that represents the additional cost of the population being caught out and having to change direction due to changes in the learning target. Even still, this bound guarantees that the mismatch load for an evolving population in any environment will not be much more than the minimum possible load 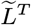 in the long run.

This result leads to a corresponding upper bound on the population’s empirical regret in any environment.

##### Theorem 3.

*(Appendix D.3.2) For any game matrix* ***G*** *and for any sequence of environmental conditions* ***x***^**0**^, …, ***x***^*T*^, *the total empirical regret* 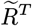 *with respect to* 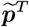 *of the trajectory of type distributions* ***p***^**0**^, …, ***p*** *generated by replicator dynamics is bounded from above at all times by*

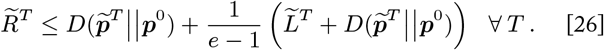

In the fully general case, the population’s regret is not bounded by a finite value and may continue to increase over time. The gradual nature of selection causes the population to accrue regret whenever it must move to a new ESS. If environmental fluctuations cause the ESS to change such that that the population cannot settle into an optimum for the long haul, then the population will continue to accumulate regret as it adapts to each new learning target. This general regret bound makes no assumptions whatsoever about the types, conditions, fitnesses, or sequence of environments that constitute the learning problem. Therefore, this bound provides an upper limit on the total empirical regret a population can possibly experience.

When the environment is variable, the population’s information gain may rise and fall over time as it adapts to changing conditions. We can interpret these fluctuations as the population gaining information about the present environment and “unlearning” information it had acquired about spurious trends in the past. For example, consider the Rock-Paper-Scissors scenario shown in Figure 5 (center). In response to conditions that initially favor the S type, the population “overlearns” a S-dominant strategy, and the information gain exceeds the initial potential information as the population overshoots the ESS composition that is optimal for the long term. Ongoing shifts in the frequency-dependent environment lead to oscillations in information gain as the population cyclically learns and unlearns strategies dominated by each type in turn. Eventually the population learns a balanced optimal strategy as it settles into the stationary ESS, and the total information gain converges on the initial potential information.

Despite possible fluctuations in information gain, selection continually makes progress in learning about the current ESS 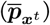. That is, the potential information 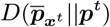 that remains from the population’s current state ***p***^*t*^ decreases monotonically over intervals where the ESS 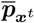 remains constant (Appendix B.2.2).

In all cases, we find that the information gain is bounded by the empirical regret at all times

##### Theorem 4.

*(Appendix D.4.3) For any game matrix* ***G*** *and for any sequence of environmental conditions* ***x***^**0**^, …, ***x***^*T*^, *the total information gain I*^*T* +1^ *of the trajectory of type distributions* ***p***^0^, …, ***p***^*T* +1^ *generated by replicator dynamics is bounded at all times by the total empirical regret* 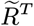 *with respect to the empirically optimal strategy* 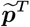

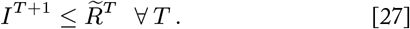

For a biological population that evolves by natural selection, every bit of information about the environment requires at least a commensurate reduction in lineage fitness relative to the empirically optimal strategy. This result holds for any evolutionary learning problem (i.e., any choice of **G** and sequence of ***x***^*t*^) and confirms that there is a minimum fitness cost of information for natural selection in all contexts (note that our Proposition 2 is a special case of Theorem 4 when 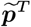 is monomorphic such that 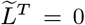 and thus 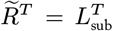 Appendix D.4.3).

In general, when the environment changes arbitrarily the population pays a higher price for information. We have seen that the upper bound on regret is higher in environments for which the optimal strategy in hindsight changes over time due to additional load associated with the population learning evolutionarily stable states that are not ultimately optimal in the long run (Theorem 3). In environments that fluctuate with a pattern that can be learned, selection will eventually settle on a strategy that approximates the empirically optimal strategy and achieves vanishing per-round regret (e.g., Figure 5, right). In such a case, the cost per bit converges on a finite value in the long run. When the environment varies unpredictably, there may be no stable empirically optimal strategy, and regret and the cost per bit may not converge. In any case, the maximum regret cost of information for selection is bounded.

### Comparing natural selection to other learning algorithms

Replicator dynamics is not the only conceivable algorithm that the population player could have used to update its strategy. Any algorithm that adapts the population’s strategy by increasing the weight of types that have observed low fitness losses in the past would be reasonable. The learning algorithm known as Follow the Leader (FTL) is an extreme implementation of this heuristic, which places all of the weight in each round on the single type that has received the lowest cumulative loss against the previous conditions of the environment (Appendix C.2.3). When the same type always has the highest fitness, an FTL learner will fix the optimal type after a single round, and its only regret will be the excess loss observed for the initial type distribution. Replicator dynamics will not be as fast, but selection will still converge on the optimal type. In such a case, a population that learns using selection will have lower relative lineage fitness and more regret than one using FTL, but this “inefficiency” is bounded, and all regret is converted to information (Theorem 1, Theorem 2).

Learning processes such as FTL that react *more* strongly than selection to each round of observed losses can achieve lower regret than selection in simple environments, but environments that are highly variable or unpredictable can drive such reactive learning processes to high regret. On the other hand, learning processes that are *less* responsive than selection can experience high regret in simple environments because they approach optimal strategies even more gradually. The balance that selection strikes in minimizing loss by responding to previous observations while maintaining diversity to hedge against the future makes it generally robust to arbitrary environmental change.

Theorem 3 provides a benchmark for comparing the worst-case performance of selection to other learning algorithms, as is standard practice in computer science. When the environment is variable such that the optimal strategy has non-trivial load, the worst-case regret (and thus the maximum cost of information) of selection is lower than that of other learning processes that are substantially more or less reactive. (Appendix D.3.3, Figure D6). Further delineating the classes of environments where selection typically outperforms other learning processes is an area for future research.

## Discussion

Our results, summarized in Table 1, highlight the intimate relationship between fitness and information. When a population has types with suboptimal fitnesses for the current conditions, natural selection incorporates this information about the environment by gradually adapting its composition. If the population does not suffer suboptimal fitnesses then there is nothing it needs to learn. Naturally, then, the accumulation of fitness losses must precede and outpace the acquisition of information, but selection effectively converts observed losses to information gain. As a population gains information it arrives at a more adaptive matching of types to environmental conditions and achieves higher relative lineage fitness than a population without this information. Potential information not yet held is potential growth forsaken.

**Table 1.** Summary of results relating regret and information gain under increasingly general conditions. The arrow notation 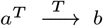 signifies convergence (lim_*T*→∞_*a*^*T*^ = *b*), and the angled arrow notation 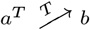 signifies convergence from below (lim_*T*→∞_ *a*^*T*^ = *b* and *a*^*T*^ ≤ *b* ∀ *T*).

| Environment | Homogeneous ($m = 1$) | Heterogeneous ($m > 1$) | | |
| --- | --- | --- | --- | --- |
| | Constant ($\mathbf{x}^t = \mathbf{x}^t \forall t$) | Constant | Time-Varying | |
| | Stationary ESS ($\bar{\mathbf{p}} = \mathbf{p}_{\mathbf{x}^t} \forall t$) | Stationary ESS | Stationary ESS | Arbitrary |
| Cost of information | $I^T \nearrow L_{\text{sub}}^T$<br>Kimura (1961),<br>Proposition 2 | $I^T \xrightarrow{T} \bar{R}^T$<br>Theorem 2 | $I^T \xrightarrow{T} \bar{R}^T$<br>Theorem 2 | $I^T \leq \tilde{R}^{T-1} \forall T$<br>Theorem 4 |
| Regret bound | $L_{\text{sub}}^T \nearrow -\log p_{\mathbf{x}^t}^0$<br>Kimura (1961),<br>Proposition 1 | $\bar{R}^T \nearrow -\log p_{\mathbf{x}^t}^0$<br>Theorem 1 | $\bar{R}^T \nearrow D(\bar{\mathbf{p}} \parallel \mathbf{p}^0)$<br>Theorem 1 | $\tilde{R}^T \leq D(\tilde{\mathbf{p}}^T \parallel \mathbf{p}^0) + \frac{1}{e-1} (\bar{L}^T + D(\tilde{\mathbf{p}}^T \parallel \mathbf{p}^0)) \forall T$<br>Theorem 3 |

A learning theoretic view of evolution establishes regret, a measure of relative load, as the appropriate definition of the cost of selection. Populations that evolve by selection reliably learn a strategy that maximizes relative lineage fitness and achieves vanishing per-generation regret relative to a fixed learning targets (i.e., a stationary ESS). In general, selection balances load minimization and diversity maximization, which enables the process to fare well in complex environments. These results generalize the concept of substitution load and breathe new intuition into the concept of “the cost of selection.” In addition, we revive, formalize, empirically validate, and extend Kimura’s initial insight that there is a fundamental fitness cost for every bit of information that is acquired by selection. That said, our theory and experiments demonstrate that natural selection is a highly effective learning process that converts all regret to information with respect to particular learning problems. In addition, we show that the price of information is better for selection than for some alternative learning processes in the worst case.

We have focused on natural selection as a process that acquires adaptive information about standing variation. However, selection interacts with other forces in biological evolution. Notably, mutation introduces new types and expands the set of compositions that selection can learn about. Together mutation and selection move populations across adaptive landscapes, but finding optima in complex landscapes is a computationally hard problem (Kaznatcheev 2019). Reconciling the effectiveness of selection as a learning process with the complexity of evolutionary search is important to understanding the performance of Darwinian evolution in an algorithmic sense. In addition, determining the extent to which recombination impacts the rate of information acquisition, how this depends on the fitness landscape, and whether any such effects confer adaptive advantages at the level of the individual is another interesting direction for future research. We anticipate that continued integration of evolutionary theory, information theory, and learning theory will lead us to a richer understanding of adaptive evolution.

## Materials and Methods

### Bacterial strains

*Escherichia coli* B (REL606) strains were used in selection experiments and related assays. A “wild type” strain (WT) and three strains with unique mutations in the RNA polymerase *rpoB* gene (M1, M2, M3) were obtained with permission from the −80^°^C strain archive from Lindsey et al. (2013). Mutations to the *rpoB* gene conferred each mutant strain with a distinct exponential growth rate that was reduced from that of the WT strain (Supplemental Figure E1). As the strain with the optimal growth rate, the WT strain was transformed with a plasmid engineered to carry the green fluorescent protein (GFP) gene *mGFPmut2*. The *rpoB* mutant strains were each transformed with plasmids engineered to carry the red fluorescent protein (RFP) gene *mScarlet-I* A full description of all strains and plasmids used in this study can be found in Appendix E.1.

### Selection experiments

Strains were cultured in standard Luria-Bertani (LB) broth with 15*μ*g/mL tetracycline for marker plasmid retention (hereafter referred to as “media”). Selection competition experiments were conducted between three pairs of strains: WT vs. M1, WT vs. M2, and WT vs. M3. Each pair of strains participated in three competitions with different initial strain frequency compositions: WT at approximately 12.5%, 25%, and 50% of the initial population, respectively. Selection experiments were initiated by combining the strains to be competed in the designated ratios at a density of ∼1 × 10^6^ cfus/mL in 25 mL of fresh media. Every 4 hours, a 1 mL sample was taken from each competition culture to measure strain densities and frequencies using flow cytometry (see Appendix E.2.2 for more information). At the same time, a sample of the culture was transferred to 25 mL of fresh media using an adaptive transfer protocol that ensured that the culture would remain in exponential growth while maintaining a measurable density for flow cytometry (see Appendix E.2.1 for more information). Competitions proceeded in this fashion for 36 hours. See Appendix E.2: Supplemental Methods for more information.

## Acknowledgments

We thank Michael Lachmann, Joe Felsenstein, Kevin Gross, Frazer Meacham, Emily Brunelli, and the Kerr Lab for valuable discussions. We also thank Katie Dickinson for exceptional guidance and assistance with experimental techniques and logistics, and Reilly Falter and Alena Beloborodyy for their contributions at the bench. RSM and OK were supported by the NSF Graduate Research Fellowship Program and by the BEACON Center for the Study of Evolution in Action. AK was supported by a fellowship from the James S. McDonnell Foundation.

## Appendices

*These appendices aim to provide comprehensive background, derivation, and interpretation for all of the key concepts and results discussed in this work. It is our hope that any interested reader, regardless of background, can achieve an understanding of these topics by working through this material without having to rely on other sources. This work draws from several rich areas of theory, but necessary concepts from information theory, game theory, and learning theory are introduced throughout. We take care to derive definitions and results step-by-step and with interpretation, and most results require only basic mathematical tools. Appendices A, B, and C review relevant concepts and results from the literature and establish the formalisms used in this work*. ***Readers interested only in proofs of key results from this work can turn to Appendix D. Supplemental methods and results from our selection experiments can be found in Appendix E***.

### Appendix A: Modeling Natural Selection

#### A.1 The replicator dynamics model of natural selection

Replicator dynamics are a standard model of evolution that describe the action of natural selection in large populations of replicators without mutation, recombination, or other sources of new variation. Replicators are entities that generate copies of themselves and may represent alleles, genotypes, strategies, beliefs, etc. Populations may consist of multiple types of replicators, each with a characteristic fitness that gives their rate of reproduction. Replicator types with above-average fitness will increase their frequency in the population, and types with below-average fitness will decrease in frequency. Replicator dynamics are widely studied in population genetics, ecology, evolutionary game theory, learning theory, economics, and other contexts.

##### A.1.1 Discrete-time replicator dynamics (discrete generations)

###### Fitness and growth

First we consider a population of replicators with a set of *n* alternative types that reproduce in discrete generations. Let 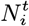 be the number of individuals of the *i*th type in generation *t*. The expected number of offspring left by each individual of type *i* in a generation is given by *W*_*i*_, which is referred to as the **Wrightian fitness** of the *i*th type (In general, the fitness of a type may depend on the current environmental conditions, which we will develop further in Appendix A.2). The number of individuals of type *i* after a total of *T* generations is given by

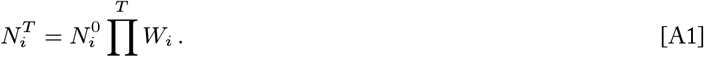

where 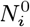 is the initial number of individuals of type *i*. The expected number of descendants per initial individual after *T* generations is given by the ratio 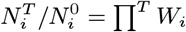.

Suppose that we observe 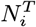 individuals of type *i* after *T* generations, and we wish to know the effective growth rate of this subpopulation over this interval. In other words, what average growth rate 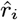 corresponds to the reproductive output of individuals of this type? Solving the following expression for 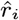, we have

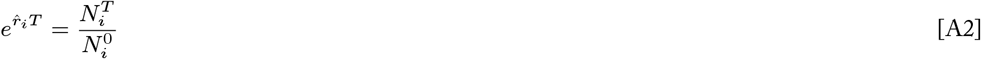

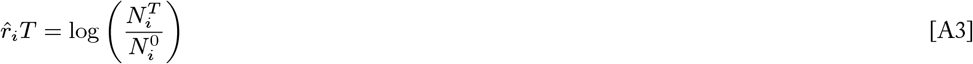

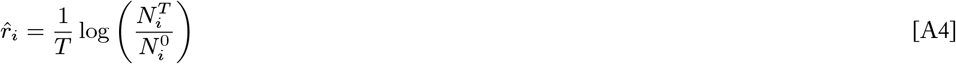

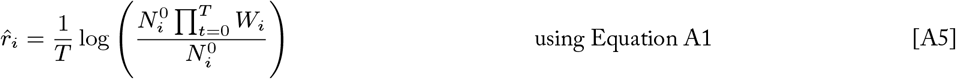

using Equation A1

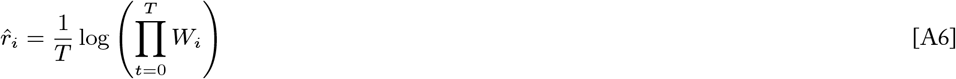

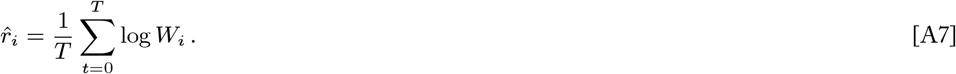

The effective growth rate of the *i*th type over many generations is equal to the expected logarithm of the type’s fitness, and we refer to the **log fitness** log*W*_*i*_ as the effective growth rate of type *i* in a single generation (i.e., 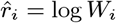 for *T* = 1).

The **relative fitness** of the *ith* type, *w*_*i*_, is considered with respect to the highest fitness of any type in the population, *W*_∗_

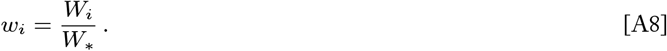

By this definition, the relative fitness of the optimal type is equal to 1 (i.e., *w*_∗_ = *W*_∗_*/W*_∗_ = 1), and the relative fitnesses of all other types take values in [0, 1]. As such, we may also choose to represent the relative fitness of the *i*th type as

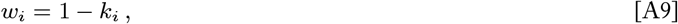

where *k*_*i*_ ∈ [0, 1] is the **Wrightian selection coefficient** of the *i*th type, and *k*_∗_ = 0 is the selection coefficient of the optimal type. This expression of relative fitness can be interpreted as saying that for every 1 offspring left by the optimal type there are 1 − *k*_*i*_ offspring left by the *i*th type.

###### Dynamics of type frequency change

The frequency of type *i* in the population in generation *t* is defined as

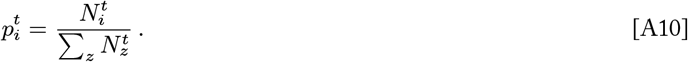

Then the change in the frequency of type *i* from one generation to the next is given by

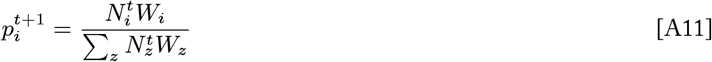

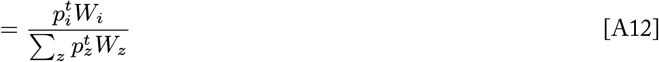

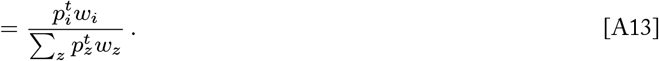

**Discrete-time replicator dynamics** refer to this dynamical map

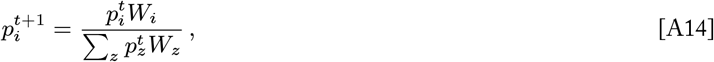

where 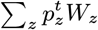 is the mean fitness of the population at time *t*. This dynamic re-weights the frequencies of types proportional to their fitnesses such that types with above average fitness increase in frequency and those with below average fitness decrease in frequency.

##### A.1.2 Continuous-time replicator dynamics (overlapping generations)

###### Fitness and growth

While not the focus of the main text, we can also consider the evolution of populations with overlapping generations. Rather than adding individuals to the population at discrete unit intervals, individuals are added to the population at multiple instances per unit time. To capture this, we can express the expected reproductive output per unit time as

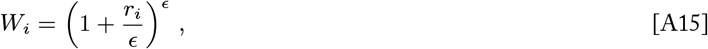

where *r*_*i*_ gives the expected number of offspring above replacement (or below, if negative) that each *i* individual contributes to the population per unit time, and *ϵ* gives the number of instances that offspring are added per unit time. When *ϵ* = 1, then this definition reduces to the discrete generation case, where *W*_*i*_= 1 + *r*_*i*_ defines the absolute Wrightian fitness of the *i*th type. On the other hand, fully continuous growth is represented by allowing new individuals to be added to the population at an infinite number of points per unit time, which results in the familiar exponential growth term

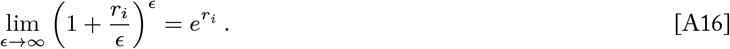

Then the number of *i* individuals at time *t* in a continuously growing population is given by

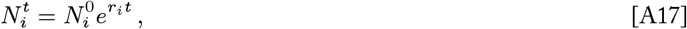

which is known as the Malthusian growth model. Here we see that the parameter *r*_*i*_ sets the exponential growth rate of the *i*th type, and we refer to this rate *r*_*i*_ as the **Malthusian fitness** of the *i*th type. The **Malthusian selection coefficient** *s*_*i*_ of the *i*th type is defined as the difference between the Malthusian fitness (i.e., growth rate) of the optimal type and that of the *i*th type

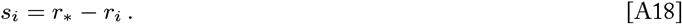

###### Dynamics of type frequency change

In this section, we consider the continuous analog of the discrete replicator dynamics presented above. The frequency of the *i*th type at time *t*, 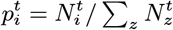, is defined as it was for the discrete-time model. We find an expression for the change in this frequency by taking the derivative

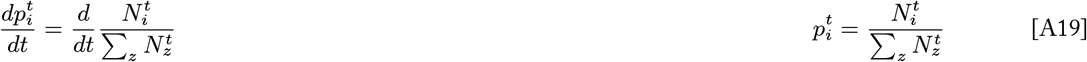

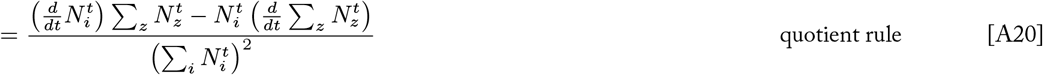

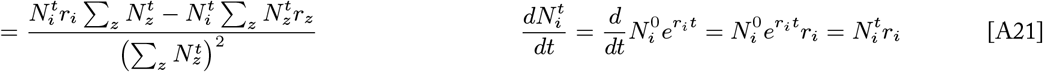

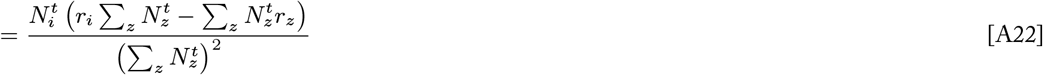

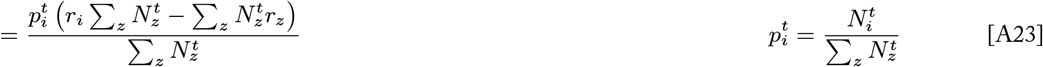

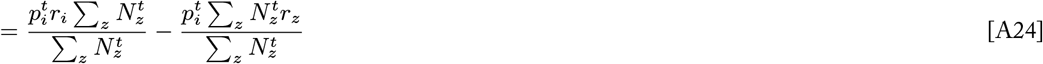

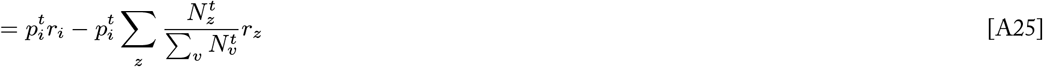

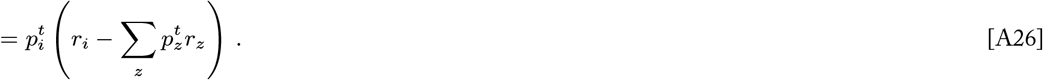

Thus, the **continuous-time replicator dynamics** are given by the system of differential equations

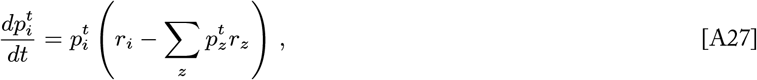

where 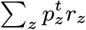 is the mean Malthusian fitness (i.e., growth rate) of the population at time *t*. Like the discrete-time dynamics, this dynamic re-weights the frequencies of types according to their fitnesses relative to the population mean fitness.

#### A.2 Modeling variable environments

In general, the fitness of a individual depends on both its type and its environment, and we are ultimately interested in modeling the process of natural selection in contexts where conditions and fitnesses change over time. In this section we extend the standard replicator dynamics models to support both heterogeneous and time-varying environments, broadly defined.

Suppose the population occupies an environment that is comprised of a set of *m* distinct environmental conditions. We assume that every individual in the population experiences an independent environmental condition, as if each individual occupies a separate part (i.e., micro-environment) of the physical environment. The association of specific individuals to particular conditions is assumed to be random, and the probability that a given individual experiences the *j*th condition at time *t* is given by 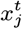. The condition distribution ***x***^*t*^ defines the make up of the environment at time *t*, and scenarios of environmental change can be modeled by specifying a particulars sequence of environments {***x***^0^, ***x***^1^, …, ***x***^*T*^ }.

We take a very expansive view of what constitutes an “environment.” Essentially any set of contextual factors that have a conditional impact on the fitnesses of types can be considered an environmental condition. For example, conditions may represent the abiotic conditions experienced by individuals, the types of other replicators encountered by individuals, or even the identity of other alleles carried by individuals. Similarly, the sequence of environments {***x***^0^, ***x***^1^, …, ***x***^*T*^ } may be generated according to any process, including as a function of time, as a function of type frequencies, as a stochastic or random process, as a “pre-programmed” sequence, etc. Throughout this work we consider various environmental contexts of interest as well as the fully general setting where no assumptions are made about the sequence of environmental condition distributions.

In the main text, we frame this model of discrete-time replicator dynamics in variable environments in terms of repeated play of a two-player, population versus environment game, which lends itself to learning theoretic analysis; see Appendix C.2.2 for more information about this formalization.

##### Discrete-time replicator dynamics in variable environments

Let *W*_*ij*_ denote the Wrightian fitness of an individual of type *i* in environmental condition *j*. The average fitness of type *i* across the distribution of environmental conditions ***x***^*t*^ is denoted 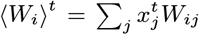, and the average fitness the population across all types and conditions is denoted 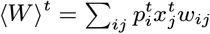. Then the discrete-time replicator dynamics in a heterogeneous environment becomes

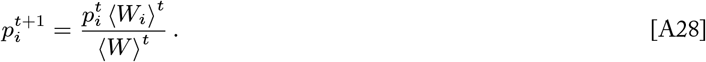

##### Continuous-time replicator dynamics in variable environments

Let *r*_*ij*_ denote the Malthusian fitness (i.e., growth rate) of an individual of type *i* in environmental condition *j*. The average growth rate of type *i* across the distribution of environmental conditions ***x***^*t*^ is denoted 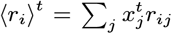, and the average growth rate of heterogeneous environment becomes the population across all types and conditions is denoted 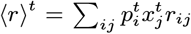. Then the continuous-time replicator dynamics in a heterogeneous environment becomes

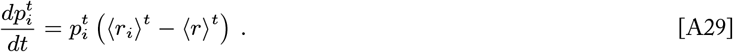

### Appendix B: Natural Selection as an Information Acquisition Process

#### Information Measures

Information theory uses probability theory and statistics to formalize the familiar notion that information is that which reduces uncertainty. The more an observation reduces one’s uncertainty about the state of some system, the more information that observation provides. Information-theoretic measures allow us to rigorously quantify both uncertainty and information in a system.

#### Entropy quantifies uncertainty

Consider a random variable *X* that can take some value *i* ∈ 𝒳 (e.g., the outcome of a die roll), and suppose you assign a probability *p*(*i*) to each outcome value based on your belief of how likely it is to occur. If you are fairly certain that one of the outcomes will occur, you will ascribe high probability to that outcome and low probabilities to all other outcomes. If you are more uncertain about which outcome will occur, your probability distribution *p* will be more uniform. This notion of uncertainty is quantified by the **entropy**:

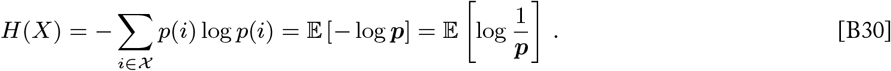

Entropy is a property of the spread of the distribution ***p*** and does not depend on the values of the random variable *X*. As such, the entropy of *p* may also be denoted *H*(*p*).

Entropy can be interpreted as the expected **surprisal** of an outcome, where the surprisal of the outcome *i* under the probability distribution *p* is defined as log (1*/p*(*i*)). If an outcome that was ascribed a high probability of occurring does occur, this gives low surprisal (log (1*/p*(*i*)) → 0 as *p*(*i*) → 1), but the occurrence of an outcome that was ascribed low probability gives high surprisal (log (1*/p*(*i*)) → ∞ as *p*(*i*) → 0). The more uncertain you are about the outcome — the more uniform your probability distribution — the more surprising observed outcomes are on average, and the higher the entropy.

The **joint entropy** *H*(*X, Y*) of two discrete random variables *X* and *Y* is the entropy of their joint probability distribution

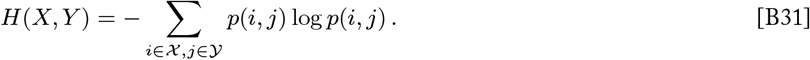

The **conditional entropy** *H*(*X*|*Y*) measures the expected amount of uncertainty about *X* that remains when the value of *Y* is known, averaged over possible values of *Y*

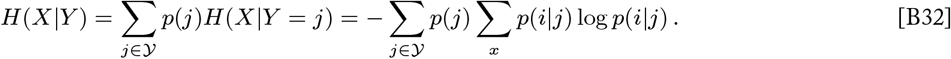

Reflecting the relationship between information and uncertainty, other information measures can be expressed as linear combinations of entropies.

#### Relative entropy (KL divergence) quantifies information gain

Consider a random variable *X* that is hypothesized to follow some probability distribution ***q***. Now suppose that you receive new evidence that leads you to update your hypothesis to some new probability distribution ***p***. How much information did this update provide? **Information gain** quantifies how much this new evidence reduces your uncertainty (i.e., expected surprisal) as the reduction in entropy under the posterior distribution ***p*** relative to the prior distribution ***q***

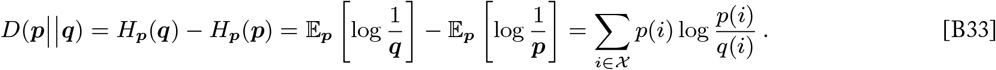

Here both expectations of surprisal (i.e., entropies) are calculated under the posterior distribution ***p***, which is the latest working hypothesis of the true event probabilities (Note: *H*_***p***_(***q***) = −Σ_*x*_ ***p***_*i*_ log ***q***_*i*_ is termed the *cross entropy* between ***p*** and ***q***). Information gain *D*(***p*** ∥ ***q***) is synonymous (i.e., equivalent) to the **relative entropy** or **Kullback-Leibler divergence** between distributions ***p*** and ***q***. This quantity can be interpreted as the excess uncertainty (surprisal) we suffer by holding the hypothesis ***q*** in a world where event probabilities actually follow distribution ***p***.

Information gain (i.e., relative entropy, KL divergence) is always non-negative and is zero if and only if ***p*** = ***q***. It is often useful to think of this measure as the “distance” between distributions, but the KL divergence is not a true distance measure as it is asymmetric (*D*(***p***∥***q***) ≠ *D*(***q*** ∥ ***p***)) and does not follow the triangle inequality.

#### Mutual information quantifies non-independence

Often we are interested in quantifying how informative an observation is about the state of a system. In other words, how much is our uncertainty about the state of a system *X* reduced by the state of an observation *Y* ? Of course, the observation *Y* can only carry information about the system *X* if the states of *X* and *Y* are somehow dependent on each other. The more dependent *X* and *Y* are on each other, the more knowledge of *Y* specifies the state of *X* (and vice versa).

The **mutual information** I(*X*; *Y*) of random variables *X* and *Y* quantifies precisely this notion of information as the reduction in entropy (i.e., uncertainty) of one random variable given observation of another random variable

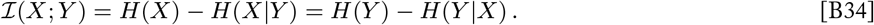

Mutual information can be equivalently expressed as the divergence from independence of two random variables. When *X* and *Y* are independent random variables, their joint distribution is equal to the product of their marginal distributions. Mutual information is equivalent to the relative entropy (KL divergence) between the actual joint distribution ***p***_*XY*_ and the hypothetical product distribution ***p***_*X*_ ***p***_*Y*_ that assumes independence.

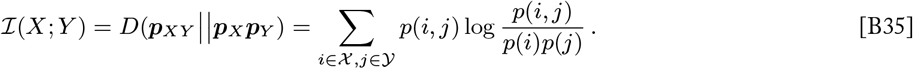

Mutual information can be interpreted in terms of relative entropy as the amount of information we gain by updating our posterior to reflect some dependence between *X* and *Y* from a prior that assumed independence.

#### Bits as units of information

When the base of the logarithms is 2, entropy and corresponding information measures are measured in *bits*. For an interpretation of the bit as a unit of measure, consider the following example (adapted from Cover and Thomas (2006)). Let *X* be a random variable that can take some value *i* ∈ {*a, b, c, d*} with probability distribution

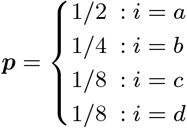

Suppose *X* takes a random value and we wish to determine this value with as few binary (“yes-no”) questions of the form, “*Is X* = … *?*” as possible. The most efficient approach is to ask the question whose two outcomes are closest to equiprobable at each stage. Doing so either identifies the value of *X* or reduces the space of possible outcomes by about half. In this example, one should start by asking, “*Is X* = *a*?” If yes, *X* has been determined with 1 question; if no, follow up by asking “*Is X* = *b*?”, and so on. *X* is a random variable, so the number of binary questions necessary to determine its value is also a random variable. The minimum expected number of binary questions to determine *X* in this example is

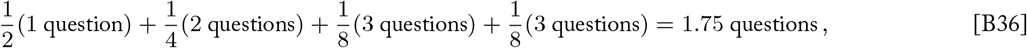

which is equal to the entropy of *X*

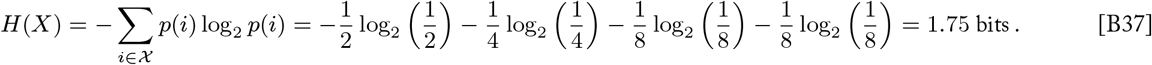

The minimum expected number of binary questions to determine *X* is equal to the entropy *H*(*X*) for dyadic probability distributions (i.e., where all probabilities are powers of 2—such as the one in this example) and lies between *H*(*X*) and *H*(*X*) + 1 in general.

Therefore, bits can be interpreted as the expected number of binary questions necessary to determine the value of a random variable. Bits of *entropy H*(***p***) represent uncertainty in terms of the average number of questions that must be asked to arrive at certainty about a random variable *X* that takes values according to ***p***. Bits of *relative entropy D*(***p***||***q***) represent a relative excess in uncertainty in terms of the expected number of extra questions that must be asked when choosing questions based on some distribution ***q*** instead of the true underlying distribution ***p***. Bits of *mutual information I*(*X*; *Y*) represent reduction in uncertainty in terms of how many questions about *X* are saved on average when given some other observation *Y*. In all cases, 1 bit refers to 1 binary question that must be asked on average to arrive at certainty. *In other words, 1 bit corresponds to 1 two-fold difference in uncertainty*.

#### B.2 Natural selection acquires information about the environment

##### B.2.1 Information gain

In the context of natural selection, the distribution of type frequencies ***p***^*t*^ can be seen as the population’s hypothesis about which types are most suited for the current environment. The relative reproductive success of each type provides evidence about the suitability of each type, and replicator dynamics updates the population’s hypothesis such that highly fit types are given more weight, and poorly fit types are given less. This process encodes information about the observed fitnesses of types into the population’s type composition. We can measure how much information is gained in these updates by measuring how much the population’s type frequency distribution changes during this process. The relative entropy (Kullback-Leibler divergence) between the population’s initial type distribution ***p***^0^ and its updated distribution ***p***^*T*^ quantifies the amount of **information gain** *I*^*T*^ for a period of selection of duration *T*

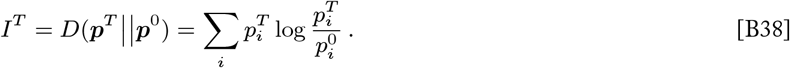

In Bayesian terms, *D*(***p***^*T*^ ∥ ***p***^0^) is a measure of the information gained by revising the population’s hypothesis from the prior distribution ***p***^0^ to the posterior distribution ***p***^*t*^.

##### B.2.2 Potential information

Relative entropy quantifies how much information is gained in moving from one probability distribution (e.g., a prior) to another (e.g., a posterior). We have defined the relative entropy *D*(***p***^*t*^ ∥ ***p***^0^) as the information that a population has gained by natural selection up to time *t* (Appendix B.2.1). Here we consider the relative entropy (KL divergence) of some fixed distribution 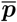 from the population’s current frequency distribution ***p***^*t*^:

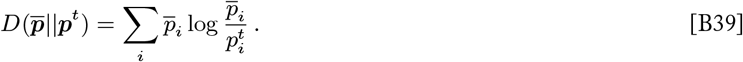

This relative entropy represents the amount of information that would be gained if the population’s type frequency distribution were to change from ***p***^*t*^ to 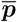. We can interpret this quantity as the **potential information** of the system: the information that is available to be gained in moving to 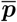 from the population’s current state.

###### Potential information is always decreasing with respect to evolutionarily stable states

We would like to determine the conditions where a population undergoing natural selection can be said to be learning. One way of formalizing this question is to ask under which conditions will the potential information 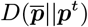 decrease? The potential information 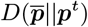 represents how much learning the population has left to do (i.e., how much information it lacks) from its current state. If the potential information is decreasing, then the population is making progress in learning ‘about’ the reference distribution 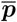.

To answer this question, we begin by calculating the rate of change of the potential information:

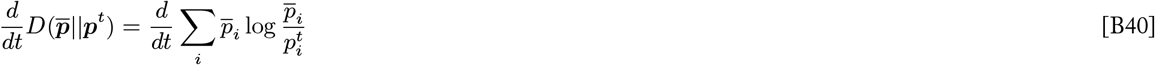

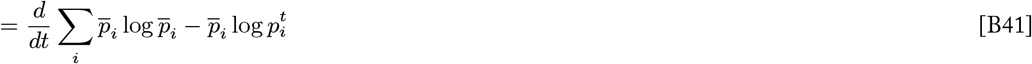

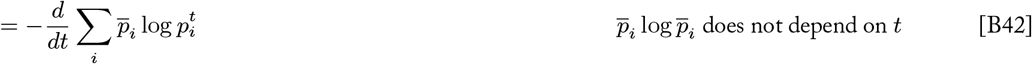

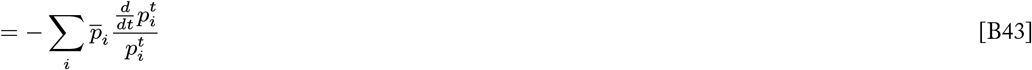

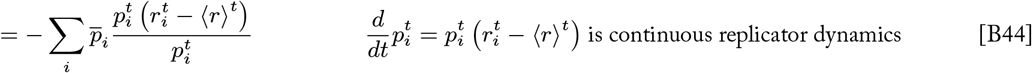

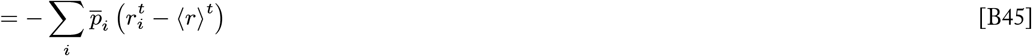

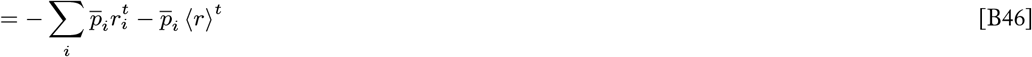

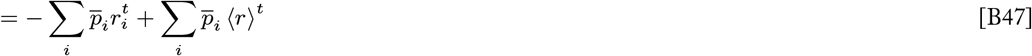

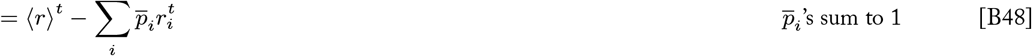

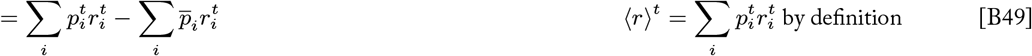

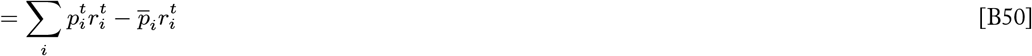

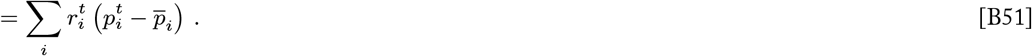

The potential information is decreasing when its rate of change is negative:

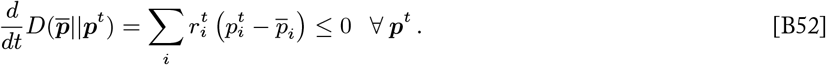

When this condition is met, the population can be guaranteed to be learning ‘about’ the reference distribution 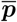.

A type frequency distribution is an evolutionarily stable state (ESS) if those frequencies are restored by selection after a small disturbance. In other words, a composition 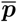 is evolutionarily stable if a small population of invaders with types distributed according to any other distribution ***p*** cannot out-compete the original population or perturb its frequency distribution. That is, a distribution 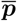 is an ESS if

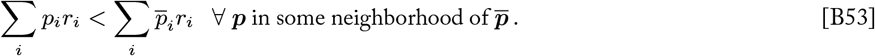

Basically, if the average fitness of a population with distribution 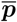 is greater than the average fitness of a population with any other distribution, then 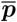 is an ESS.

We can rearrange the criteria for an ESS,

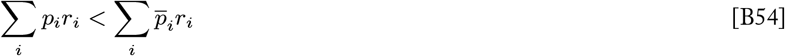

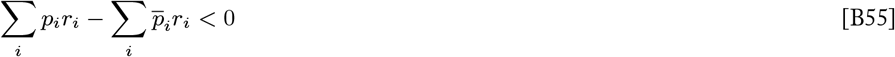

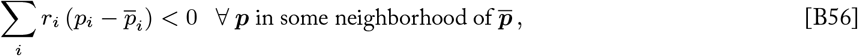

and realize that this is the same condition that guarantees that the potential information 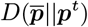 is always decreasing (Equation B52). Therefore, when an evolutionarily stable state 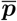 is accessible from the population’s current distribution state ***p*** (i.e., the population is in a basin of attraction for the ESS 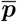), the potential information is guaranteed to be decreasing, and the population is guaranteed to be learning ‘about’ the ESS. As the population asymptotically approaches an evolutionarily stable composition 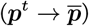, it gains all of the information that can be gained about the current environment 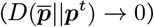.

Given an evolutionarily stable state 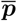, the potential information 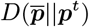 satisfies the criteria to be a Lyapunov function *V* for the replicator dynamic:

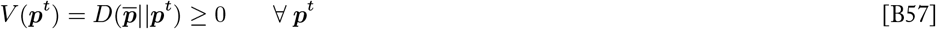

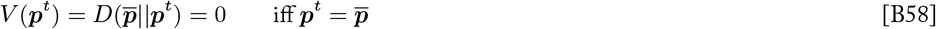

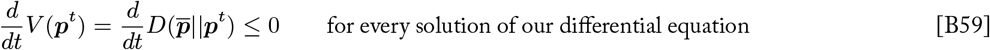

The existence of such a Lyapunov function is necessary and sufficient to prove the state 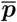 to be an asymptotically stable equilibrium. Therefore, we might say that evolutionary stability is fundamentally characterized in terms of information.

###### Potential information and coding inefficiency

In information theory, the KL divergence *D*(***q***||***p***) is often used to measure the inefficiency of encoding the state of a system that follows distribution ***p*** using a coding scheme that assumes a state distribution ***q*** rather than the true distribution. The meaning of inefficiency in this interpretation depends on the application. A classical example is encoding values of a random variable using binary strings. A coding scheme that seeks to use as few bits as possible might assign short binary strings to values that are presumed to have high probability and longer strings to values that are less common. In this case, the KL divergence *D*(***q***||***p***) measures how many more bits will be needed on average if such a code is devised under an assumption that the random variable takes values distributed according to ***q*** instead of the optimal distribution ***p***.

In the context of natural selection, the population’s type composition can be seen as encoding which types are most suited for the environment. Then the potential information 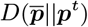 can be interpreted as the inefficiency of fielding a population according to the type distribution ***p***^*t*^ in an environment where the evolutionarily stable composition 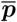 is in fact (locally) optimal. The inefficiency of the population’s coding scheme is manifest in its composition being mismatched with the distribution of environmental conditions to some extent, which results in lower than optimal expected fitness. As selection moves the population toward 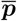, information is gained and the inefficiency of the population’s encoding is reduced. Understanding how this informational inefficiency changes over time and how it relates to the growth of the population are central aims of this work.

##### B.2.3 Natural selection increases mutual information between types and environments

Adaptive genetic information transmitted from one generation to the next provides the organism with some reduction in uncertainty about the environment. That is, if the genetic sequences that organisms possess are correlated with the environments they experience, then the genome encodes a representation of the environment that reduces uncertainty about prevailing conditions (Shea 2007). For example, imagine that you randomly draw a genotype from a population in some environment. If there is a relationship between environmental states and the genotypes you’re likely to observe in those states, then observing a particular genotype reduces your uncertainty about the environmental history to some extent. This notion of information is relevant to organisms, which receive information about how to get by in the current environment from a genotype drawn from the parental gene pool.

Natural selection establishes such a correlation between genetic sequences and environmental conditions. Due to the variation in traits conferred, not all genes are equally likely to be transmitted to the next generation. The sequences that are associated with phenotypes that confer high relative fitness in an environment increase in frequency, and an adaptive matching between genotypes and environments develops. Thus selection results in an accumulation of information about the environmental history. Previous work has shown that this kind of information has fitness value; that is, the long-term fitness of a lineage is proportional to the amount of information it has about the environment (Bergstrom and Lachmann 2004, Donaldson-Matasci et al. 2010, Rivoire and Leibler 2011, Rivoire 2016, Hilbert 2017).

Mutual information between genomes and environmental histories quantifies precisely this notion of information as the reduction in uncertainty (measured as entropy *H*) derived from the non-independent co-occurrence of genotypes and environmental conditions. Considering possible genotypes and environmental histories^*^ as random variables *G* (distributed according to *P* (*G*)) and *E* (distributed according to *P* (*E*)), respectively, the mutual information between genomes and environmental histories ℐ (*G*; *E*) is defined:

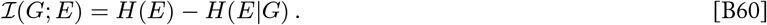

Equivalently, this mutual information measures the divergence of genotypes and environmental histories from independence:

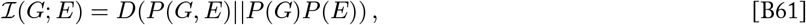

where *P* (*G, E*) is the joint probability distribution of *G* and *E*.

Let time *t* = 0 be defined as the time at which the current selection conditions begin. Genotypes and environments are assumed to be independent before the onset of these selection conditions, and their joint distribution at *t* = 0, *P*^0^ (*G, E*), reflects this initial independence:

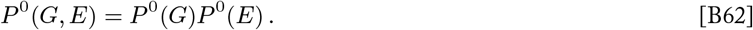

Natural selection establishes a covariance between genotypes and environmental histories due to environment-dependent changes in type frequencies. Therefore, the mutual information between genotypes and environments established due to a period of selection *T* can be quantified as the divergence of the post-selection (i.e., dependent) joint distribution of genotypes and environments from their pre-selection (i.e., independent) distributions

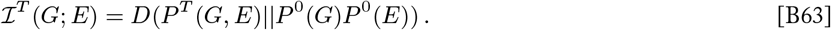

McGee and Bergstrom (2022) show that this mutual information can be evaluated as follows:

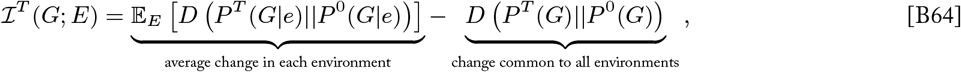

where *P* ^*T*^ (*G*|*e*) is the genotype frequency distribution in a particular environment *e* at time *T*, and *P* ^*T*^ (*G*) is the overall distribution of genotypes when populations from all environments under consideration are pooled together. This expression tells us that the mutual information ℐ^*T*^ (*G*; *E*) at time *T* can be decomposed as the average amount of change in genotype frequencies in each environment minus the change that is common across environments. Thus adaptive information refers to the amount of change that is *specific* to *particular* environments. Adaptive information accumulates when populations that experience different environmental histories evolve in distinct, environment-dependent ways. Therefore, adaptive genetic information is accumulated by selection but not by drift, unbiased random mutation, or migration in expectation (McGee and Bergstrom 2022).

Note that the distribution *P* ^*t*^(*G*|*e*) is synonymous the with type frequency distribution of a single population that occupies a particular environmental context, which we denote throughout this work as ***p***^*t*^. Then the expression for the mutual information accumulated through *T* generations of selection can be rewritten

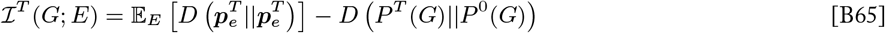

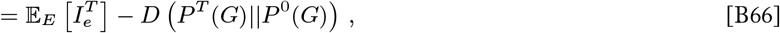

where 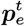 and 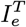 denote the type frequency distribution and the information gain (as in Equation 2, Appendix B.2.1) of the population experiencing a counterfactual environmental history *e*. Therefore, the accumulation mutual information between genotypes and environmental histories is equal to the average information gain of populations in different environmental contexts minus the genotype frequency change that is redundant across populations. This relates the information gain due to selection for a single population (the focus of this work) to the build-up of adaptive information about counterfactual environmental histories. (See McGee and Bergstrom (2022) for more on the meaning and measurement of this adaptive information.)

### Appendix C: Natural Selection as a Learning Process

#### C.1 Natural selection and Bayesian learning

Suppose that a learner wishes to infer the true state of a system, which may take any one of *n* possible states. The learner holds a hypothesis about the state of the system, which is represented by a probability distribution that assigns a weight Pr(*h*_*i*_) to each alternative. When new evidence *E* is obtained, the learner updates their prior hypothesis such that alternatives that give relatively high likelihood Pr(*E*|*h*_*i*_) to the observed evidence are up-weighted in the posterior hypothesis. Bayesian learning refers to this process of iteratively updating a hypothesis in light of new evidence using Bayes’ rule:

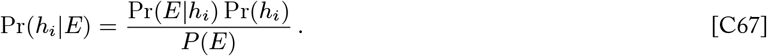

In the context of natural selection, each type in the population can be thought of as an alternative strategy for survival, and the distribution of type frequencies ***p*** can be seen as the population’s hypothesis about which types are most suited for the current environment. Like Bayesian learning, natural selection updates the population’s prior hypothesis in light of new evidence provided by the fitness landscape. As selection changes the frequencies of types according to their relative fitnesses, the distribution of types shifts to favor those that have generated organisms well-adapted to the environment. Indeed, there is a formal connection between Bayesian learning and replicator dynamics. These two update rules share the same form — selection increases the frequency of types with high relative fitness in the same way that Bayes’ rule increases the weight of alternatives that give high relative likelihood to the observed evidence:

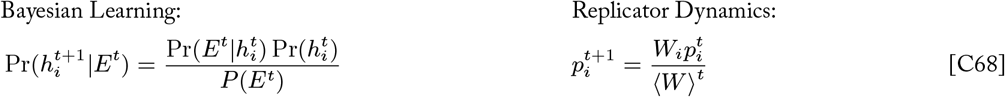

Natural selection can thus be understood as a learning process that “infers” which types are best adapted to the environment, and a number of recent studies have formalized selection and other evolutionary processes in the framework of Bayesian computations (Harper 2009b, Shalizi 2009, Campbell 2016, Watson and Szathmáry 2016, Czégel et al. 2020). In fact, Bayesian updating is a special case of discrete replicator dynamics, where the “fitness” of each alternative is given by its likelihood, and many results for replicator dynamics can be applied to Bayesian analysis more generally (Shalizi 2009).

#### C.2 Natural selection as an online learning process

##### C.2.1 The online learning problem

In the field of computational learning theory, *online learning* refers to a process of iteratively updating a strategy for responding to problems given feedback about the quality of past responses. That is, a learner is presented with a sequence of problems to which they must provide a response (e.g., an answer to a question, a prediction about the state of a system), and the learner holds a strategy that is used to generate their response to each problem in turn. After providing a response to each problem, the learner receives feedback about the quality of their response via a *loss* function that measures the discrepancy between the learner’s response and the optimal response for that problem. The learner’s goal is to minimize the cumulative loss that they suffer in the long run, which they may achieve by updating their strategy after each round such that their responses are more accurate in subsequent rounds. The problem of adapting a strategy for sequential prediction has been studied in a number of fields, including machine learning, game theory, and information theory.

Here we formalize the online learning problem in the game-theoretic framework of playing repeated games against Nature. In the following section, we interpret this formalism for the context of an evolving population (Appendix C.2.2). The repeated games framework is closely related to the prediction with expert advice framework that has been widely studied in computer science.

Consider repeated play of a two-player game in normal form. The game is defined by a matrix **G** with *n* rows and *m* columns representing the pure strategies available to the learner (row player) and the environment (column player), respectively. A *pure strategy* refers to the choice of a specific row or column, while a *mixed strategy* refers to a distribution over rows or columns. A *loss* function *ℓ*(*i, j*) defines the outcome associated with the learner’s pure strategy *i* being paired with the environment player’s pure strategy *j*. An optimal pure strategy *i* given the environment’s choice of pure strategy *j* receives zero loss (i.e., *ℓ*(*i, j*) = 0), whereas a pure strategy that fares more poorly against the environment’s play receives a correspondingly higher loss. The environment player’s payoffs or losses are left unspecified.

A play of the game in round *t* consists of the population player choosing a mixed strategy ***p***^*t*^ and the column player choosing a mixed strategy ***x***^*t*^. The **expected loss** of the learner when mixed strategies ***p***^*t*^ and ***x***^*t*^ are used is given by

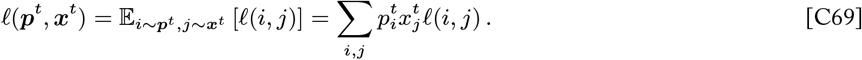

Following Freund and Schapire (1999), the loss that the learner suffers in each round is equal to the expected loss of their mixed strategy. The loss received by the learner is revealed after the selection of mixed strategies in each round of the game, but the environment player’s strategies and the game matrix as a whole are unknown to the learner. The goal of the learner is to minimize its **cumulative loss**

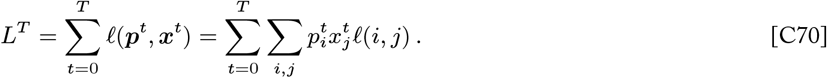

The learner may achieve this goal by iteratively updating their mixed strategy so as to learn a mixed strategy that has minimal per-round expected loss.

Depending on the definition of the game **G** and the environment player’s strategies ***x***^*t*^, even the optimal strategy available to the learner may still incur non-zero expected loss in each round, which results in a cumulative loss that continues to increase over time. In addition, we will consider the general case where we make no assumptions about the sequence of mixed strategies chosen by the environment player. In general, the environment may select strategies deterministically, stochastically, arbitrarily, or even adversarially (i.e., the environment can select ***x***^*t*^ with knowledge of ***p***^*t*^). An adversarial environment can make the learner’s cumulative loss arbitrarily large by choosing strategies ***x***^*t*^ that give a worst-case loss for the learner’s choice of ***p***^*t*^. For these reasons, it difficult to evaluate the relative performance of online learning algorithms on the basis of the learner’s cumulative loss alone.

Instead, online learning algorithms are commonly analyzed in terms of **regret**, which measures the difference between the cumulative loss of a learner and that of a competing fixed strategy ***q***.

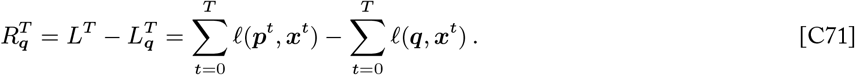

In computational learning theory, regret is typically evaluated with respect to the optimal strategy in hindsight

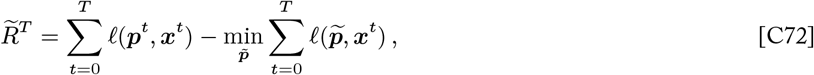

where the optimal strategy 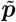 is defined as the fixed mixed strategy that minimizes the cumulative loss for the sequence of environment plays observed through round *T*. In this work, we also evaluate regret with respect to an optimal strategy that is defined in terms of an evolutionarily stable state, which has particular relevance for the learning problems faced by evolving populations (see Appendix C.3 for more information).

Regret analysis captures the ability of a learning algorithm to arrive at a strategy that does as well as an optimal strategy would in a given scenario. Thus, the learner’s goal can be restated as seeking to learn a mixed strategy that minimizes regret with respect to an optimal strategy ***q***. A learning process is said to be *no-regret* if it updates the learner’s mixed strategy such that their per-round regret vanishes in the long run, achieving

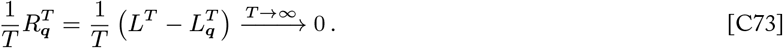

A no-regret learning process is optimal in the sense that it achieves the lowest possible per-round regret, which implies that the learner arrives at a strategy that has a per-round loss very nearly equal to that of the optimal strategy.

##### C.2.2 The Population versus Environment game

The repeated play formalism of the online learning problem (described in detail in Appendix C.2.1) can be interpreted as the basis for a model of natural selection in variable environments.^*^ We consider a game between two players: the population^†^ (acting as the learner) and the environment. The set of pure strategies available to the population player (i.e., rows of the game matrix) represents the set of replicator types in the population, and the population’s strategy ***p***^*t*^ represents the type frequency distribution of the population at time *t*. The set of pure strategies available to the environment player (i.e., columns of the game matrix) represent a set of possible environmental conditions that make up the environment. We assume that every individual in the population experiences an independent environmental condition, as if each individual occupies a separate part (e.g., micro-environment) of the physical environment. Then the environment player’s mixed strategy ***x***^*t*^ represents the probability distribution of environmental conditions that are experienced by individuals in the population. A play of the game in round *t* can be interpreted as the population assigning a type to each individual with frequencies ***p***^*t*^ and the environment assigning a condition to each micro-environment with probabilities ***x***^*t*^.

The outcome of each play of the game is scored according to a game matrix **G**. Each element of the game matrix *G*_*ij*_ gives the payoff for a type *i* that experiences an environmental condition *j*. Each individual of type *i* that experiences environmental condition *j* receives a loss *ℓ*(*i, j*). Typically the loss *ℓ*(*i, j*) is given by a function that measures the difference between the payoff for the *i*th type in the *j*th condition and the optimal payoff achievable in that condition. For example,

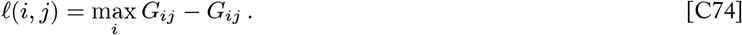

As we will see in Appendix C.2.4, payoffs and losses are closely related to fitness when considering a population player that updates their strategy according to natural selection in this setting. As in the general online learning setting described above, the loss suffered by the population player in round *t* is defined as the expected loss 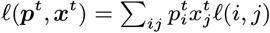, which is equivalent to the mean loss of all individuals in the population. The goal of the population player is to adapt its mixed strategy (type frequency distribution) ***p***^*t*^ in order to minimize its cumulative loss and achieve low regret. As we will see in Appendix C.2.4, the replicator dynamics model of natural selection is equivalent to an online learning algorithm for precisely this setting.

We take a very expansive view of what constitutes an “environment.” Environmental conditions may represent the abiotic conditions experienced by individuals, the types of other replicators encountered by individuals, or even the identity of other alleles carried by individuals. Similarly, the environment player may select their sequence of strategies {***x***^0^, ***x***^1^, …, ***x***^*T*^ } according to any process, including as a function of time, as a function of the population player’s strategy (i.e., as a function of type frequencies), as a stochastic or random process, as a “pre-programmed” sequence, etc. While the environment may select the frequency distribution of conditions ***x***^*t*^ with knowledge of the population’s type frequency distribution ***p***^*t*^, the association of particular environmental conditions (micro-environments) to specific individuals is assumed to be random and out of the control of both players. Throughout this work we consider various environmental contexts of interest as well as the fully general setting where no assumptions are made about the sequence of environmental condition distributions.

##### C.2.3 The Multiplicative Weights Updating (MWU) algorithm

Online learning problems of this form are common, and a simple yet powerful learning algorithm known as **Multiplicative Weights Updating (MWU)** has been shown to be an effective solution to such problems in computer science, machine learning, economics, and other fields. Here we highlight the conceptual intuitions underlying this algorithm by going through a loss-optimization derivation of MWU before stating the MWU update rule.

###### Derivation of MWU

[*This section derives the MWU update rule using the method of Lagrange multipliers, the details of which are not critical for understanding this work. The logic that sets up the optimization may provide some insights, even if Lagrange optimization is unfamiliar. Readers simply interested in the definition of MWU can skip to the next section*.]

A learner facing an online learning problem seeks a method for adaptively selecting strategies such that their long-term cumulative loss is minimized. Intuitively, it might seem reasonable for the learner to choose the mixed strategy that would perform the best against the previously observed plays of the environment—this is known as the **Follow the Leader (FTL)** algorithm:

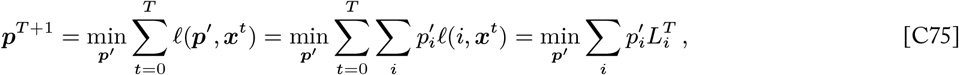

where 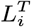 is the cumulative expected loss of the pure strategy *i*. This update rule effectively places all of the weight on the single pure strategy that has seen the lowest cumulative loss so far. This Follow the Leader algorithm is inherently unstable as all of the weight can switch from one pure strategy to another from round to round. Furthermore, there is no guarantee that a strategy that performs well on past observations will continue to do so against future plays of the environment, so a learner using this algorithm can be easily tricked into adopting strategies with poor long-term loss.

In order to add stability to the mixed strategy updates, we can add a regularization term ℛ (***p***) to the optimization that generates the updated mixed strategy, where the regularizer ℛ is some convex function. An appropriately chosen regularizer function can offset the influence of the loss minimization term by promoting more uniform weight distributions that do not change too quickly. Such regularization makes the algorithm less vulnerable to short term plays of the environment that might otherwise fool the learner into adopting strategies with high long-term loss. The resulting update rule is known as the **Follow the Regularized Leader (FTRL)** algorithm:

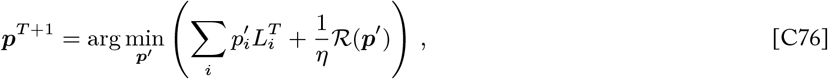

where *η* ∈ (0, ∞) is a learning rate parameter that modulates the relative emphasis that is given to minimizing loss in hindsight versus constraining change in each update via the regularization term. When the learning rate *η* is large, the weights can change greatly in each update (approximating the un-regularized FTL as *η* → ∞). When the learning rate is small, the FTRL update favors the regularization criteria, which typically prevents weight consolidating on a small number of pure strategies in the short-term. In general, the FTRL algorithm balances minimizing long-term loss against maintaining diversity in the mixed strategy distribution during the transient of the learning process.

Intuitively, the negative entropy −*H*(***p***) = Σ_*i*_ *p*_*i*_ log *p*_*i*_ is a reasonable choice for the regularization function. First, the negative entropy is a strongly convex function, which is a requirement of our choice of regularizer function. More importantly, entropy is a measure of the spread of a distribution that is maximized for the uniform distribution, so the *negative* entropy is *minimized* for the uniform distribution. Therefore, using the negative entropy of the mixed strategy as the regularizer will lead the minimization in the FTRL update rule to balance loss minimization against maintaining more uniform weight distributions:

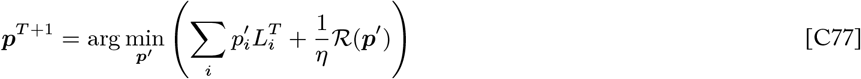

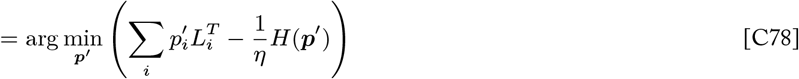

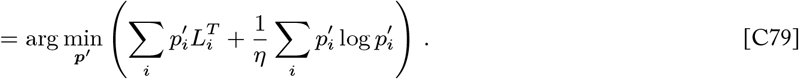

We can set out to solve for the distribution ***p*** ′ that achieves this FTRL update by finding the minimum of this expression. This is a straightforward optimization problem, because the expression to be minimized is strictly concave and has a unique minimum. We solve this optimization using the method of Lagrange multipliers, where we include the constraint that 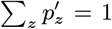. We begin by defining the Lagrange function ℒ (not to be confused with *L*, which denotes cumulative loss)

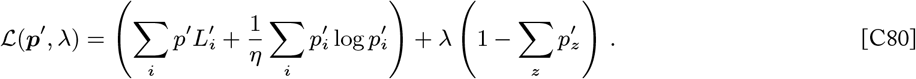

Set the derivative of the Lagrange function equal to 0 and evaluate the system of equations that includes the contraint

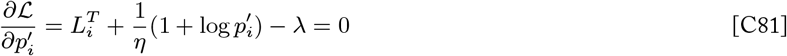

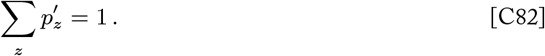

Solving for 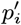 in terms of *λ*

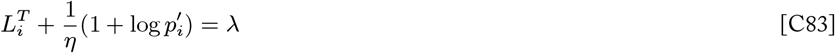

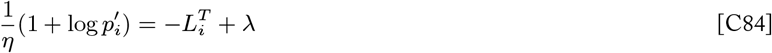

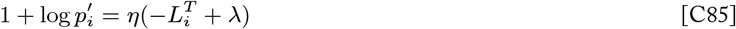

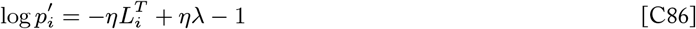

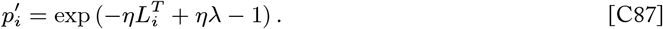

Plugging this expression for 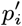 into the constraint to solve for *λ*

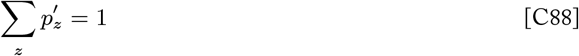

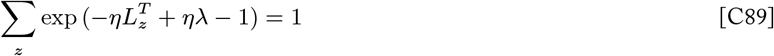

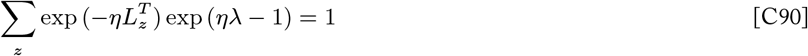

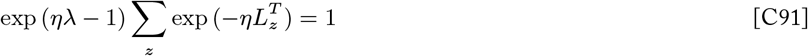

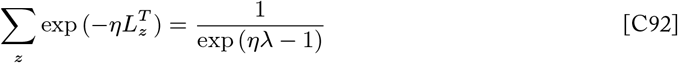

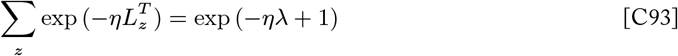

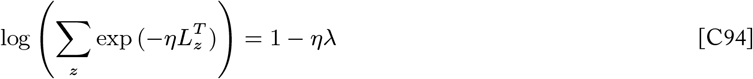

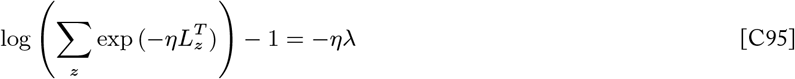

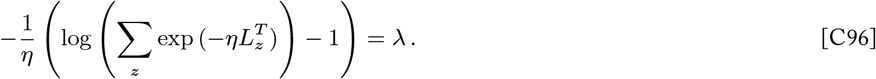

Plugging this expression for *λ* into the expression we previously found for 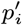 in terms of *λ* (Equation C87) we can ultimately solve

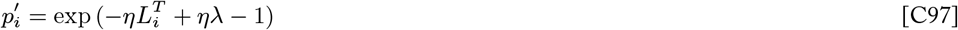

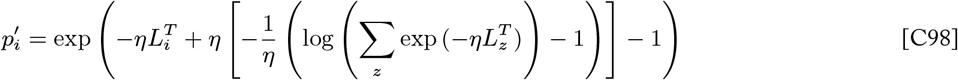

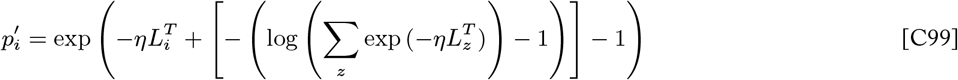

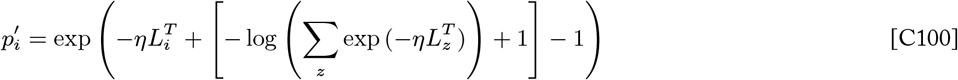

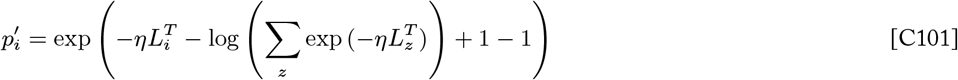

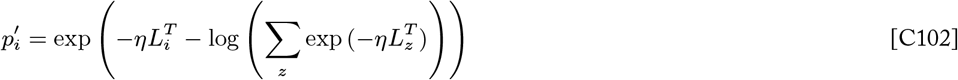

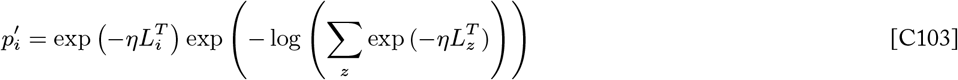

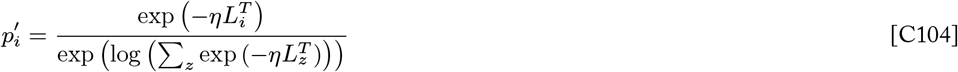

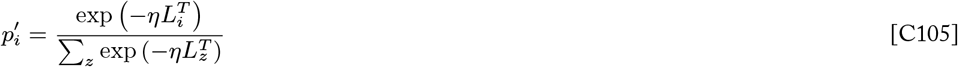

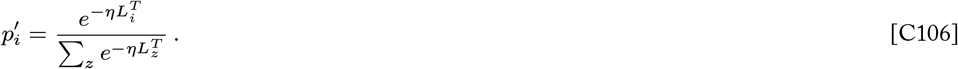

We have found an expression for the mixed strategy distribution ***p***′ that solves the FTRL optimization problem (Equation C79). Therefore, we find that applying the update 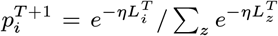 for each pure strategy *i* updates the mixed strategy to a distribution that minimizes a combination of expected cumulative loss and negative entropy.

This is a useful update rule in and of itself, but we can go a step further to put this update rule in a form that depends only on the player’s loss in the most recent round, which conveniently does not require tracking cumulative loss. Using the recursion 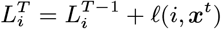, we can express this update rule in terms of the most recent single-round losses alone:

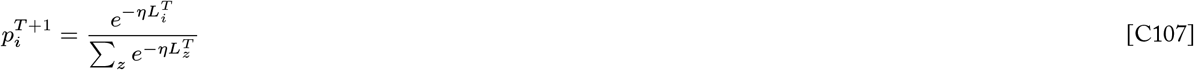

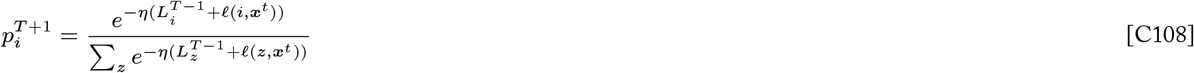

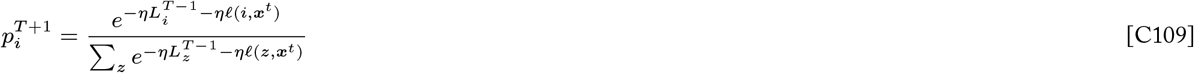

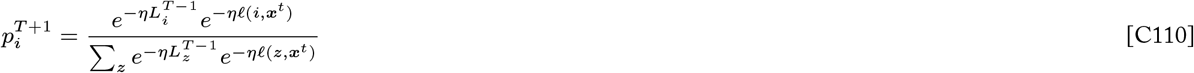

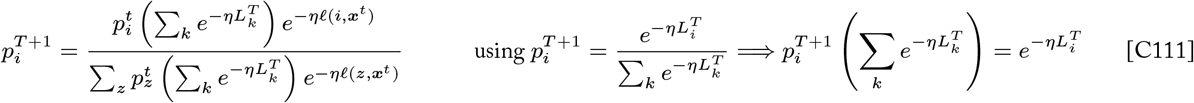

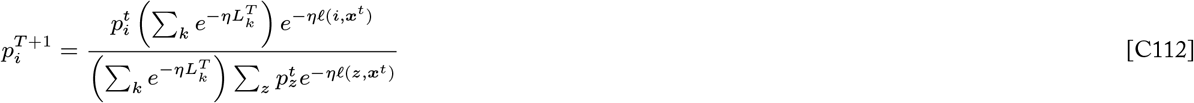

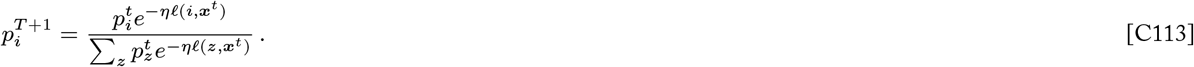

Finally, we have an update rule (Equation C113) that implements FTRL with negative entropy regularization.

###### The MWU Update Rule

In the previous section, we derived an update rule that implements the *Follow the Regularized Leader* algorithm with negative entropy regularization, and in so doing updates the learner’s mixed strategy distribution so as to optimize a potential function of expected total loss and strategy entropy. This update rule has been repeatedly derived in many contexts and goes by several names, including **Multiplicative Weights Updating (MWU)**:

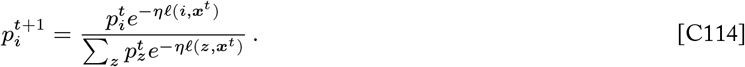

This algorithm is no-regret in many contexts and achieves reasonable regret bounds in general. This means that a learner who uses MWU to update their strategy learns an approximation of an optimal strategy that achieves a cumulative loss that is nearly as low as that of the optimal strategy. In some settings, such as where the learning rate can be tuned or where the loss function has certain properties, it can be shown that MWU is an optimal learning process in the sense that no other learning algorithm can possibly achieve a tighter bound on regret than MWU (Freund and Schapire 1999). In this work, we are particularly interested in the regime where MWU is equivalent to natural selection (see Appendix C.2.4). Regret bounds for natural selection are given in Appendix D.3.

##### C.2.4 Natural selection as a fitness-based instance of MWU

In Appendix C.2.2 we establish the online learning problem as the basis for a model of evolution, and in Appendix C.2.3 we derive the MWU update rule, which is known to be an effective learning process for problems of this kind. Recent work has shown that discrete-time replicator dynamics is equivalent to an instantiation of the MWU learning process (Chastain et al. 2014, Mehta et al. 2015, Meir and Parkes 2015, Chastain 2017). In this section, we complete the connection between online learning and natural selection by describing the equivalence of replicator dynamics and MWU and interpreting the loss function and learning rate implicit in this equivalence.

It is straightforward to recognize that MWU and replicator dynamics^*^ have the same form — MWU increases the weight of alternatives that have relatively low loss, and selection increases the frequency of types with high relative fitness:

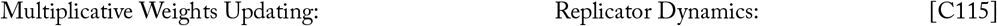

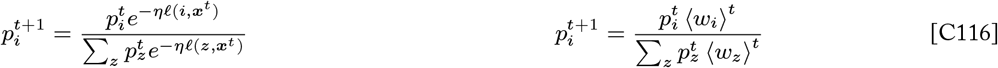

These two update rules are equivalent where the following identity holds

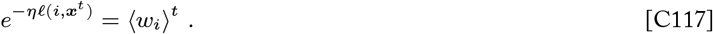

Evaluating this equation further we have

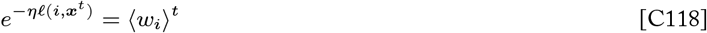

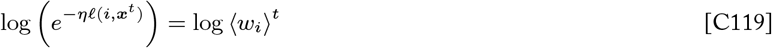

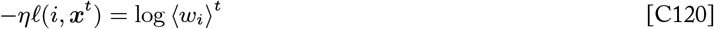

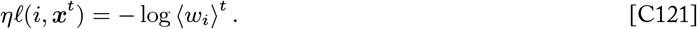

At this point, if we let *η* = 1 then the expected loss of the *i*th type ought to be equal to the negative log expected relative fitness of the *i*th type.

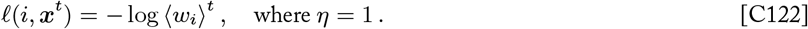

From here we would like to find the loss function *ℓ*(*i, j*) for which this equivalence holds. Let us propose that the choice of loss function *ℓ*(*i, j*) = − log *w*_*ij*_ satisfies this equation, which we will now prove.

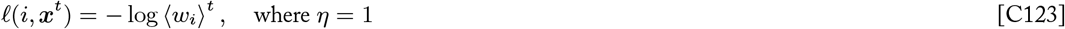

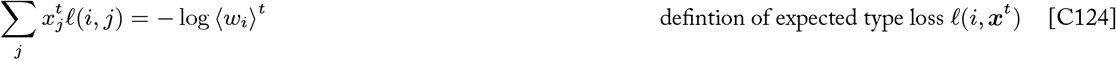

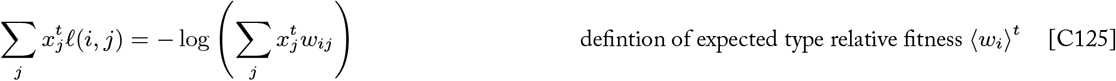

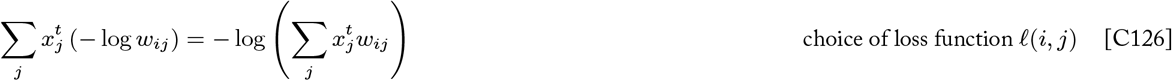

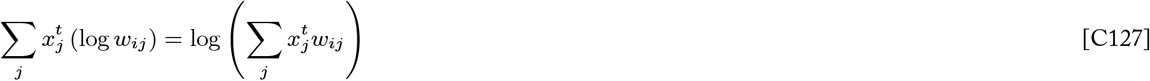

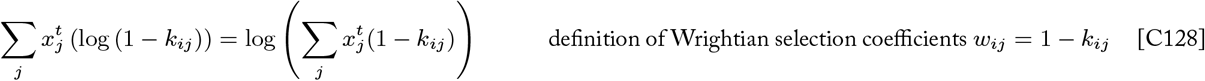

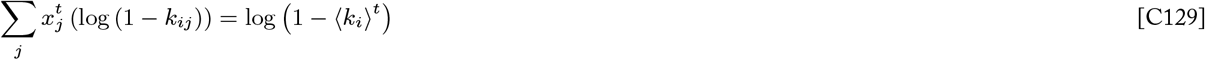

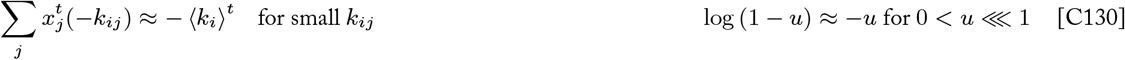

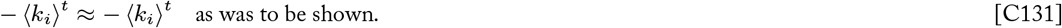

Therefore, Multiplicative Weights Updating is equivalent to discrete-time replicator dynamics for haploid asexual populations under the following conditions:

1. Loss function *ℓ*(*i, j*) = − log *w*_*ij*_.
2. Learning rate *η* = 1.
3. In the limit of weak selection, where *k*_*ij*_ ⋘ 1 for all *i, j* (or in the limit of continuously overlapping generations).

Using the definition of relative fitness *w*_*ij*_, the loss function *ℓ*(*i, j*) = − log *w*_*ij*_ can be expressed and interpreted in another form.

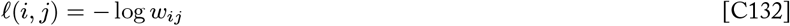

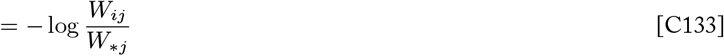

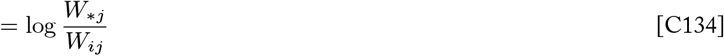

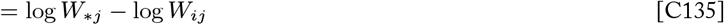

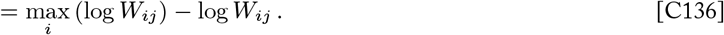

Thus we see that this loss function measures the difference between the log fitness (growth rate) for the *i*th type in the *j*th condition and the optimal payoff achievable in that condition. This is a sensible way to define the validity of the individual “response” *i* to the “context” *j* in learning-theoretic terms. Moreover, this difference gives the loss of potential fitness that an individual experiences by having a suboptimal type for the condition they experience, which is meaningful in the context of a population evolving by natural selection. Identifying log fitness as the payoff compared by the loss function (Equation C74) tells us that the *n*-by-*m* matrix **G** defining the Population versus Environment game gives the log fitnesses (growth rates) of all *n* types in all *m* conditions (i.e, *G*_*ij*_= log*W*_*ij*_).

The learning rate *η* sets the relative emphasis that Multiplicative Weight Updating gives to minimizing loss based on previous options versus maintaining spread over types as a form of regularization (Equation C76). Learning rates *η >* 1 react strongly to each observed round of losses, while learning rates *η <* 1 put more emphasis on maintaining entropy in the learner’s strategy. The fact that a learning rate *η* = 1 is implicit in the equivalence between natural selection and MWU tells us that selection balances maximizing expected cumulative fitness (minimizing expected fitness loss) and maintaining diversity exactly equally (Chastain et al. 2014).

#### C.3 Two classes of learning problems for evolving populations

For a population that learns by natural selection, it is useful to distinguish two classes of learning problems that lead to different conceptions of strategy optimality and corresponding definitions of regret.

The nature of the learning problem faced by a population depends on the sequence of environments it experiences. A given distribution of environmental conditions ***x***^*t*^ can be interpreted as determining a *fitness landscape*: the set of expected fitnesses for each type in the population (i.e., the vector **G*x***^*t*^ gives the fitness landscape at time *t*). The central problem for the evolving population can be seen to be arriving at an equilibrium state that maximizes fitness locally with respect to the current fitness landscape. The dynamics of selection continually move the population toward an accessible stable state, which decreases potential information and increases fitness with respect to the present environment. In other words, what selection learns is an evolutionarily stable type composition for the current conditions. When the makeup of the environment changes the fitness landscape changes as well, the evolutionarily stable state that represents the population’s learning target may or may not change depending on the particular sequence of changes to the fitness landscape. A sequence of environments for which the evolutionarily stable state accessible to the population remains stationary represents a **fixed learning problem**. On the other hand, many sequences of environments do cause the evolutionarily stable states to change over time, which we refer to as **variable learning problems**. These classes of learning problems are defined formally below.

**Definition**. *A type distribution* 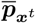 *is an* **evolutionarily stable state** *of replicator dynamics with respect to the distribution of environmental conditions x*^*t*^ *if for all distributions* 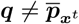 *in a neighborhood* 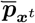

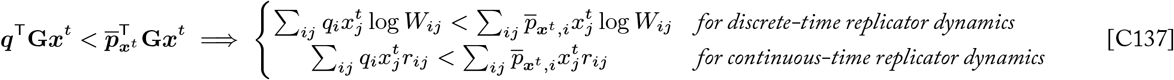

**Definition**. *The* **basin of attraction** *of an evolutionarily stable state* 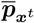 *is the set of all initial conditions from which trajectories of replicator dynamics approach* 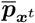. *Formally, the basin of attraction* 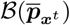 *is a positively invariant set (i.e*., *all trajectories starting in* 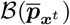 *remain in* 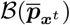) *that contains* 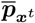 *and for which there exists a Lyapunov function V* (***p***) *that satisfies*

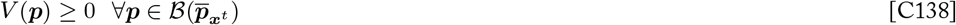

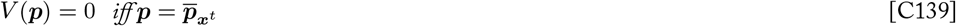

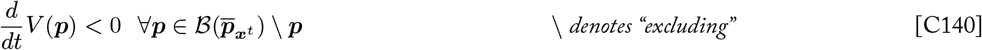

*Then, by La Salle’s Invariance Principle, every trajectory starting at* 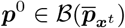 *tends to* 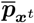 *as t* → ∞.

**Definition**. *An evolutionarily stable state* 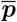 *is said to be* **stationary** *over a sequence of environments* ***x***^0^, …, ***x***^*T*^ *if the equilibrium point remains constant and the population’s trajectory p*^*t*^ *remains within its basin of attraction throughout the sequence. That is*,

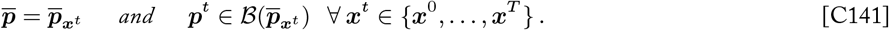

**Definition**. *Any sequence of environments* ***x***^0^, …, ***x***^*T*^ *for which the population’s initial type distribution p*^0^ *is in the basin of attraction of a stationary evolutionarily stable state* 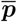 *is said to represent a* **fixed learning problem** *for the population*.

**Definition**. *Any sequence of environments* ***x***^0^, …, ***x***^*T*^ *that does not meet the criteria of a fixed learning problem — that is, where the evolutionarily stable state accessible to the population changes throughout the sequence of environments — is said to represent a* **variable learning problem**.

### Appendix D: The Cost of Natural Selection

#### D.1 Mismatch load quantifies the cumulative fitness loss of selection

The *genetic load* of a population refers to “the extent to which [the population] is impaired by the fact that not all individuals in the population are of the optimum type” (Crow 1958). A population that includes suboptimal types for the current environment will have a lower average population fitness than a population that has the optimal composition, and genetic load measures the proportion by which the population fitness is decreased due to the presence of the inferior types. A number of factors can contribute to the presence of suboptimal types in a population and thus to the overall genetic load. For example, *mutation load* refers to the depression in population fitness due to the occurrence of deleterious mutations, and *segregation load* refers to a depression in population fitness due to the production of inferior homozygotes by allele segregation in the context of heterozygote advantage (Crow 1958, Kimura 1960). The component of load that is the focus of this paper is the *substitution load* and its generalization, *mismatch load* : the depression in population fitness that results from mismatch of types and environmental conditions.

##### D.1.1 Substitution load

In the following sections, we walk through Haldane’s and Kimura’s definitions of substitution load in terms of Wrightian fitnesses (i.e., discrete generations) and Malthusian fitness (i.e., continuous growth), respectively (see Appendix A for fitness definitions).

###### Substitution load for discrete-time replicator dynamics

The basic idea of genetic load was introduced by J.B.S. Haldane (1937). Haldane (1957) was also the first to consider a “cost of natural selection,” in which he quantified the number of selective deaths that must occur in order to replace one allele with a more fit one. Haldane (1957) considers a haploid population with discrete generations that undergoes selection in response to a change in the environment. The result of this change is that one or more types that were previously rare become beneficial. Overall, the population is less adapted to the new environment, and its reproductive capacity is lowered due to the predominance of types that have poor viability, fertility, etc. in these conditions. Natural selection gradually improves the composition of the population, but “meanwhile a number of [selective] deaths, or their equivalents in fertility, have occurred” (Haldane 1957). Haldane sets out to quantify the total number of “selective deaths” that occur during the process of selection substituting the newly optimal type.

Let the absolute Wrightian fitness of each type *i* in the new environment be given by *W*_*i*_.* The fitness of the *ith* type relative to the fitness of the optimal type is given by

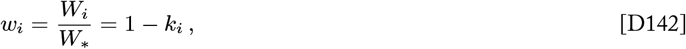

where *k*_*i*_ ∈ [0, 1] is the Wrightian selection coefficient of the *i*th type, and *k*_∗_ = 0 is the selection coefficient of the optimal type. In other words, 1 − *k*_*i*_ individuals of type *i* survive and reproduce for every one of the optimal type. Therefore the selection coefficient *k*_*i*_ can be interpreted as the per capita number of “selective deaths” of type *i*. Then the expected number of selective deaths in the population in single a generation is given by

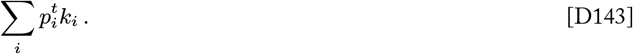

Haldane (1957) goes on the evaluate the total number of expected selective deaths accumulated over *T* generations of selection

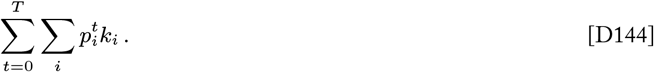

Motoo Kimura (1960, 1961) further developed this concept into the more general definition of substitution load. Kimura defined substitution load as the cumulative difference between a population’s average growth rate and the growth rate of the optimal type over the course of an allele substitution. Kimura’s definition clarifies that selection and the resulting substitution load need not involve actual deaths.

While Kimura defined this quantity in terms of continuous growth rates (see the next section), we consider an analogous definition for discrete generations in this paper. The log fitness log *W*_*i*_ gives the expected instantaneous growth rate of the subpopulation of type *i* individuals in a single generation (Appendix A.1.1). The difference between the growth rate of the optimal type and the population’s average growth rate in a single generation is then given by

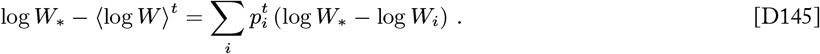

This difference gives the expected depression in single-generation growth that the population experiences due to its mixed composition relative to a population that consists of only the optimal type. Natural selection will gradually increase the the frequency of the optimal type in the evolving population, and this difference will decrease over time. Substitution load 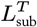 is defined as the total sum difference between the evolving population’s growth rate and the optimal growth rate over *T* generations of a gene substitution process in a constant environment

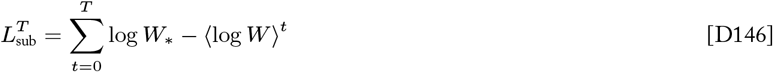

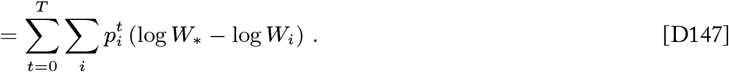

This quantity gives the expected fold difference in long-term growth of the evolving population relative to a population with an optimal composition, where the ‘fold’ refers to the base of the logarithms. When logarithms to base 2 are used, a substitution load 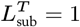 refers to 1 two-fold reduction in the population’s long term growth relative to the optimum.

Kimura’s substitution load is equivalent to Haldane’s expression for the number of selective deaths in the limit of weak selection or continuously overlapping generations. To see this, we can work out an expression for the discrete-time definition of substitution load in terms of Wrightian selection coefficients.

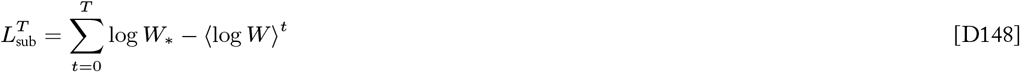

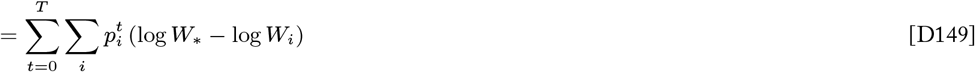

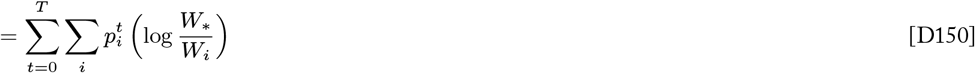

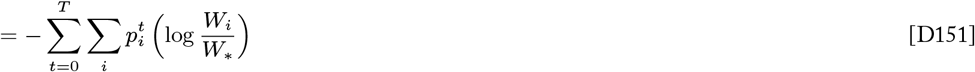

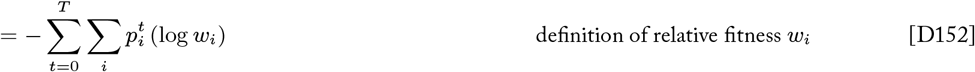

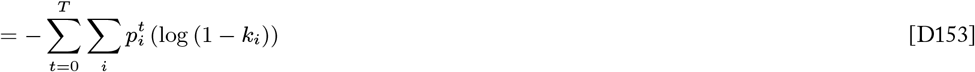

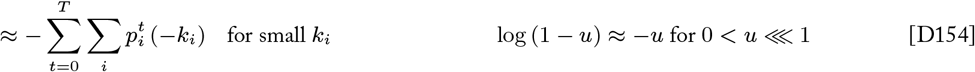

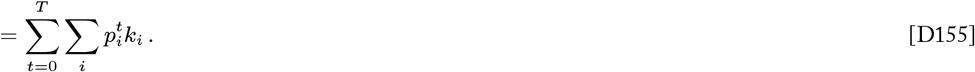

The final line is equal to the number of selective deaths in Haldane’s model (Equation D144), and thus Haldane’s quantity is equivalent to discrete-time substitution load in the limit of weak selection (small *k*_*i*_ for all *i*). The connection between the discrete generations and continuous growth definitions of substitution load can be seen in the following section.

###### Substitution load for continuous-time replicator dynamics

Now we turn to Kimura’s definition of substitution load for haploid populations with overlapping generations in a constant, homogeneous environment (Kimura 1960, 1961). Consider a population that consists of multiple types^*^, where 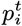 and *r*_*i*_ give the frequency and Malthusian fitness (exponential growth rate) of the *i*-th type at time *t*, respectively. Suppose the optimal type, which confers the maximum growth rate *r*_∗_ = max_*i*_ *r*_*i*_, has an initial frequency 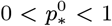. The difference between the growth rate of the optimal type and that of the *i*th type is given by the Malthusian selection coefficient

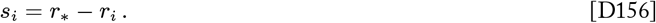

This fitness difference can be thought of as the growth penalty that the population receives for each individual that is of the *i*th type rather than the optimal type for the environment. Then the population’s expected fitness reduction due to the presence of inferior types at a given time *t* is the frequency-weighted average of these fitness differences

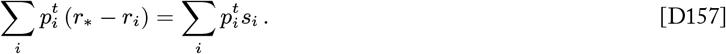

This expected fitness reduction integrated over a period of selection of duration *T* defines the **substitution load**

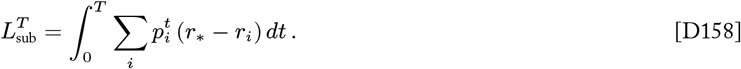

Substitution load can be equivalently expressed as the cumulative difference between the optimal fitness and the population’s mean fitness 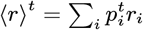

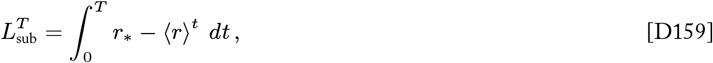

or as the integral of the population’s mean selection coefficient 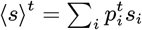

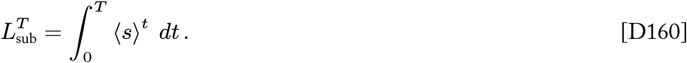

We can evaluate this integral to obtain a closed-form expression for substitution load. Evaluating this integral with respect to time is not straight-forward, but we can make use of the definition of continuous-time replicator dynamics to change the integration variable to one that is simpler to work with. Recall the continuous-time replicator dynamics that governs how type frequencies change over time

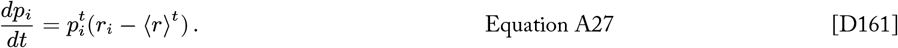

We can rearrange this differential equation to find an expression for *dt* in terms of the frequency and fitness variables.

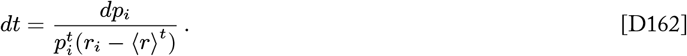

If we consider the replicator dynamics for the optimal allele in particular, the expression we obtain for *dt* includes in the denominator a term that appears in the substitution load integral, which will prove useful for simplifying the integral

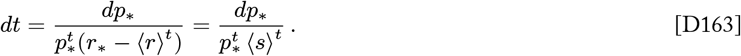

Now we can use this expression for *dt* to change the integration variable and evaluate the simplified integral

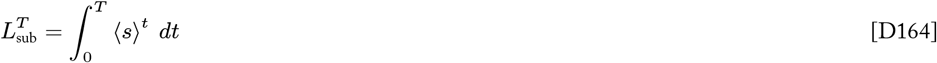

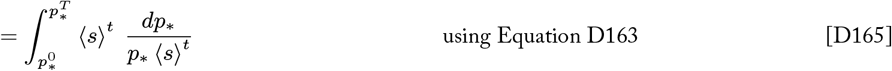

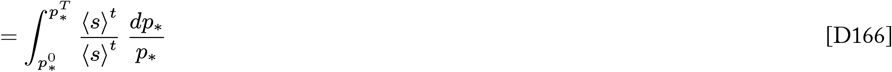

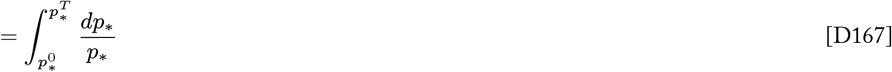

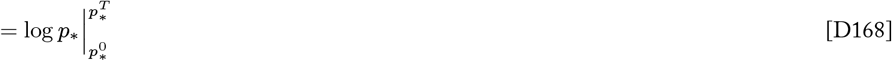

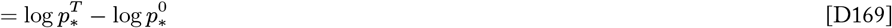

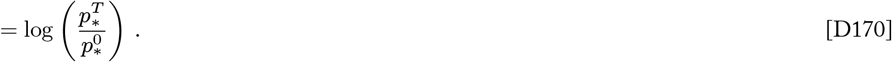

Here we see that the substitution load after a period of selection of duration *T* is a function of the pre- and post-selection frequencies of the optimal allele. In the long run, as the population approaches fixation for the optimal allele, the total substitution load converges to

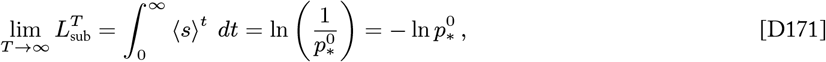

a value that depends only on the initial frequency of the optimal allele. Interestingly, the total substitution load for a complete allele substitution does not depend on the strength of selection. The relative fitness advantage of the optimal allele over other alleles in the population will modulate the time it takes for selection to fix this allele, but the total load will be the same in any case.

##### D.1.2 Mismatch load

The classical definitions of substitution load introduced by Haldane (1957) and Kimura (1960, 1961) consider a model of evolution in a homogeneous and constant environment. In such a case, there is a strictly optimal type with maximal fitness for the given environment, and the result of natural selection is a sweep where the optimal type substitutes all others. Substitution load measures the cumulative loss of fitness associated with selection completing this substitution gradually. This is a useful thing to measure, but this basic setting is restrictive.

We would like to extend this notion of load to more general settings with heterogeneous and time-varying environments. In Appendix A.2 and Appendix C.2.2 we present a model of evolution in variable environments. The environment is considered to be constituted by a set of distinct conditions. Each individual in the population experiences an independent micro-environment characterized by particular condition, and the fitness *W*_*ij*_ (or *r*_*ij*_ for continuous dynamics) of each individual depends both on its type *i* and on the environmental condition *j* that it experiences. The core concept that substitution load measures is the accumulation of fitness losses due to the presence of types that are poorly suited for the conditions they experience. In a heterogeneous environment, different environmental conditions may favor different types. In principle, overall fitness is maximized if every individual possesses the type that is optimal for the specific environmental condition that they experience. However, we assume that types are distributed over conditions randomly, so some type-condition mismatch is inevitable. We introduce the concept of **mismatch load**, which quantifies the cumulative loss of potential fitness due to the mismatch of types and environmental conditions.

Consider an environmental condition *j*. An individual of type *i* that experiences this condition has fitness *W*_*ij*_, while the fitness of the type that is optimal for this condition is denoted by *W*_∗*j*_ = max_*i*_ *W*_*ij*_. The amount of potential fitness that an individual misses out on by having a suboptimal type *i* in condition *j* is given by the difference

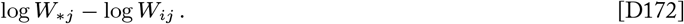

The expected amount of fitness loss incurred by the population due to type-condition mismatch over the distribution of types ***p***^*t*^ and distribution of conditions ***x***^*t*^ in generation *t* is given by

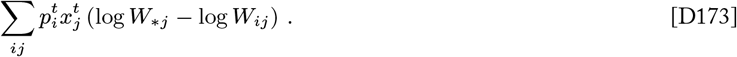

Mismatch load *L*^*T*^ measures the cumulative loss of fitness due to type-condition mismatch over *T* generations

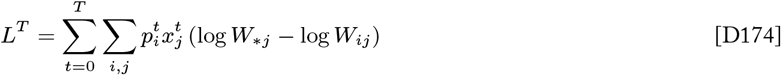

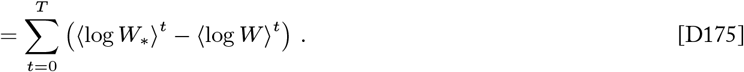

Mismatch load is equivalent to substitution load (Equation D147) when there is only one environmental condition (i.e., *m* = 1, ***x***^*t*^ is a point distribution). Like substitution load, mismatch load quantifies the fold reduction in total growth that an evolving population experiences due to having a composition that is not optimally matched to the environmental conditions. In the case of a gene substitution in a constant environment, substitution load converges on a finite value as selection approaches fixation of the optimal type. However, when the environment is heterogeneous there may not be a single type that is universally optimal in all conditions. In such a case, there is no type composition that can be expected to achieve zero mismatch given random associations of types with conditions. Therefore, mismatch load does not necessarily converge on a finite value and may accrue even for the type composition that gives the optimal expected fitness over all types and conditions.

###### Mismatch load is equivalent to cumulative expected fitness loss

In Appendix C.2.4 we show that replicator dynamics are equivalent to an instantiation of the MWU online learning algorithm characterized by the learning rate *η* = 1 and fitness-based loss function

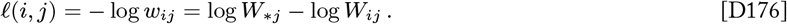

This function defines the *loss* that an individual of type *i* incurs in environmental condition *j*. In biological terms, this loss function gives the loss of potential fitness that an individual of type *i* experiences due to having a suboptimal type in condition *j*. The expected loss of all type *i* individuals taken over the distribution of environmental conditions ***x***^*t*^ at time *t* is given by

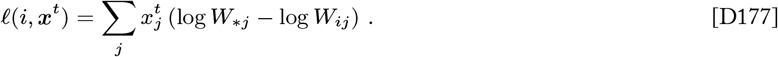

The expected loss of fitness experienced by the overall population in generation *t* (i.e., the expected loss taken by the population player in round *t* of the Population versus Environment game) is given by

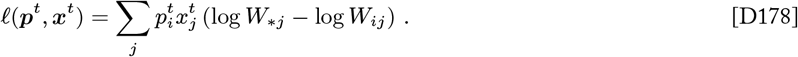

Then the cumulative expected loss of the population summed over *T* generations (i.e., *T* rounds of the PvE game) is given by

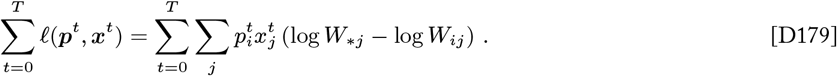

We recognize this quantity as being identical to the definition of mismatch load (Equation 8, Equation D175, Appendix D.1.2)

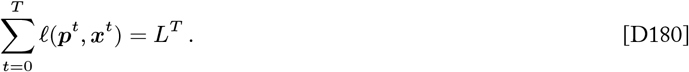

This connection reinforces the meaning of mismatch load and makes intuitive sense. Natural selection is equivalent to the MWU online learning algorithm when the learning-theoretic loss function is defined as the fitness loss each individual incurs given the environmental conditions they experience. The more type-condition mismatch occurs, the more loss the population is expected to incur and accrue, both in terms of learning-theoretic loss and long-term fitness loss.

#### D.2 Regret quantifies the cost of selection

In Appendix C.2 we introduced the concept of regret in the context of online learning. Regret is used in learning theory to evaluate the performance of a learning process by comparing the responses of a learner to the responses produced by a fixed reference strategy. Responses are typically scored using a loss function that quantifies the validity of a response relative to the best response in a given situation. When the reference strategy is taken to be optimal, regret represents the cost of the learning process: the excess loss a learner suffers by having to learn an appropriate response as opposed to knowing the solution all along.

In this section, we show the connection between regret and load and establish two definitions of regret in the context of an evolving population.

##### D.2.1 Regret quantifies relative mismatch load

In this section, we evaluate the definition of regret that applies to natural selection as a learning process. In general, regret is defined as the difference between the cumulative expected loss of the learner and the cumulative expected loss of a fixed reference strategy ***q*** (Appendix C.2.1).

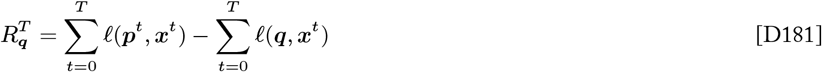

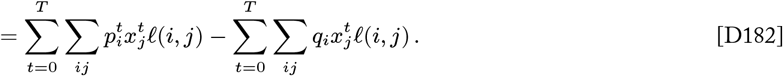

If we plug in the loss function *ℓ*(*i, j*) = log *W*_∗*j*_ − log *W*_*ij*_ that is implicated in using natural selection as the learning process (Appendix C.2.4) we have

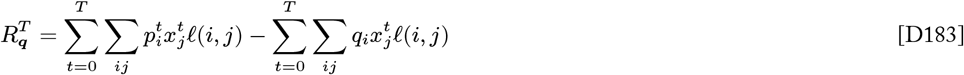

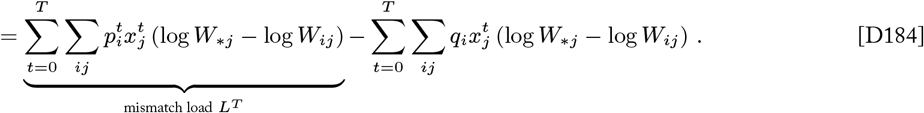

In Appendix D.1.2 we show that the cumulative loss of natural selection is equivalent to the definition of mismatch load (Equation 8, Equation D175, Appendix D.1.2). The second term has the same form, but is calculated with respect to the fixed reference distribution ***q***. Therefore we can interpret the regret of selection as the difference between the evolving population’s mismatch load and the mismatch load experienced by the reference type composition ***q***

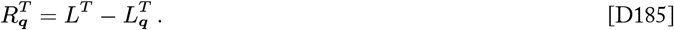

Thus regret represents a measure of relative mismatch load. If the reference distribution ***q*** is taken to be an optimal type composition in some sense, then regret measures the amount by which the mismatch load experienced by the evolving population exceeds the mismatch load it could have achieved if it adopted the optimal fixed composition from the beginning. This is a rigorous measure of the cost of the selection process that quantifies how much loss of long-term fitness the population incurs by gradually evolving as opposed to enjoying an optimally adapted composition from the beginning. In this work we consider the regret of selection with respect to two alternative definitions of fixed optimal compositions; see Appendix D.2.3 for more details.

##### D.2.2 Regret is equivalent to relative lineage fitness

We have shown that regret is a measure of relative mismatch load in the context of natural selection, and thus regret quantifies the amount of cumulative fitness loss an evolving population experiences beyond that of some reference composition. The excess fitness loss that corresponds to regret causes the evolving population to grow more slowly than a hypothetical population that has the reference composition all along. As such, we can interpret the regret as the cumulative fold difference in expected growth between a hypothetical population using the reference strategy ***q*** and the evolving population using ***p***^*t*^:

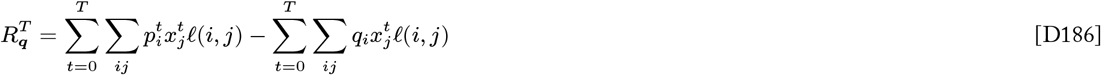

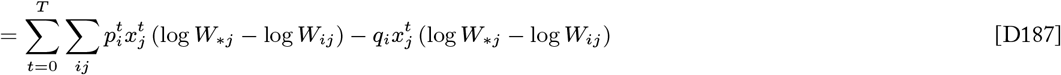

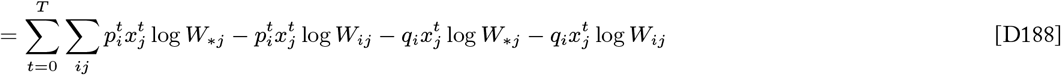

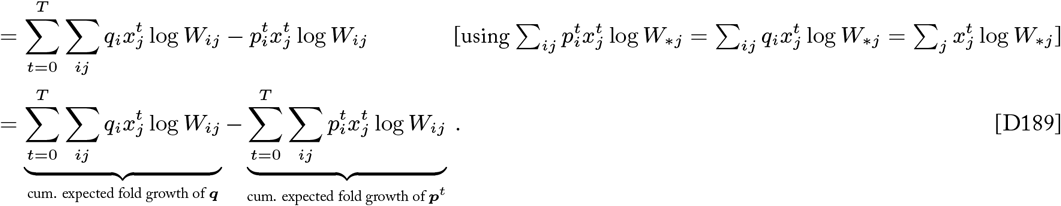

If we consider a lineage to be the collection of individuals derived from an initial population, regret can be equivalently expressed in terms of the population’s total expected lineage size relative to the size of the reference lineage with composition ***q***:

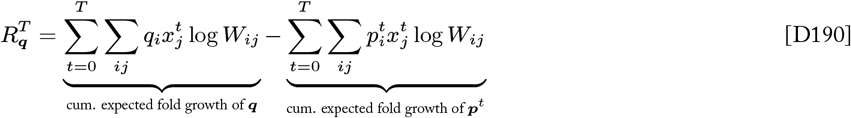

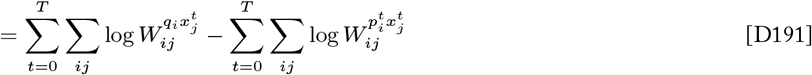

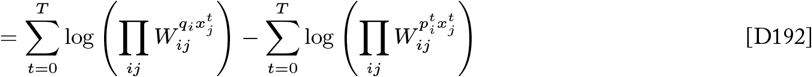

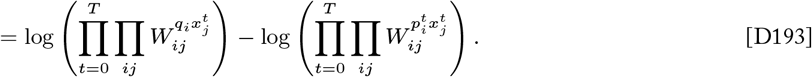

Here we recognize 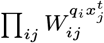 in Equation D192 as the geometric mean fitness over composition ***q*** and environment ***x***^*t*^. Then the quantity 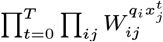 in Equation D193 gives the total expected geometric growth of a lineage with composition ***q*** over T generations. Therefore, we can see that regret can be re-expressed as the difference in the log (absolute) growth of the reference population with composition ***q*** and that of the evolving population using ***p***^*t*^.

If we let the reproductive output of a lineage be denoted as

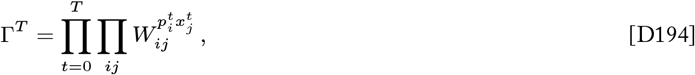

then we can simplify the above expression for regret:

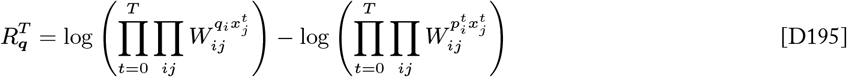

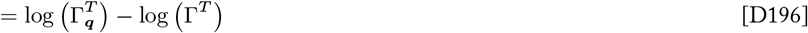

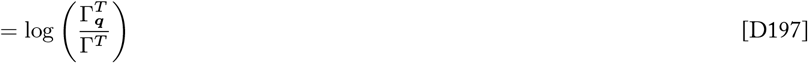

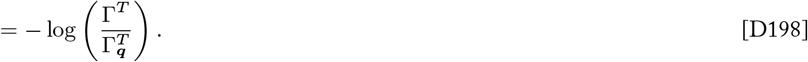

Here we see that regret can be equivalently expressed as the negative log ratio of the population’s total lineage size Γ^*T*^ to the size of the reference lineage 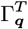 with fixed composition ***q***.

We may choose to consider the reference lineage that has the optimal cumulative growth 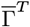 for a particular sequence of environments (two notions of optimality are considered in Appendix D.2.3). Then, just as relative fitness can refer to the short-term reproductive output of a type relative to the maximum output of other types, we can interpret the ratio 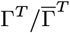 as the **relative lineage fitness**: the cumulative growth of the evolving lineage relative to that of the optimal lineage for the given sequence of environments. In this case, regret is equal to the negative log relative lineage fitness, and minimizing regret coincides with maximizing relative lineage fitness.

##### D.2.3 ESS regret and empirical regret

Regret measures the performance of a learning process in terms of the cumulative loss of the learner’s strategies relative to that of a fixed reference strategy. Typically the reference strategy is taken to be a strategy that is optimal in some sense. For a population that learns by natural selection, it is useful to distinguish two classes of learning problems that lead to different definitions of strategy optimality and thus regret.

###### ESS regret measures the cost of selection with respect to fixed learning problems

First we consider a definition of regret that pertains to a population that evolves by natural selection in response to a fixed learning problem (Appendix C.3). In this context, selection moves the population toward a stationary evolutionarily stable state (ESS). By definition, the evolutionarily stable state is an local optima where the population has higher expected fitness than all other neighboring type frequency distributions for the given environment (Maynard Smith 1982). Natural selection is a process by which the population learns this optimal composition. To evaluate the cost of natural selection as a learning process in this setting, it is natural to measure how the cumulative fitness loss of the evolving population compares to the cumulative fitness loss that would have have been achieved if the population used the optimal ESS composition all along in the same sequence of environments. This measures defines what we refer to as the *ESS regret*, or simply *regret* in the paper.

**Definition**. *For a sequence of environments* ***x***^0^, …, ***x***^*T*^ *that constitutes a fixed learning problem with a stationary evolutionarily stable state* 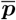, *the regret of a learning process that generates a sequence of type frequency distributions* ***p***^0^, …, ***p***^*T*^ *measured with respect to* 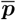 *is referred to as the* **ESS regret**

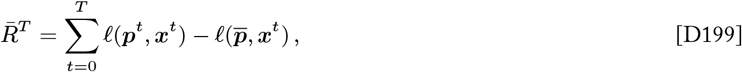

*where the notation* 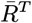 *is used as a shorthand for* 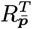 *for typesetting clarity*.

###### Empirical regret measures the cost of selection in generalized settings

ESS regret is a natural measure that captures the cost of selection for the core problem of interest for evolving populations—finding evolutionarily stable states—however it is only applicable to fixed learning problems characterized by stationary evolutionarily stable states. In general, a population may face a sequence of environments for which the accessible evolutionarily stable state shifts over time. In addition, it may not always be reasonable to assume that the evolutionarily stable state is known and usable as a reference distribution for the measurement of regret. For example, an experimentalist may not have sufficient knowledge of the empirical Population versus Environment game they are studying to be able to identify the stable state(s) that characterize the learning problem faced by their population.

In computational learning theory, regret is conventionally measured with respect to the fixed strategy that would have had the minimum load possible against the observed sequence of environments in hindsight.

**Definition**. *For any sequence of environments* ***x***^0^, …, ***x***^*T*^, *the regret of a learning process that generates a sequence of type frequency distributions* ***p***^0^, …, ***p***^*T*^ *measured with respect to the fixed strategy* 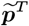 *that gives the minimum loss over the sequence of environments observed in hindsight is referred to as the* **empirical regret**

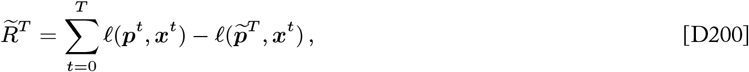

*where the* ***empirically optimal strategy*** 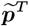 *is defined as*

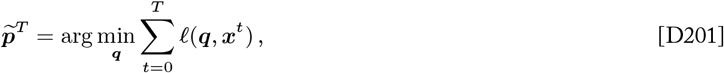

*and where the notation* 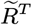 *is used as a shorthand for* 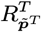 *for typesetting clarity*.

This version of regret is defined solely with respect to the empirical sequence of environments that is observed by the learner and does not rely on any defining characteristics of the sequence of environments or the learning process itself. Therefore this definition of regret is fully general and can be used to measure the cost of any learning process in any online learning setting. Given the generality of this definition, bounds that are established for the empirical regret hold for all games and for all possible environmental sequences and therefore represent worst-case regret guarantees for a learning process.

#### D.3 Regret bounds for natural selection

Now that we have established regret as a meaningful measure of the cost of a learning process, we would like to establish bounds on the cost of selection. Following in the footsteps of foundational works in online learning theory (Kivinen and Warmuth 1995, Freund and Schapire 1999, Cesa-Bianchi and Lugosi 2006), our basic method for deriving bounds will be a kind of amortized analysis that tracks the progression of the learning process using a potential function. We will consider some potential function of frequency vectors *ϕ*(***p***) that is assumed to be non-negative and bounded from above

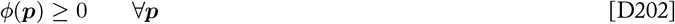

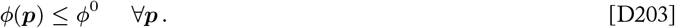

The difference in potential *ϕ*(***p***^*t*^) − *ϕ*(***p***^*t*+1^) (or 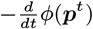 for a continuous process) is one way of describing the progress made by the learning process at time *t*. If we can prove that

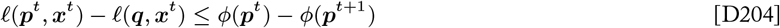

for all possible updates made by the learning process, then by summing the progression over time we will obtain

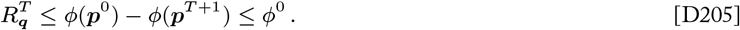

In other words, if the instantaneous regret is bounded by the progress in that step, and the total amount of progress possible is bounded, then we obtain a bound for the total regret.

The particular potential function that we will use in this analysis is the aptly named *potential information*

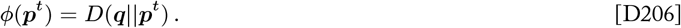

The potential information is defined as the KL divergence of a reference distribution ***q*** from the learner’s current distribution ***p***^*t*^, which can be interpreted as the amount of information the learner stands to gain by updating to the distribution ***q***. This choice of potential function is not arbitrary but rather follows from the information geometry that underlies the dynamics of selection. The potential information is the dynamical potential for which selection is a gradient flow—the surface upon which selection moves the population along the steepest path (Harper 2009a, Harper and Fryer 2015). The tight connection between potential information and regret that will be developed in the following sections reinforces the fundamental role of information in understanding the dynamics and performance of natural selection.

The following sections work through the proof method outlined above to establish regret bounds for replicator dynamics.

##### D.3.1 Regret bound for fixed learning problems

First we consider a population evolving in a sequence of environments characterized by a stationary evolutionarily stable state (i.e., “fixed learning problems”). In this context, we can establish tight, finite, and converging upper and lower bounds on regret, as anticipated by Kimura. We do so first for continuous replicator dynamics, where dynamical and geometrical intuitions can be developed most naturally, before proving the analogous bounds for discrete replicator dynamics that are the focus of the paper.

###### Continuous-time replicator dynamics

The main result of this section is the following theorem, which places an upper bound on the total regret of continuous-time replicator dynamics with respect to a stationary evolutionarily stable state.

###### Theorem D1.

*For any game matrix* ***G*** *and for any sequence of distributions of environmental conditions* ***x***^0^, …, ***x***^*T*^ *such that the population’s initial type distribution* ***p***^0^ *is in the basin of attraction of an evolutionary stable state* 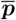 *that remains stationary for all t* ∈ [0, *T*], *the total regret* 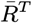 *with respect to* 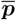 *of the trajectory of type distributions* ***p***^0^, …, ***p***^*T*^ *generated by continuous-time replicator dynamics is bounded from above by the initial potential information*

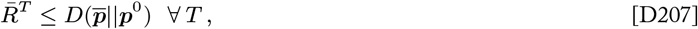

*with equality as T* → ∞

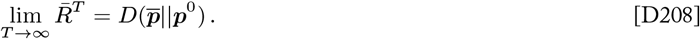

The proof of this theorem follows the logic outlined above (Equation D202 - Equation D205). First we relate the instantaneous regret to the instantaneous change in potential information. Then we establish the potential information 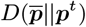 as a non-negative and bounded potential function. The proof then follows from these propositions.

We begin by deriving a fitness-based expression for the change in potential information under replicator dynamics. This result will be handy for later steps.

###### Lemma D1.

*Let* ***p***^*t*^ *be the type frequency distribution of a population that evolves according to continuous-time replicator dynamics across a distribution of environmental conditions* ***x***^*t*^ *at time t. Then the derivative of the potential information between* ***p***^*t*^ *and an arbitrary type frequency distribution* ***q*** *can be expressed*

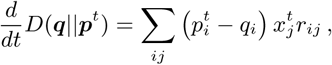

*where r*_*ij*_ *is the growth rate of type i in environmental condition j*.

*Proof*.

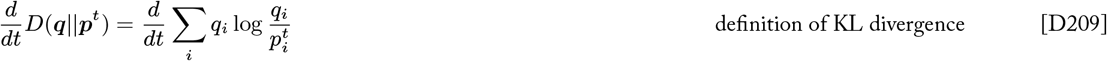

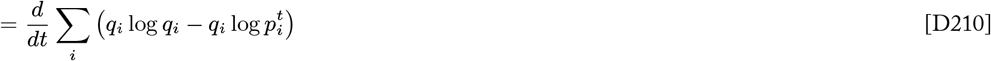

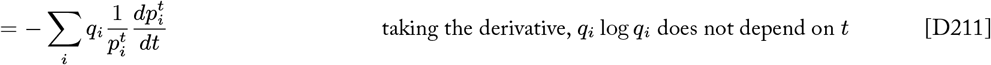

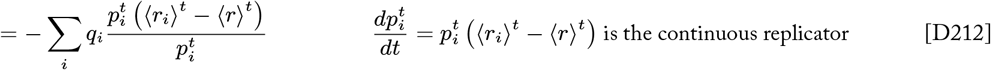

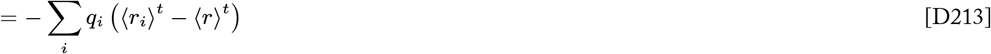

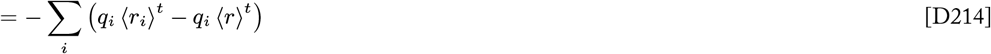

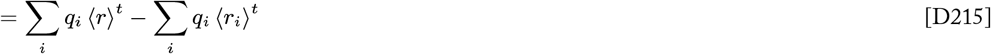

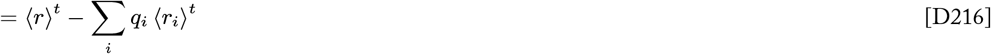

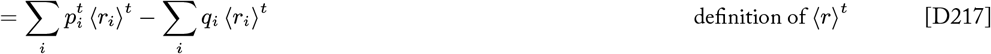

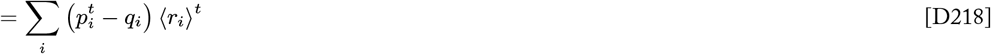

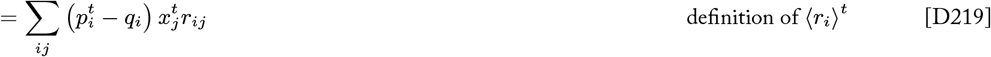

□

###### Proposition D1.

*Let* ***p***^*t*^ *be the type frequency distribution of a population that evolves according to continuous-time replicator dynamics across a distribution of environmental conditions* ***x***^*t*^ *at time t. Then the derivative of the potential information between* ***p***^*t*^ *and an arbitrary type frequency distribution* ***q*** *can be expressed*

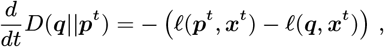

*where* (*ℓ*(***p***^*t*^, ***x***^*t*^) − *ℓ*(***q, x***^*t*^)) *is the instantaneous regret with respect to* ***q***.

*Proof*. We derive the following expression for the instantaneous regret in terms of fitness:

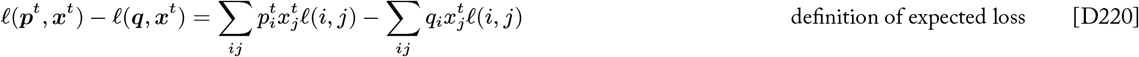

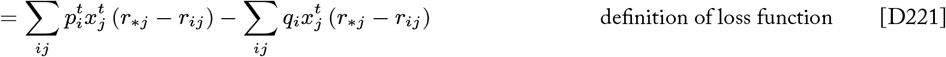

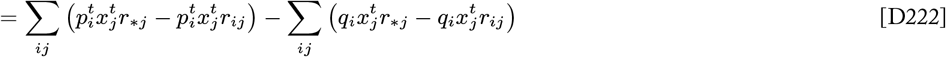

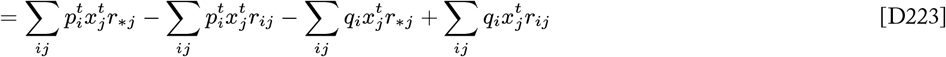

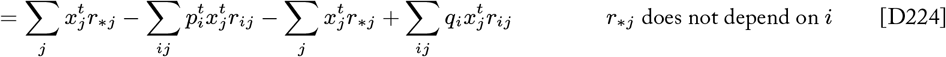

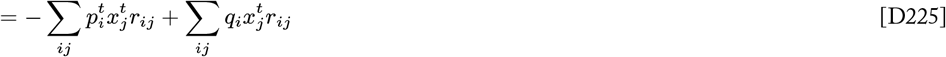

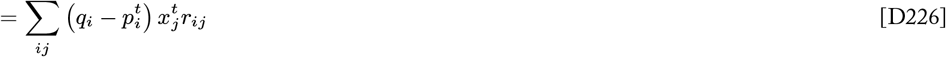

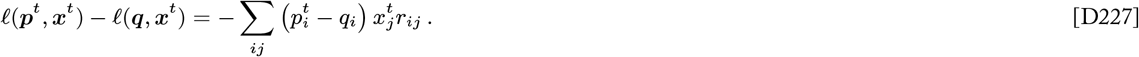

This expression for instantaneous regret is equal to the negation of the expression for the derivative of potential information that was derived as Lemma D1. Therefore we have that

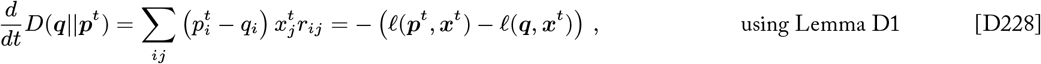

as was to be shown. □

###### Proposition D2.

*If* 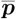 *is a stationary evolutionarily stable state, then the potential information* 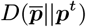 *is a Lyapunov function that satisfies for all* ***p***^*t*^ *generated by continuous-time replicator dynamics in a neighborhood* 𝒬 *of* 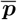

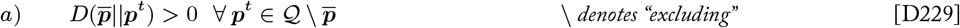

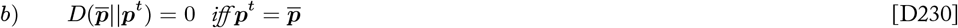

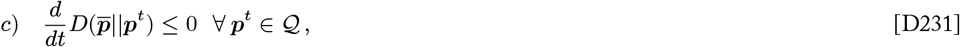

*and* 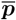 *is an asymptotically stable equilibrium point of the system at time t*.

*Proof*. Conditions (a) and (b) follow, respectively, from the facts that KL divergence is always non-negative and is equal to zero if and only if the two distributions are equal (Cover and Thomas 2006). To prove condition (c) we observe that ***p***^*t*^ is in the neighborhood of an evolutionarily stable state 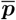 by the premise of the proposition and the definition of a stationary ESS, and thus we have

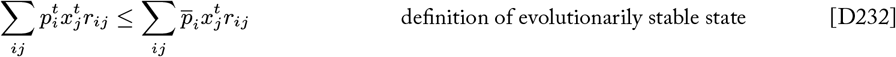

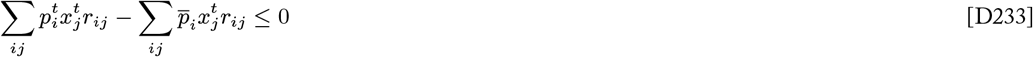

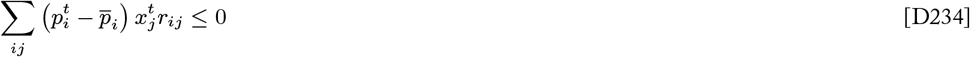

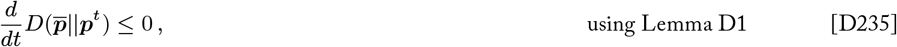

as was to be shown. Therefore the potential information 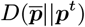 satisfies the conditions of a Lyapunov function for the continuous-time replicator dynamics, the existence of which implies that 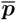 is an *asymptotically stable equilibrium point* by the Lyapunov Asymptotic Stability Theorem. □

###### Corollary D2.1.

*If* 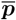 *is a stationary evolutionarily stable state, then the potential information* 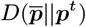 *is bounded from above by its initial value*

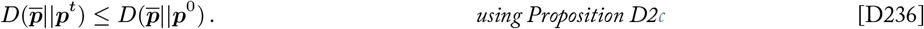

###### Corollary D2.2.

*If* 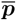 *is a stationary evolutionarily stable state, then its basin of attraction* 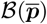 *is the largest positively invariant set for which* 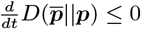 *under continuous-time replicator dynamics for all* 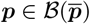.

###### Proof of Theorem D1

Proposition D1 relates the instantaneous change in potential information to the instantaneous regret. Considering the regret with respect to a stationary evolutionarily stable state 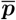, we have that

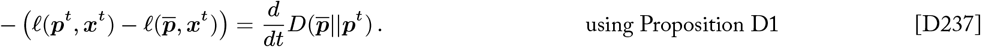

Integrating over a period of selection of duration *T* we obtain

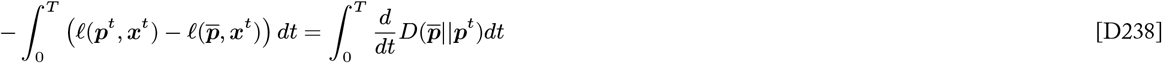

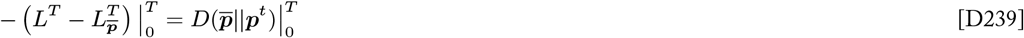

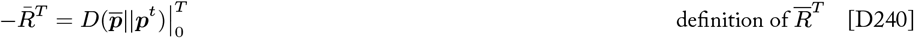

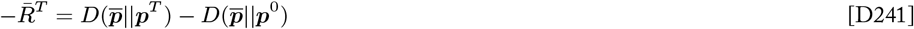

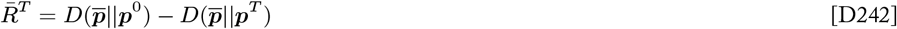

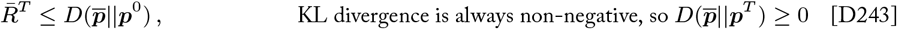

which proves that the initial potential information is an upper bound on the total regret. The stationary evolutionarily stable state 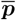 is an asymptotically stable equilibrium point, and the population’s initial type distribution is within its basin of attraction as stated in the premise of the theorem. Therefore, 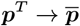 as *T* → ∞, and we have that

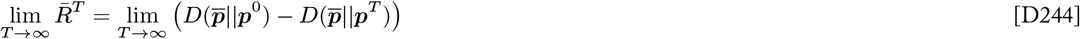

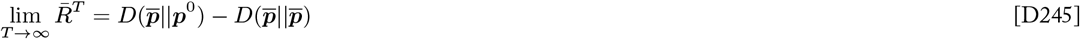

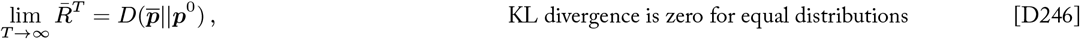

which proves that the upper bound is tight, as the total regret converges on the initial potential information as *T* → ∞. □

###### Discrete-time replicator dynamics

The main result of this section is the following theorem, which places an upper bound on the total regret of discrete-time replicator dynamics with respect to a stationary evolutionarily stable state.

###### Theorem 1.

*For any game matrix* ***G*** *and for any sequence of distributions of environmental conditions* ***x***^0^, …, ***x***^*T*^ *such that the population’s initial type distribution* ***p***^0^ *is in the basin of attraction of an evolutionary stable state* 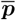 *that remains stationary for all t* ∈ [0, *T*], *the total regret* 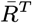 *with respect to* 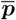 *of the trajectory of type distributions* ***p***^0^, …, ***p***^*T*^ *generated by discrete-time replicator dynamics is bounded from above by the initial potential information*

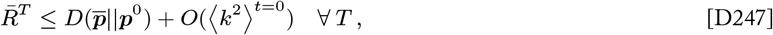

*with equality as T* → ∞

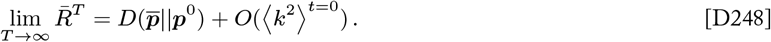

*where k*_*ij*_ *is the Wrightian selection coefficient of type i in state j (Appendix A.1.1) and* 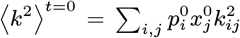 *is the mean squared selection coefficient across types and conditions at time t* = 0.

This result is nearly identical to the corresponding bound for continuous-time replicator dynamics given in Theorem D1, save for a “gap” term related to the strength of selection. This “gap” can be interpreted as a small amount of additional regret that may be incurred due to imprecisions in discrete replicator dynamics’ approximation of the continuous path of convergence to the evolutionarily stable state. When the strength is selection is high, discrete updates may carry the population further in tangential directions, and the expected imprecision of the discrete-time trajectory is greater. In the limit of weak selection, where most individuals have similar relative fitnesses and thus the selection coefficients *k*_*ij*_ are small, this imprecision is negligible and the discrete-time bound approaches equality with that of continuous-time replicator dynamics.

The proof of this theorem proceeds similarly to the proof of the bound for continuous replicator dynamics in the previous section. First we relate the instantaneous regret to the instantaneous change in potential information. Then we establish the potential information 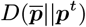 as a non-negative and bounded potential function. The proof then follows from these propositions.

Discrete-time replicator dynamics are a special case of the Multiplicative Weight Updating (MWU) algorithm where the learning rate *η* = 1 and a particular fitness-based loss function is used (Appendix C.2.4). We begin by working through intermediate results for the general class of MWU algorithms where the learning rate *η* and the loss function *ℓ*(*i, j*) are left unspecified. This provides some insights about how the learning rate, which modulates the learning algorithm’s balance of loss minimization and diversity maintenance, affects the relationship between potential information and regret. Ultimately, Theorem 1 and the associated propositions are stated in terms of the replicator dynamics case where *η* = 1 and *ℓ*(*i, j*) = − log *w*_*ij*_.

###### Proposition D3.

*Let* ***p***^*t*^ *be the type frequency distribution of a population that evolves according to discrete-time replicator dynamics over a distribution of environmental conditions* ***x***^*t*^ *at time t. Then the change in the potential information between* ***p***^*t*^ *and an arbitrary type frequency distribution* ***q*** *can be expressed*

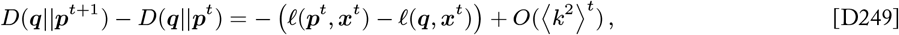

*where k*_*ij*_ *is the Wrightian selection coefficient of type i in condition j (Appendix A.1.1) and* 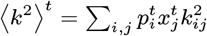 *is the expected squared selection coefficient across types and conditions at time t. We refer to the quantity (ℓ*(***p***^*t*^, ***x***^*t*^) − *ℓ*(***q, x***^*t*^)) *as the instantaneous regret with respect to* ***q***.

*Proof*. We set out to evaluate an expression for the difference in the potential information across one update of the learning process:

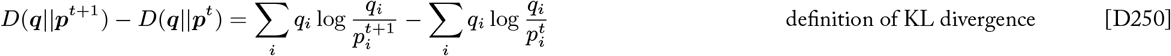

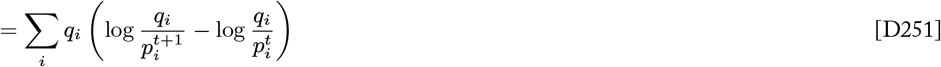

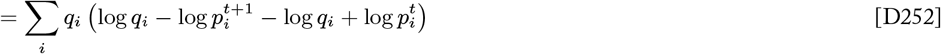

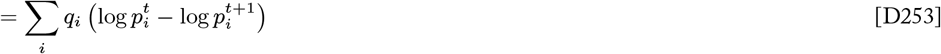

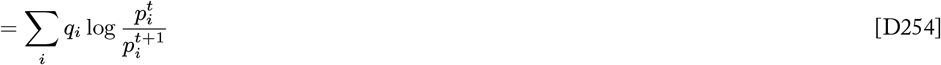

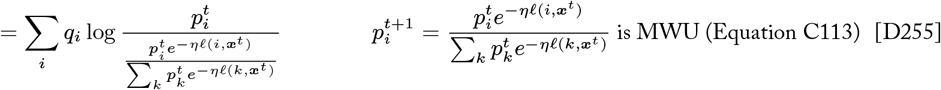

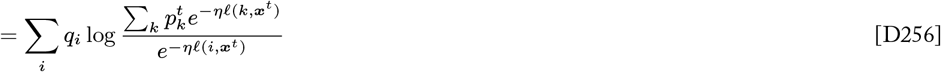

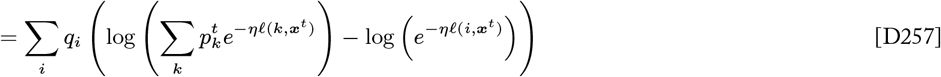

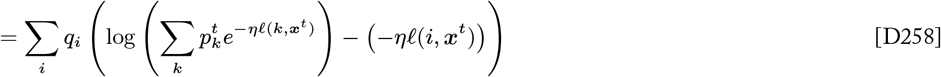

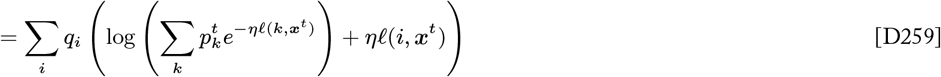

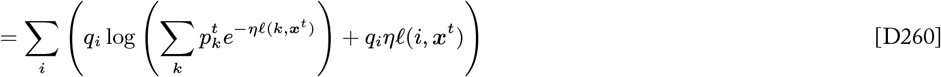

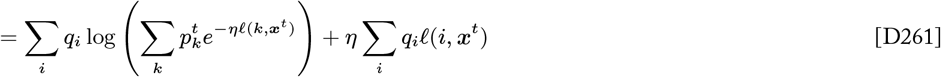

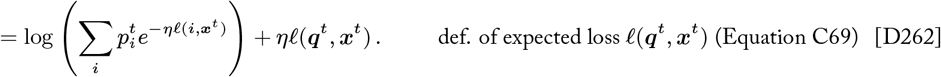

In order to simplify the first term further, we can invoke Jensen’s inequality. Jensen’s inequality states that for a *concave* function *g*(*u*)

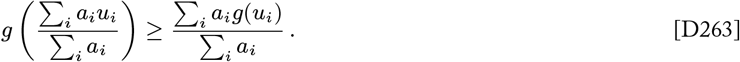

This can be equivalently restated as

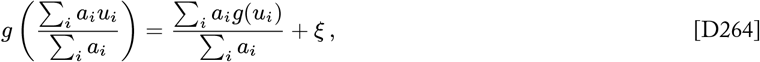

where *ξ* is the value of the *Jensen gap*—the difference between the left- and right-hand sides of Jensen’s inequality

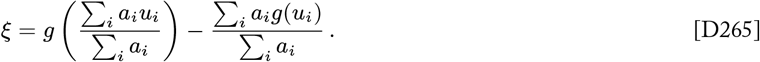

Using Equation D264, we can express the first term in Equation D262 as

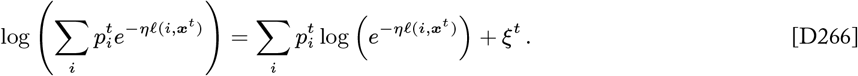

Picking up where we left off in the derivation of an expression for the change in potential information, we have

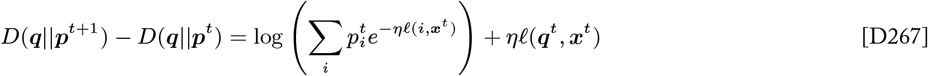

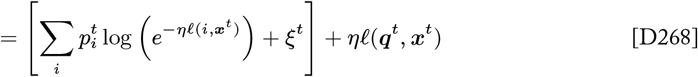

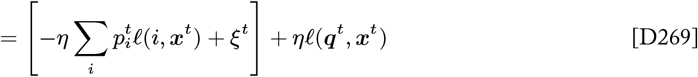

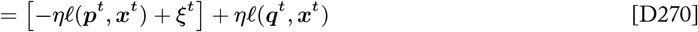

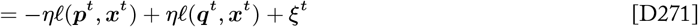

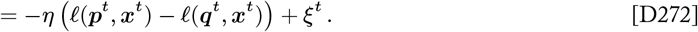

At this point we see that the change in potential information across one update of the discrete-time learning process is proportional to (equal to for *η* = 1) the inverse of the instantaneous regret experienced in the most recent time step, plus a value equal to the Jensen gap *ξ*^*t*^ that arose in our derivation. From here, we would like to evaluate the size of this Jensen gap term. Solving for *ξ*^*t*^ using Equation D265 and Equation D266, we have

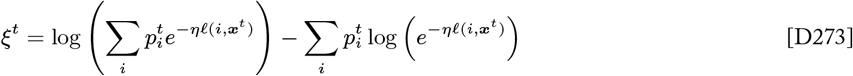

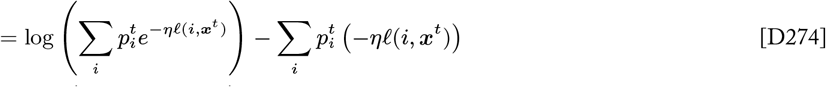

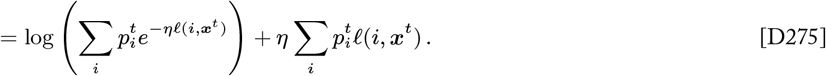

In general, the size of this Jensen gap depends on the choice of loss function *ℓ*(*i, j*), but we can evaluate the size of this gap further by considering the particular loss function *ℓ*(*i, j*) = − log *w*_*ij*_ and learning rate *η* = 1 that are implicit in the replicator dynamics instance of MWU:

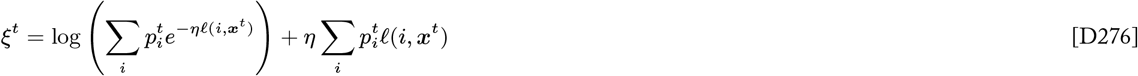

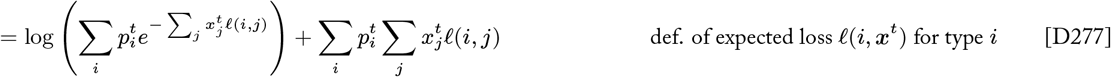

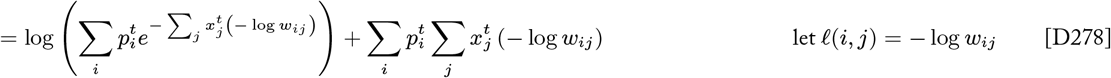

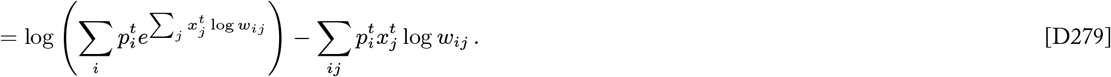

We can simplify the first term in this gap expression by making use of Jensen’s inequality (Equation D263) again. In particular, Jensen’s inequality tells us that 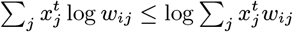, which implies that 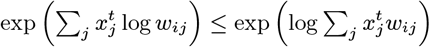.

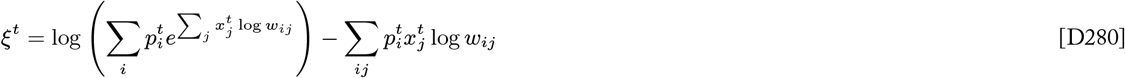

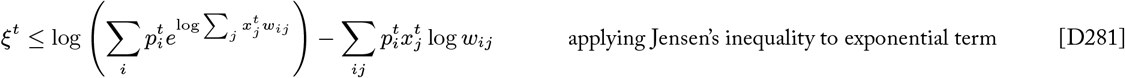

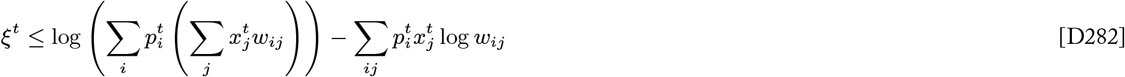

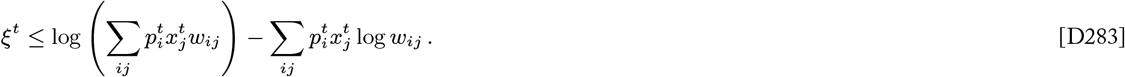

We have arrived at an inequality that bounds the Jensen gap in terms of relative fitnesses, but the scale of this bound is not particularly intuitive. We can derive a simpler expression for the magnitude of the Jensen gap by expressing relative fitnesses in terms of Wrightian selection coefficients: *w*_*ij*_ = 1 − *k*_*ij*_ (Appendix A.1.1).

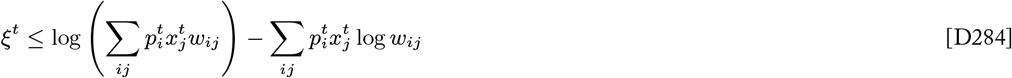

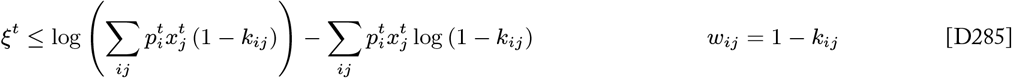

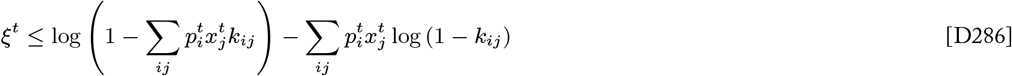

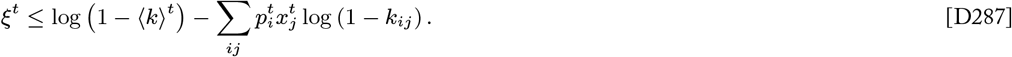

Applying the Taylor expansion 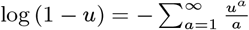 to the logarithms in both terms and simplifying

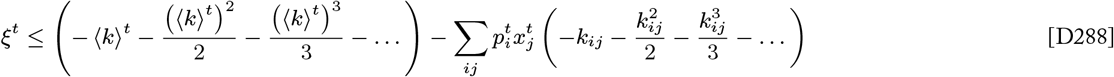

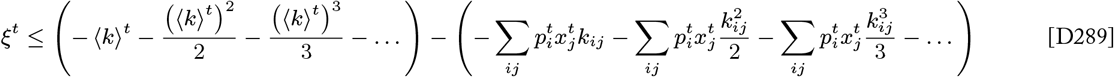

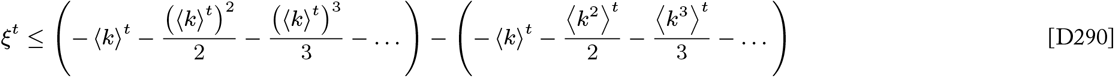

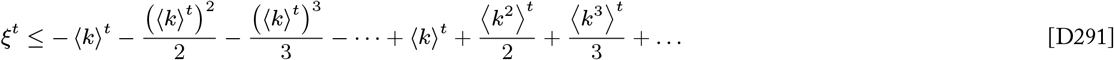

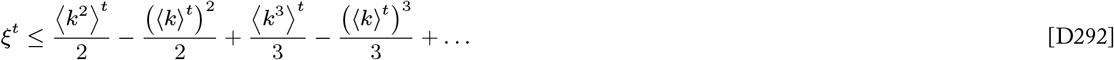

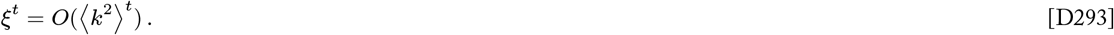

We find that the Jensen gap is bounded by an infinite series involving the expected values of the population’s selection coefficients across types and conditions. Turning to Jensen’s inequality yet again gives 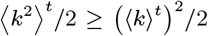, which tells us that this series is dominated by the first term 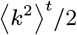. Thus we establish that the Jensen gap term *ξ*^*t*^ is on the order 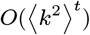 or less.

Therefore, in the case of discrete-time replicator dynamics where *η* = 1 and *ℓ*(*i, j*) = − log *w*_*ij*_, the change in potential information across one update is given by

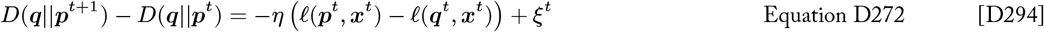

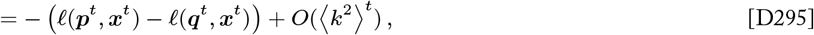

as was to be shown. □

###### Proof of Theorem 1.

Proposition D3 relates the instantaneous change in potential information to the instantaneous regret for discrete-time replicator dynamics. Considering the regret with respect to a stationary evolutionarily stable state 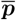, we have that

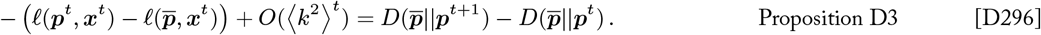

Summing over a period of selection of duration *T* we obtain

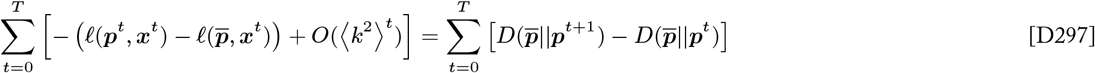

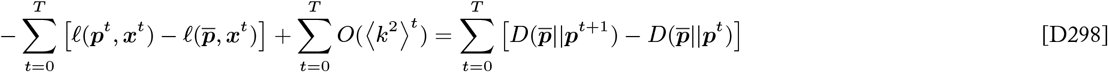

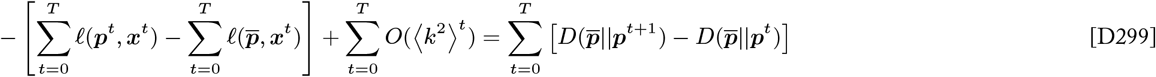

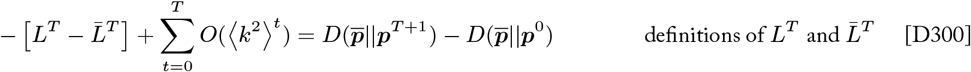

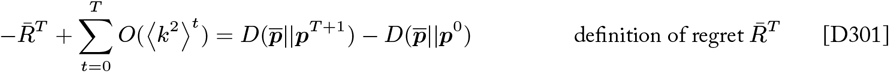

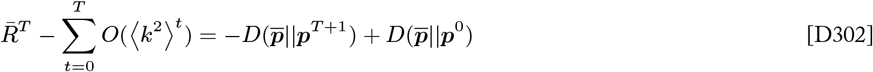

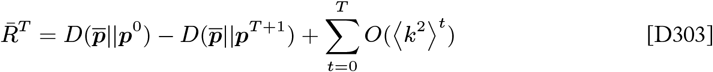

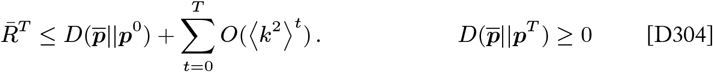

We know that 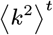 is decreasing monotonically when selection is approaching a stationary evolutionarily stable state, so 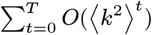 is sublinear in *T*. Therefore we can restate the bound on regret

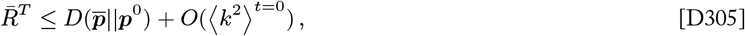

as was to be shown. The stationary evolutionarily stable state 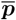 is an asymptotically stable equilibrium point, and the population’s initial type distribution is within its basin of attraction as stated in the premise of the theorem. Therefore, 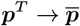 as *T* → ∞, and we have that

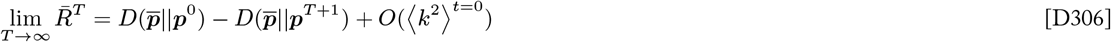

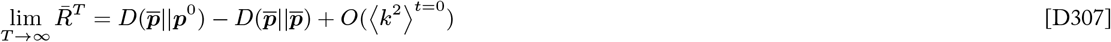

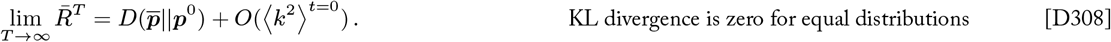

as was to be shown. □

When most individuals have similar relative fitnesses and thus the selection coefficients *k*_*ij*_ are small, the Jensen gap is also small. In the limit of weak selection where the selection coefficients *k*_*ij*_ are nearly zero for all types *i* in all conditions *j*, the Jensen gap is vanishing and the results stated in Proposition D3 and Theorem 1 are equal to the corresponding results for continuous-time replicator dynamics (Proposition D1 and Theorem D1).

###### Corollary 1.1.

*For any game matrix* ***G*** *and for any sequence of distributions of environmental conditions* ***x***^0^, …, ***x***^*T*^ *such that the population’s initial type distribution* ***p***^0^ *is in the basin of attraction of an evolutionary stable state* 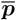 *that remains stationary for all t* ∈ [0, *T*], *the total regret* 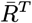 *with respect to* 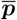 *of the trajectory of type distributions* ***p***^0^, …, ***p***^*T*^ *generated by discrete-time replicator dynamics <u>in the limit of weak selection</u> is bounded from above by the initial potential information*

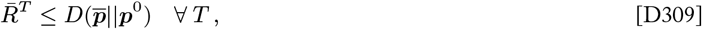

*with equality as T* → ∞

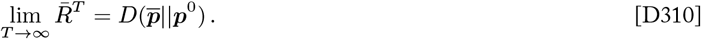

##### D.3.2 Regret bounds for variable learning problems

Now we expand our consideration to the general case where no assumptions are made about the sequence of environments or stable states. The upper bounds established in this setting provide guarantees for the maximum possible regret a population can experience in any setting, including arbitrary or adversarial sequences of environments. Therefore these results describe the worst-case performance of natural selection as a learning process.

Discrete-time replicator dynamics are a special case of the Multiplicative Weights Updating (MWU) algorithm. MWU has been widely studied in computational learning theory and related fields, and various bounds have been established for this class of algorithms in different contexts. In some cases—such as when certain assumptions can be made about the learning problem, when the loss function has certain properties, or when the learning rate can be tuned in response to the learning problem—versions of MWU achieve very tight bounds on regret. In fact, it can be proven that no other learning process can possibly achieve tighter bound than some special instances of MWU (Freund and Schapire 1999, Cesa-Bianchi and Lugosi 2006). However, the instance of MWU that is equivalent to replicator dynamics does not fall into any of these special categories. Nevertheless, general regret bounds for MWU that hold for any learning rate and any loss function can be directly applied to replicator dynamics.

The main result of this section is the following theorem, which places an upper bound on the total empirical regret of discrete-time replicator dynamics in the general case.

###### Theorem 3.

*For any game matrix* ***G*** *and for any sequence of distributions of environmental conditions* ***x***^0^, …, ***x***^*T*^, *the total empirical regret* 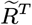 *with respect to* 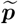 *of the trajectory of type distributions* ***p***^0^, …, ***p***^***T***^ *generated by discrete-time replicator dynamics is bounded from above by*

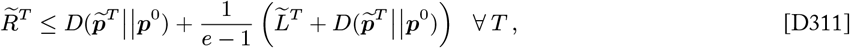

*where* 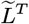 *is the cumulative loss of the empirically optimal strategy* 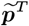, *and where e denotes Euler’s number*.

This result follows from Freund and Schapire (1999), which in turn draws on the amortized analysis introduced by Kivinen and Warmuth (1995). We recite an adapted version of their proof here, which follows similar logic as those in previous sections (e.g., Equation D202 - Equation D205). The heart of the proof is the following proposition, which bounds the cumulative loss of the learner in terms of the loss of the empirically optimal strategy and the initial potential information (this proposition is presented as Proposition 5 in the main text).

###### Proposition D4.

*For any game matrix* ***G*** *and for any sequence of distributions of environmental conditions* ***x***^0^, …, ***x***^*T*^, *the total cumulative loss L*^*T*^ *of the trajectory of type distributions* ***p***^0^, …, ***p***^*T*^ *generated by discrete-time replicator dynamics is bounded from above by*

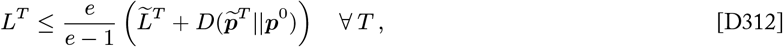

*where* 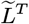 *is the cumulative loss of the empirically optimal strategy* 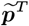, *and where e denotes Euler’s number*.

*Proof*. As with proofs in the previous sections, we begin by seeking an expression for the change in the potential information in one update in terms of single round losses:

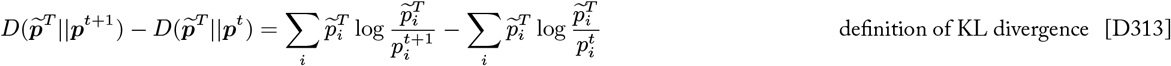

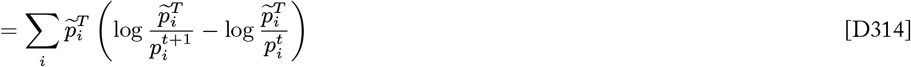

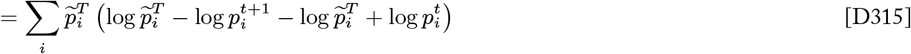

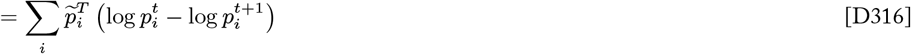

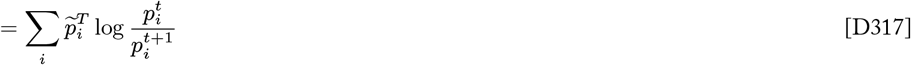

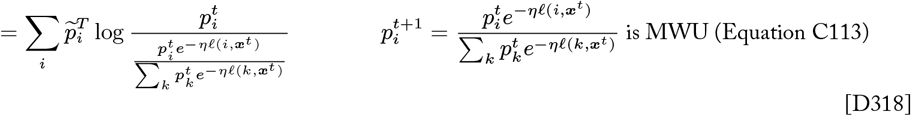

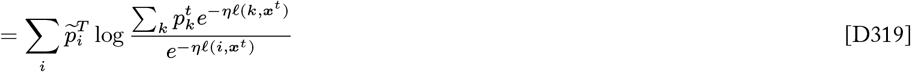

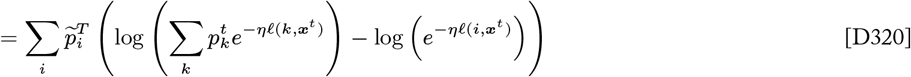

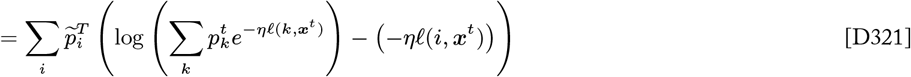

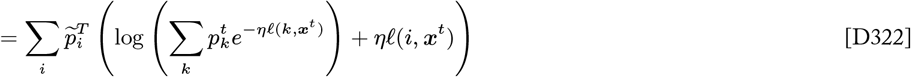

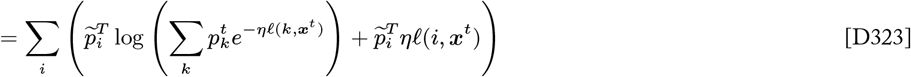

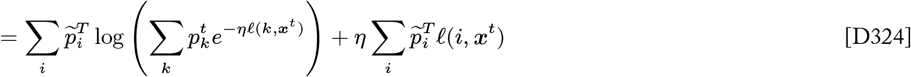

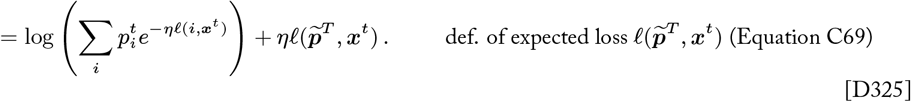

Here we make use of the fact that, by convexity, *β*^*u*^ ≤ 1 − (1 − *β*)*u* for *β* ≥ 0 and *u* ∈ [0, 1]^*^. Thus if we let *β* = *e*^−*η*^ and *u* = *ℓ*(*i*, ***x***^*t*^), then 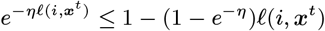. Returning to our derivation at Equation D325, we have

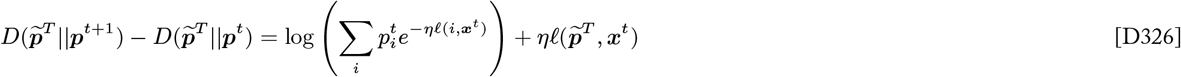

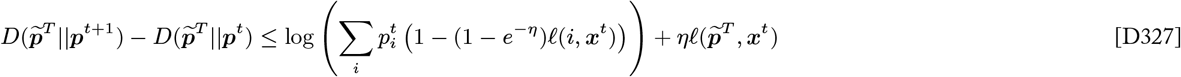

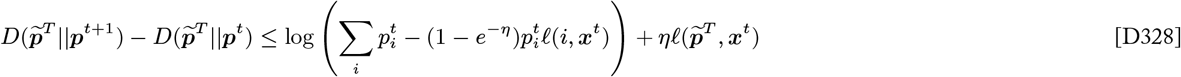

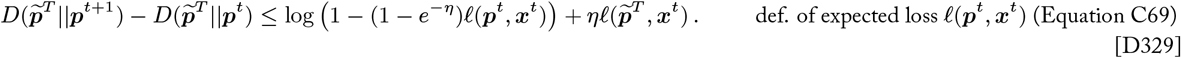

Now we simplify the first loss term using the fact that log (1 − *u*) ≤ −*u* for any *u <* 1. If we let *u* = 1 − (1 − *e*^−*η*^)*ℓ*(***p***^*t*^, ***x***^*t*^), then we have

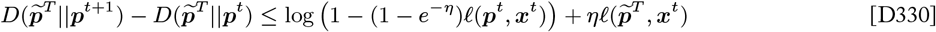

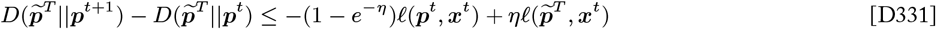

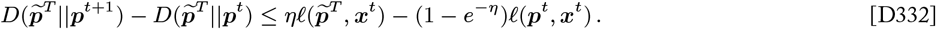

By summing this inequality over a period of duration *T* and rearranging terms we obtain

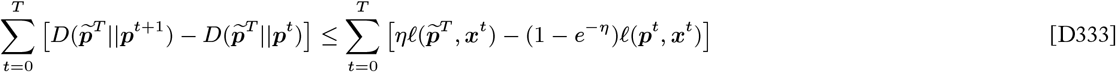

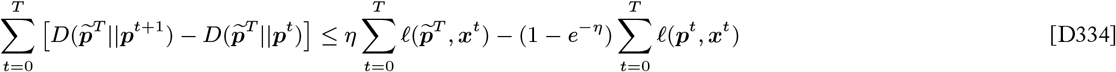

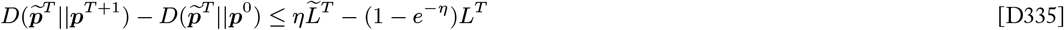

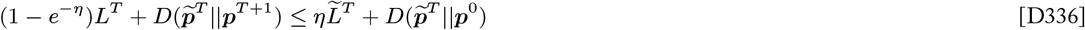

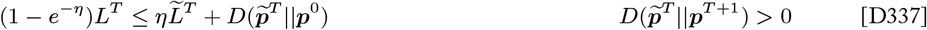

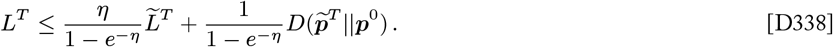

The learning rate implicit in the equivalence between replicator dynamics and MWU is *η* = 1. Plugging in this value for *η* we have

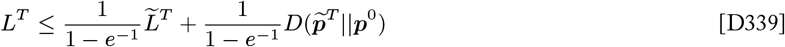

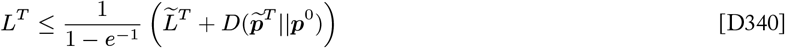

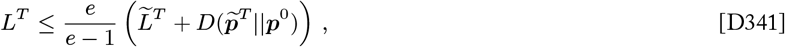

as was to be shown. □

###### Proof of Theorem 3.

The bound on cumulative load given in Proposition D4 can be rearranged to express a bound on empirical regret

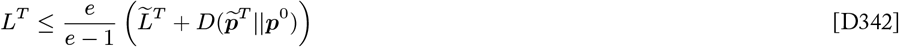

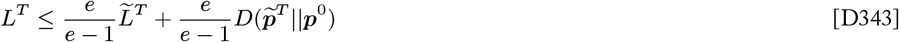

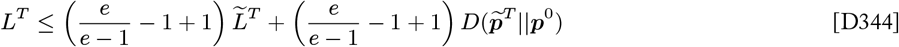

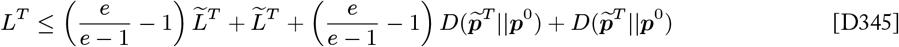

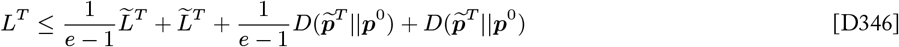

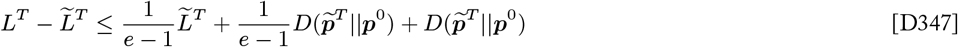

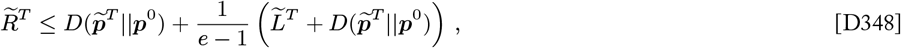

as was to be shown. □

##### D.3.3 Comparing the cost of selection to the cost of other learning algorithms

It is standard practice in computer science to evaluate the relative performance of algorithms in terms of their worst-case performance. In particular, the worst-case regret is commonly used for evaluating online learning algorithms. In general, we cannot make guarantees about the kinds of learning problems (e.g., environments) that a learner will face, so worst-case bounds that describe the maximum regret a learner can experience in any scenario are useful benchmarks. Theorem 3 provides a worst-case upper bound on regret for natural selection. This result can be compared to the worst-case regret of alternative algorithms to assess the relative effectiveness of selection as a learning process.

We can draw certain conclusions about the performance of natural selection relative to other algorithms given that selection is a member of the widely studied class of Multiplicative Weights Updating (MWU) algorithms. Recall that MWU algorithms feature a learning rate parameter (*η* ∈ (0, ∞)) that modulates a balance between concentrating weight on types that have incurred low loss in the past against maintaining diversity over types that may perform well in the future (Appendix C.2.3). MWU is more reactive to each round of loss observations when the learning rate is high (*η* → ∞), while MWU favors more stable strategies when the learning rate is low (*η* → 0). Results from MWU analysis show that MWU is optimal when the learning rate can be appropriately tuned over the duration of the learning problem, in that no other online learning algorithm can outperform (i.e., achieve lower long-term regret in the worst case) these instances of MWU (Freund and Schapire 1999, Cesa-Bianchi and Lugosi 2006). The learning rate implicit in natural selection is a fixed *η* = 1, but such results go to show that selection is in a class of algorithms that are generally effective for the learning problems faced by evolving populations.

We can directly evaluate the performance of natural selection relative to other instances of MWU with fixed learning rates. Equation D338 provides an general upper bound on the worst-case cumulative loss (mismatch load) of a learner using MWU in terms of the learning rate *η*. We used this result to evaluate a general regret bound for natural selection where *η* = 1 (Theorem 3), but we can also use this load bound to establish a worst-case regret bound for an arbitrary learning rate.

###### Corollary 3.1.

*For any game matrix* ***G*** *and for any sequence of distributions of environmental conditions* ***x***^0^, …, ***x***^*T*^, *the total empirical regret* 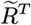 *with respect to* 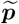 *of the trajectory of type distributions* ***p***^0^, …, ***p***^*T*^*generated by Multiplicative Weights Updating with fixed learning rate η is bounded from above by*

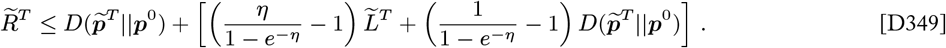

*Proof*.

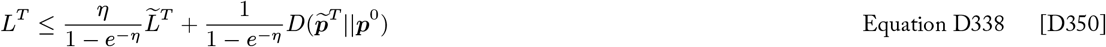

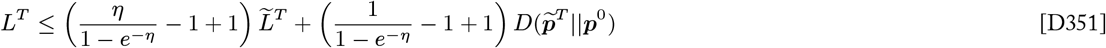

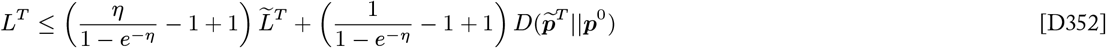

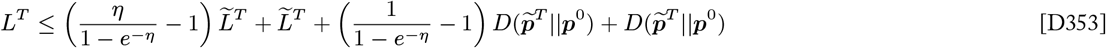

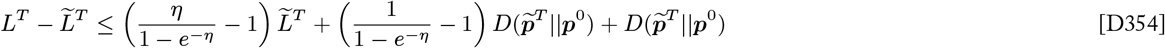

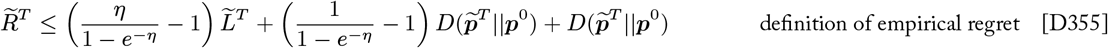

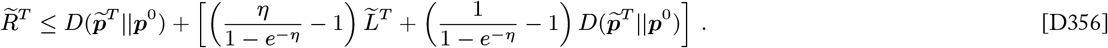

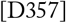

□

Corollary 3.1 shows that the worst-case regret of MWU with a fixed learning rate *η* is bounded by the initial potential information (similar to the bounds we establish for natural selection) plus a second term (in square brackets) that corresponds to additional regret that accumulates when responding to a variable learning problem. We see that this additional regret term is a weighted combination of the cumulative loss of the empirically optimal strategy 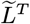 and the initial potential information. When the learning rate is high (i.e., MWU is highly reactive to each round of loss observations) the cumulative loss of the empirically optimal strategy contributes more to the excess regret, but when the learning rate is low (i.e., MWU favors diversity and type frequencies change relatively little in each round) the initial potential information contributes more heavily. Interestingly, the learning rate implicit in natural selection, *η* = 1, balances loss minimization and diversity maintenance exactly equally (Appendix C.2.4).

Figure D6 illustrates how the worst-case empirical regret (given by Corollary 3.1) changes as a function of the MWU learning rate for multiple contexts defined by the cumulative expected loss (mismatch load) of the empirically optimal strategy, which is approximately proportional to the variability of the environment. We see that the worst-case regret is typically minimized for intermediate learning rates that balance loss minimization and diversity maintenance. Learning rates lower than the optima have correspondingly higher worst-case regret because they are slow to move toward the empirically optimal strategy. Likewise, learning rates greater than the optima typically have higher worst-case regret due to the possibility of responding too strongly to short-term variations in the environment. As the variability of the environment and the the expected loss of the optimal strategy increase, the optimal learning rate decreases. On the other hand, when the environment is stable such that the optimal strategy can achieve very low cumulative loss (e.g., the 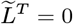 curve in Figure D6), high learning rates that move the population toward the optimal strategy quickly are optimal. The marked points at *η* = 1 in Figure D6 show how the worst-case regret for natural selection compares to the optimal worst-case regret (i.e, curve minima) for MWU with fixed learning rates in each context. We see that selection’s learning rate is suboptimal when the optimal strategy can achieve zero cumulative loss, which reinforces that the gradual nature of selection can be particularly disadvantageous in constant environments such as those considered by Haldane and Kimura. However, when the environment is heterogeneous, stochastic, and/or time-varying such that the optimal strategy has non-trivial cumulative expected loss (mismatch load), then the learning rate of selection is relatively close to the optimum. In variable environments, MWU learning rates that are much higher or lower than that of selection tend to correspond to greater regret in the worst-case.

**Fig. D6.**
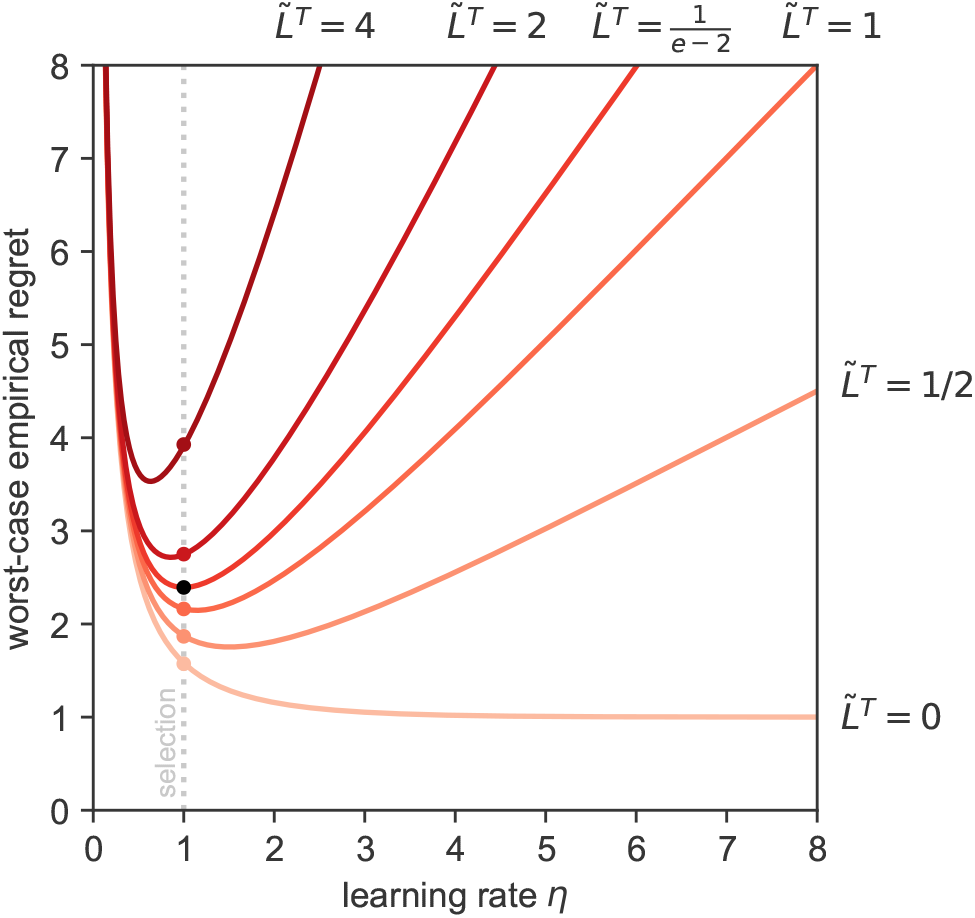
The relationship between learning rate and worst-case empirical regret for Multiplicative Weights Updating. The worst-case regret for a learner using Multiplicative Weights Updating with a fixed learning rate (Corollary 3.1) is plotted across a range of learning rates (*η*). Each curve corresponds to a different context (learning problem) characterized by the cumulative expected loss (mismatch load, 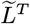) of the optimal empirical strategy, which is approximately proportional to environment variability (indicated by labels near each curve). All contexts assume 1 bit of initial potential information 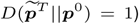. The learning rate that gives the lowest worst-case regret bound decreases as the environment variability and cumulative loss of the optimal strategy increase. Marked points show the worst-case regret for natural selection, where *η* = 1, which can be compared to the optimal regret (minima of curves) for each context. The black marker highlights that natural selection’s learning rate is optimal when the cumulative expected loss of the optimal empirical strategy is equal to 1*/*(*e −* 2) times the initial potential information.

We can use the derivative of the regret bound given in Corollary 3.1 to find the optimal learning rate that gives the lowest worst-case regret for a learning problem with a given optimal load and initial potential information:

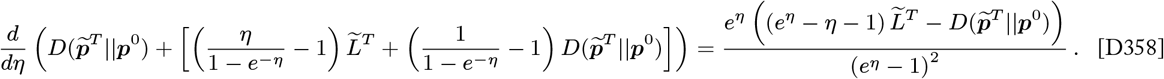

Setting this derivative equal to 0 and solving for *η*:

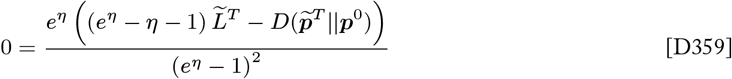

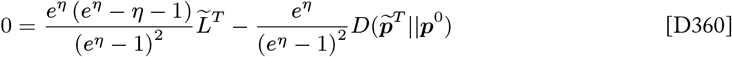

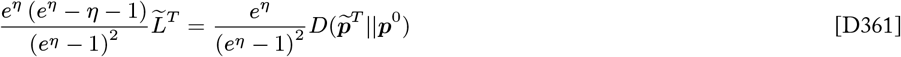

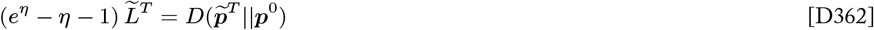

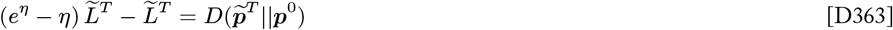

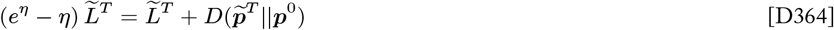

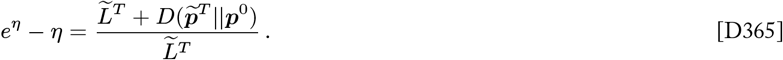

Equation D365 gives an expression for the optimal learning rate *η* in terms of the expected load of the optimal strategy and the initial potential information for a particular learning problem. Evaluating this further to obtain an explicit solution for *η* involves a product logarithm (a.k.a. Lambert *W* function) and does not lend additional intuition.

However, we can evaluate this expression further for *η* = 1 in order to understand when the learning rate implicit in natural selection is optimal:

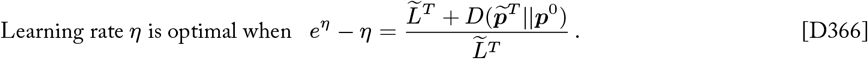

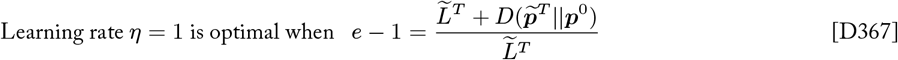

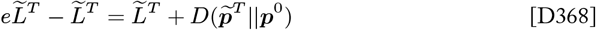

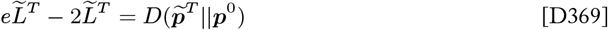

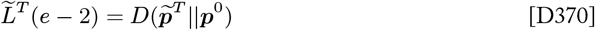

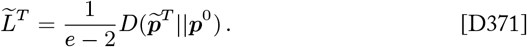

In other words, natural selection’s learning rate is optimal when the cumulative expected loss of the optimal empirical strategy is equal to 1*/*(*e* − 2) ≈ 1.4 times the initial potential information. When the load of the optimal strategy is less than 1*/*(*e* − 2) times the initial potential information, selection’s learning rate is slower than optimal, and when the load of the optimal strategy is greater than this value, selection’s learning rate is higher than the optimum.

#### D.4 The cost of information gain

##### D.4.1 Information gain in a single generation is bounded by look-ahead regret

###### Proposition D5.

*Let* ***p***^*t*^ *be the type frequency distribution of a population that evolves according to discrete-time replicator dynamics over a distribution of environmental conditions* ***x***^*t*^ *at time t. Then the information gain in one selective update, as measured by the divergence between the population’s updated and previous compositions, can be expressed*

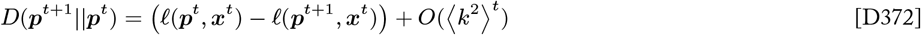

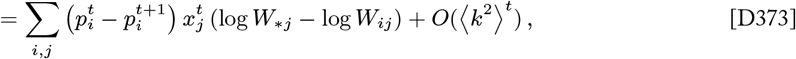

*where k*_*ij*_ *is the Wrightian selection coefficient of type i in condition j (Appendix A.1.1) and* 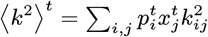 *is the expected squared selection coefficient across types and conditions at time t*.

*Proof*. This result follows directly from Proposition D3 where the reference distribution ***q*** is taken to be the type frequency distribution in the next generation ***p***^*t*+1^.

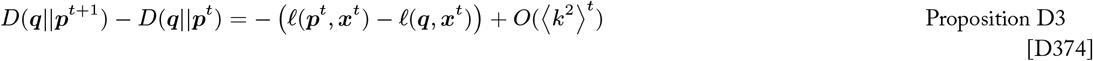

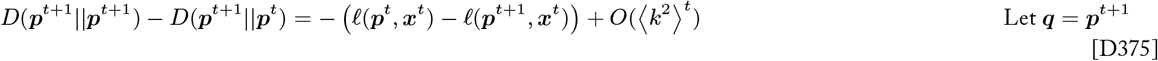

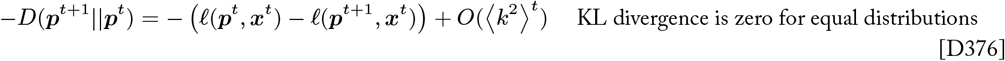

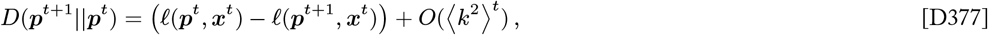

as was to be shown. □

The following corollary that applies the fitness-based loss function implicated by natural selection (Appendix C.2.4) and the limit of weak selection 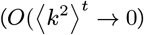 is presented as Proposition 4 in the main text.

###### Corollary D5.1.

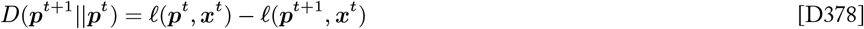

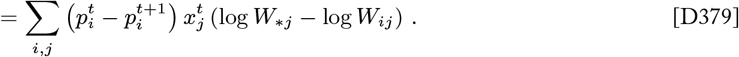

##### D.4.2 Information gain converges on ESS regret

The following theorem establishes that the evolving population’s total information gain converges on the total ESS regret of discrete-time replicator dynamics in contexts characterized by a stationary ESS.

###### Theorem 2.

*For any game matrix* ***G*** *and for any sequence of environmental conditions* ***x***^0^, …, ***x***^*T*^ *such that the population’s initial type distribution* ***p***^0^ *is in the basin of attraction of an <u>evolutionary stable state</u>* 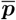 *that remains <u>stationary</u> for all t* ∈ [0, *T*], *the total information gain I*^*T*^ *and the total regret* 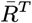 *with respect to* 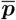 *of the trajectory of type distributions* ***p***^0^, …, ***p***^*T*^ *generated by replicator dynamics in the limit of weak selection both converge on the value of the initial potential information*

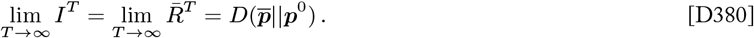

*Proof*. The stationary evolutionarily stable state 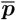 is an asymptotically stable equilibrium point, and the population’s initial type distribution is within its basin of attraction as stated in the premise of the theorem. Therefore, 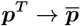 as *T* → ∞, and Corollary 1.1 of Theorem 1 tells us that

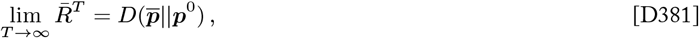

in the limit of weak selection (i.e., *k*_*ij*_ ⋘ 1 for all *i, j*) or continuously overlapping generations (Theorem D1).

The convergence of the information gain *I*^*T*^ on the same value—the initial potential information—follows from the definition of information gain and the convergence of 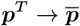.

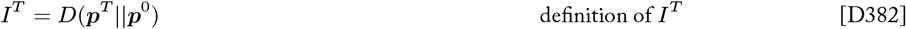

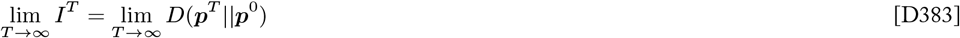

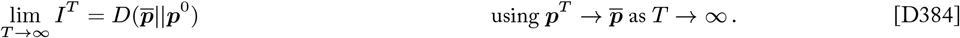

Combining Equation D381 and Equation D384, we obtain the stated result. □

##### D.4.3 Information gain is bounded by empirical regret

The following theorem establishes that the evolving population’s information gain is always bounded by the empirical regret. This bound is fully general and holds in all environmental contexts.

###### Theorem 4.

*For any game matrix **G** and for any sequence of distributions of environmental conditions* ***x***^0^, …, ***x***^*T*^, *the total information gain I*^*T* +1^ = *D*(***p***^*T* +1^||***p***^0^) *of the trajectory of type distributions* ***p***^0^, …, ***p***^*T*+1^ *generated by discrete-time replicator dynamics is at all times bounded from above by the empirical regret*

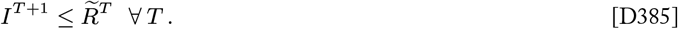

*Proof*. Proposition D3 relates the instantaneous change in potential information to the instantaneous regret with respect to some reference distribution ***q*** for discrete-time replicator dynamics.

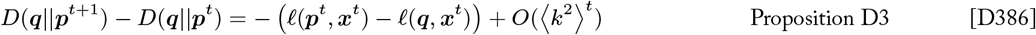

We will work from this result to establish the relationship between the total information gain and the total empirical regret. Summing over a period of selection of duration *T* we obtain

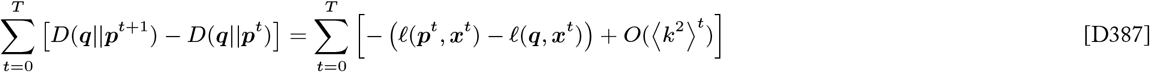

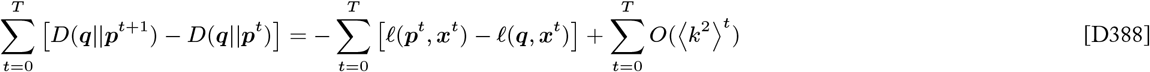

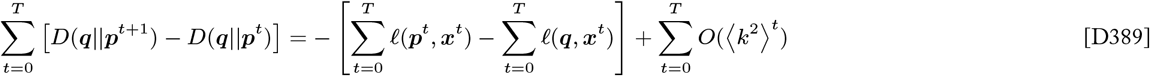

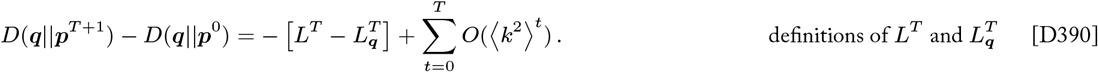

Let us evaluate with respect to the choice of reference distribution ***q*** = ***p***^*T* +1^:

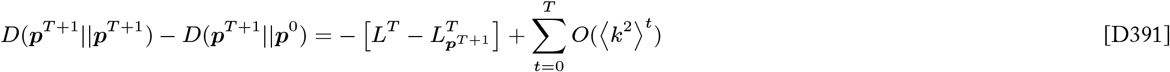

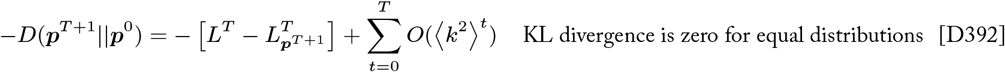

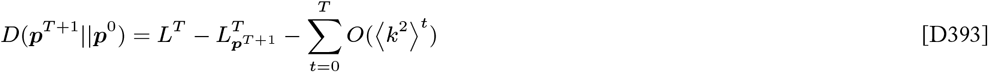

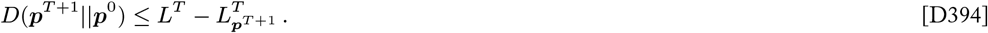

So far we have established that the information gain through *T* + 1 generations is bounded by the cumulative loss of the evolving population minus the cumulative loss of the fixed reference distribution ***p***^*T* +1^.

At this point, we recall that the empirically optimal strategy 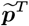 is defined as the fixed strategy with the minimum possible cumulative loss over the observed sequence of environments ***x***^0^…***x***^*T*^ in hindsight

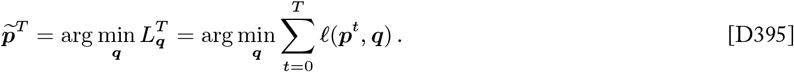

Therefore, by definition, the cumulative loss of the empirically optimal strategy 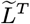 is less or equal to the cumulative loss of any other fixed strategy over the same sequence of environments

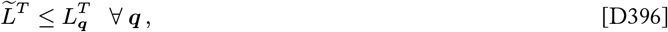

which implies that the empirically optimal cumulative loss is less than or equal to the cumulative loss of the fixed reference distribution ***p***^*T* +1^

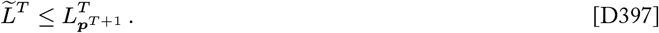

Using this fact with Equation D394, we obtain

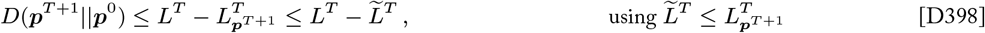

and therefore

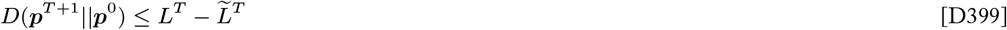

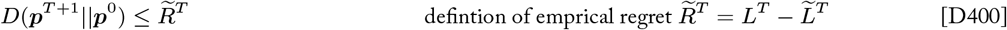

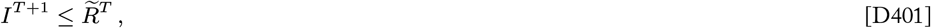

as was to be shown. □

###### Corollary 4.1.

*If the empirical regret* 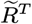 *is non-decreasing for all T, which is always the case when the population faces a fixed learning problem characterized by a stationary ESS, then*

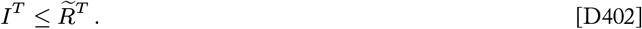

*Proof*.

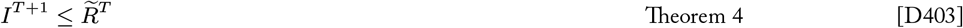

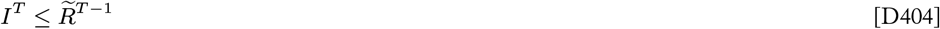

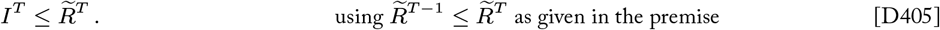

□

###### Information gain is bounded by substitution load

Substitution load is a special case of mismatch load that applies when the environment consists of a single unchanging condition (i.e., *m* = 1). In such a case, a single type has the highest fitness for all time and sweeps to fixation. Fixation of the optimal type is the empirically optimal strategy 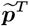 for all *T*, and the empirically optimal cumulative loss is 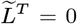 for all *T*. Therefore substitution load is equivalent to empirical regret 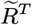 in applicable cases, and it follows from Theorem 4 that the population’s information gain is always bounded by substitution load in such cases.

###### Corollary 4.2.

*If the environment consists of a single unchanging environmental condition (m* = 1*), then the total information gain I*^*T*^ = *D*(***p***^*T*^ ||***p***^0^) *of the trajectory of type distributions* ***p***^0^, …, ***p***^*T*^ *generated by discrete-time replicator dynamics is at all times bounded from above by the substitution load*

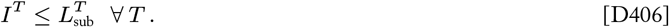

*Proof*.

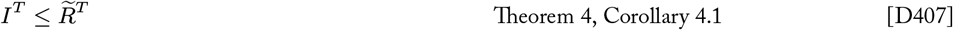

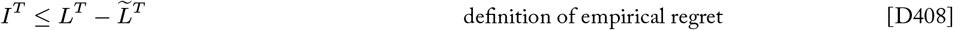

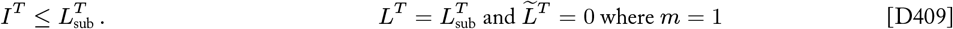

□

###### Information gain is bounded by mismatch load in general

Another notable corollary of Theorem 4 is that the information gain of a population that evolves by natural selection is always bounded by the mismatch load in the general case.

###### Corollary 4.3.

*For any game matrix **G** and for any sequence of distributions of environmental conditions* ***x***^0^, …, ***x***^*T*^, *the total information gain I*^*T*^ = *D*(***p***^*T*^ ||***p***^0^) *of the trajectory of type distributions* ***p***^0^, …, ***p***^*T*^ *generated by discrete-time replicator dynamics is at all times bounded from above by the empirical regret*

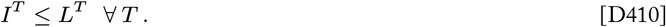

*Proof*.

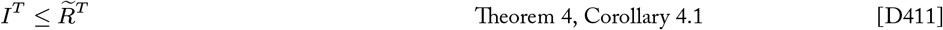

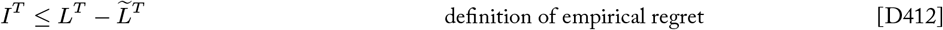

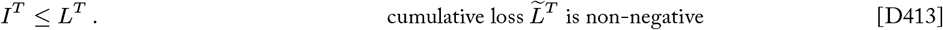

□

### D.5 Supplemental example learning problems

**Fig. D7.**
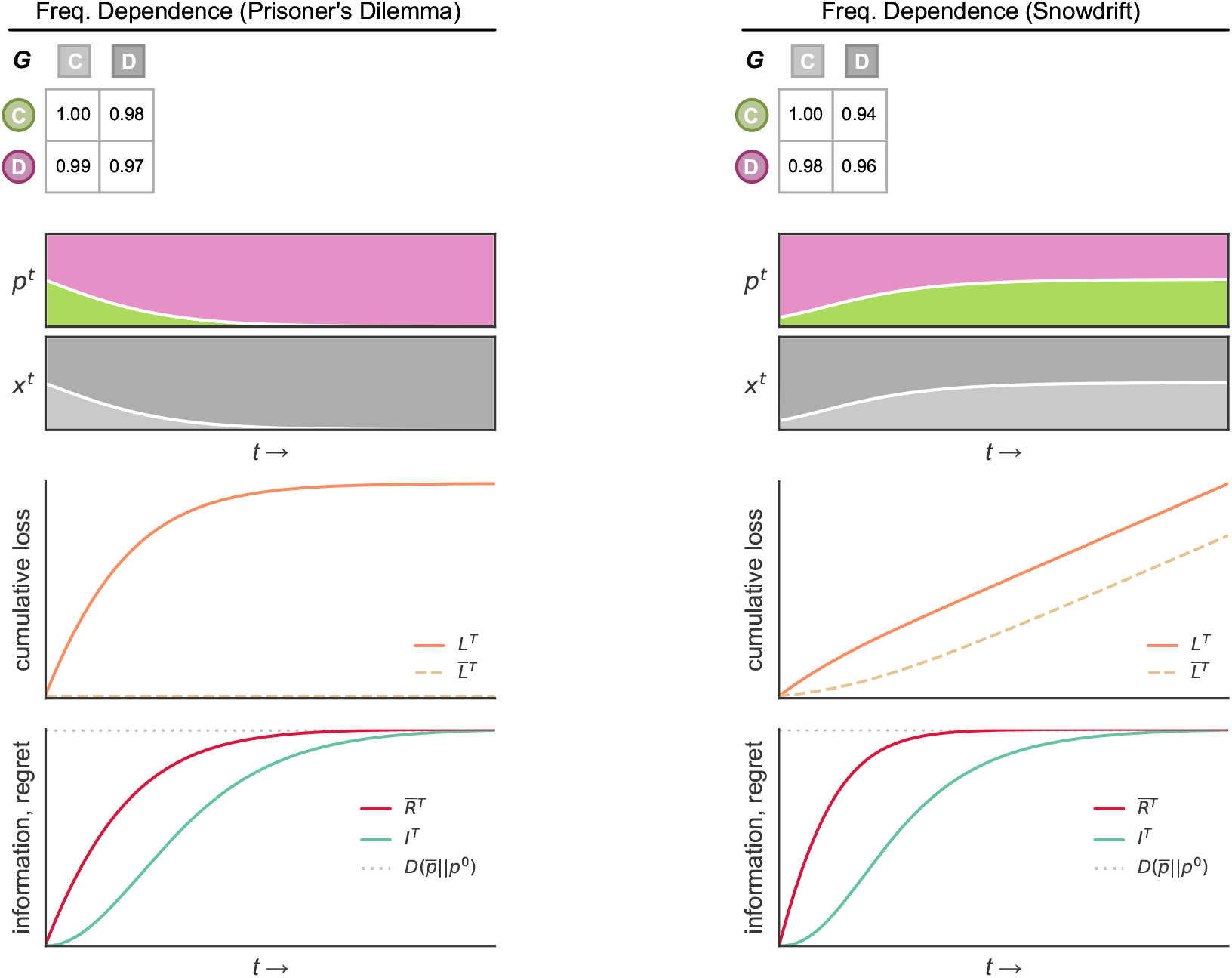
Load, regret, and information gain in different environmental contexts. Results from simulations of replicator dynamics in two additional contexts (columns) are shown. In each column, the matrix **G** defines the log fitnesses (growth rates) for 2 types in each of 2 environmental conditions. Colored Muller plots show the population’s type frequency distribution *p*^*t*^ over time, and grayscale stacked frequency plots show the distribution of environmental conditions *x*^*t*^ over time. The mismatch load of the evolving population (*L*^*T*^, orange line) and of the ESS composition (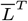, dashed gold line) are plotted over time. The bottom-most plot in each column gives the information gain (teal line) and the ESS regret (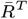, red line) over the course of selection. **(Prisoner’s Dilemma, left)** Here the environmental conditions are defined by the types (dubbed ‘Cooperate’ and ‘Defect’) themselves, and the distribution of conditions is set to the type distribution in every generation (i.e., ***x***^*t*^ = ***p***^*t*^). This frequency dependence and choice of **G** define a “Prisoner’s Dilemma” scenario. The best outcome would occur if the population plays an all-cooperate strategy, which woulds achieve the maximum possible population mean fitness. However, replicator dynamics lead toward a population of all defectors, which continually reduces the population’s mean fitness and clearly does not maximize fitness outright. If the observed sequence of environments is fixed, then the population would have been best off playing all-defect from the beginning. The all-defect strategy is a stationary ESS, and the population does indeed learn this locally optimal strategy using replicator dynamics which achieves vanishing per-round regret. **(Snowdrift, right)** Here again the environmental conditions are defined by the types (‘Cooperate’ and ‘Defect’) themselves, and the distribution of conditions is set to the type distribution in every generation (i.e., ***x***^*t*^ = ***p***^*t*^). This frequency dependence and choice of **G** define a “Snowdrift” (aka “Hawk-Dove”) game. Replicator dynamics carry the population to a polymorphic ESS where the expected fitness of cooperators and defectors is equal. The empirically optimal strategy, which globally minimizes expected loss (maximizes expected fitness) for the observed sequence of environments (given by the type frequencies in this context), is to play all-defect due to the predominance of defectors throughout the environmental history until equilibrium is reached. This is an example of a sequence of environments where the ESS and the empirically optimal strategy differ.

### Appendix E: Selection Experiments

#### E.1 Strain information

*Escherichia coli* B (REL606) strains were used in selection experiments and related assays. A “wild type” strain (WT) and three strains with unique mutations in the *rpoB* gene (M1, M2, M3) were obtained with permission from the −80^°^C strain archive from Lindsey et al. (2013). Mutations to the *rpoB* gene conferred each mutant strain with a distinct exponential growth rate that was reduced from that of the WT strain (Figure E1).

Each strain was transformed with an engineered marker plasmid. The pBR322 plasmid, which carries a *bla* gene conferring ampicillin resistance (Amp^*R*^) and a *tetA* gene conferring tetracycline resistance (Tet^*R*^), served as a vector backbone. Using Gibson assembly, the *bla* gene and corresponding promoter region was removed and replaced with an insert carrying a fluorescent protein gene under a strong constitutive *proC* promoter. As the strain with the optimal growth rate, the WT strain was transformed with a plasmid engineered to carry the green fluorescent protein (GFP) gene *mGFPmut2*. The *rpoB* mutant strains were each transformed with plasmids engineered to carry the red fluorescent protein (RFP) gene *mScarlet-I*.

Strains and culture samples were stored by mixing 1 ml of culture with 160 *μ*l of 80% glycerol and freezing at −80^°^C. Strains were revived from freezer scrapings and incubated overnight prior to all assays and experiments.

**Table E1.** Basic characteristics of bacterial strains used in selection experiments.

| Strain ID | Chromosomal Genotype |  |  | Plasmid |  |  |  |
| --- | --- | --- | --- | --- | --- | --- | --- |
|  | *E. coli* strain | Mutations | Selective Markers | Plasmid ID | Vector Backbone | FP Marker Gene Insert | Selective Markers |
| WT | B (REL606) | See Table E2 | - | pRMbcGFP | pBR322 | pEB1-mGFPmut2 | $\text{Tet}^R$ |
| M1 | B (REL606) | See Table E2 | $\text{Rif}^R$ | pRMbcRFP | pBR322 | pEB2-mScarlet-I | $\text{Tet}^R$ |
| M2 | B (REL606) | See Table E2 | $\text{Rif}^R$ | pRMbcRFP | pBR322 | pEB2-mScarlet-I | $\text{Tet}^R$ |
| M3 | B (REL606) | See Table E2 | $\text{Rif}^R$ | pRMbcRFP | pBR322 | pEB2-mScarlet-I | $\text{Tet}^R$ |

**Table E2.** Mutations present in each bacterial strain relative to the *Escherichia coli* B (REL606) reference genome (presence of a mutation in a given strain is indicated by an X in that strain’s column).

| WT | M1 | M2 | M3 | Position | Gene | Mutation | Annotation | Description |
| --- | --- | --- | --- | --- | --- | --- | --- | --- |
| × | × | × | × | 3,925,216 | *gppA* ← | C→T | L272L (CTG→CTA) | guanosine pentaphosphatase |
| × |  |  |  | 458,185 | *acrR* → | IS1 (+) +9 bp | Coding (358 366/648 nt) | DNA binding transcriptional repressor |
| | × | | | 4,162,387 | *rpoB* → | T→G | L511R (CTG→CGG) | DNA-directed RNA polymerase subunit $\beta$ |
| | | × | | 4,161,298 | *rpoB* → | A→T | Q148L (CAG→CTG) | DNA-directed RNA polymerase subunit $\beta$ |
| | | × | | 4,162,382 | *rpoB* → | C→A | S509R (AGC→AGA) | DNA-directed RNA polymerase subunit $\beta$ |
| | | | × | 4,161,292 | *rpoB* → | T→A | V146D (GTT→GAT) | DNA-directed RNA polymerase subunit $\beta$ |
| | | | × | 4,162,540 | *rpoB* → | A→C | E562A (GAA→GCA) | DNA-directed RNA polymerase subunit $\beta$ |

**Fig. E1.**
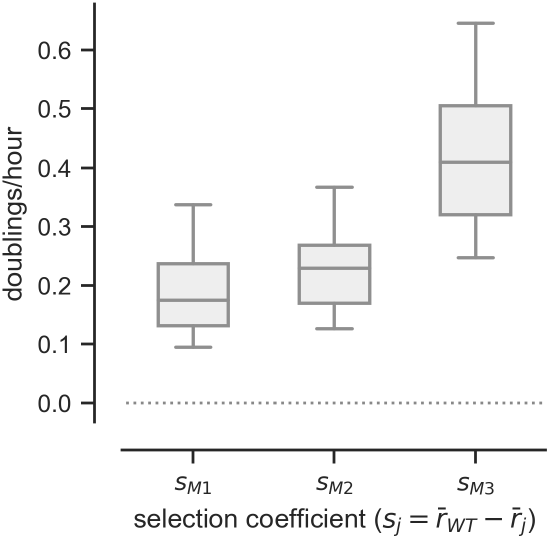
Selection coefficients. The selection coefficient is a measure of the fitness of a type (i.e., strain) relative to another. In Malthusian fitness terms, the selection coefficient of a type *j* is defined as the difference between the exponential growth rate of the optimal type and that of the *j*th strain; that is, *s*_*j*_ = *r*_*∗*_ *− r*_*j*_ (Appendix A.1.2). In our experimental system, the WT strain had the fastest growth rate. Therefore the selection coefficient of a strain *j* was defined as *s*_*j*_ = *r*_WT_ *− r*_*j*_. A positive selection coefficient *s*_*j*_ *>* 0 indicates that the growth rate of the *j*th strain is less than that of the WT strain. By definition, the selection coefficient of the WT strain is zero (*s*_WT_ = 0). Here we show the selection coefficients of the mutant strains (M1, M2, M3) relative to the WT strain as measured by an exponential growth assay. Individual strain cultures were sampled and transferred to fresh media to maintain exponential growth every 2 hours for a total of 8 hours. Cell enumeration was performed on culture samples using flow cytometry, and strain growth rates were calculated from changes in culture density over the course of the assay. Selection coefficients were then calculated according to the definition given above. These selection coefficients reflect the relative growth rates of strains after being transformed with the corresponding marker plasmids.

**Fig. E2.**
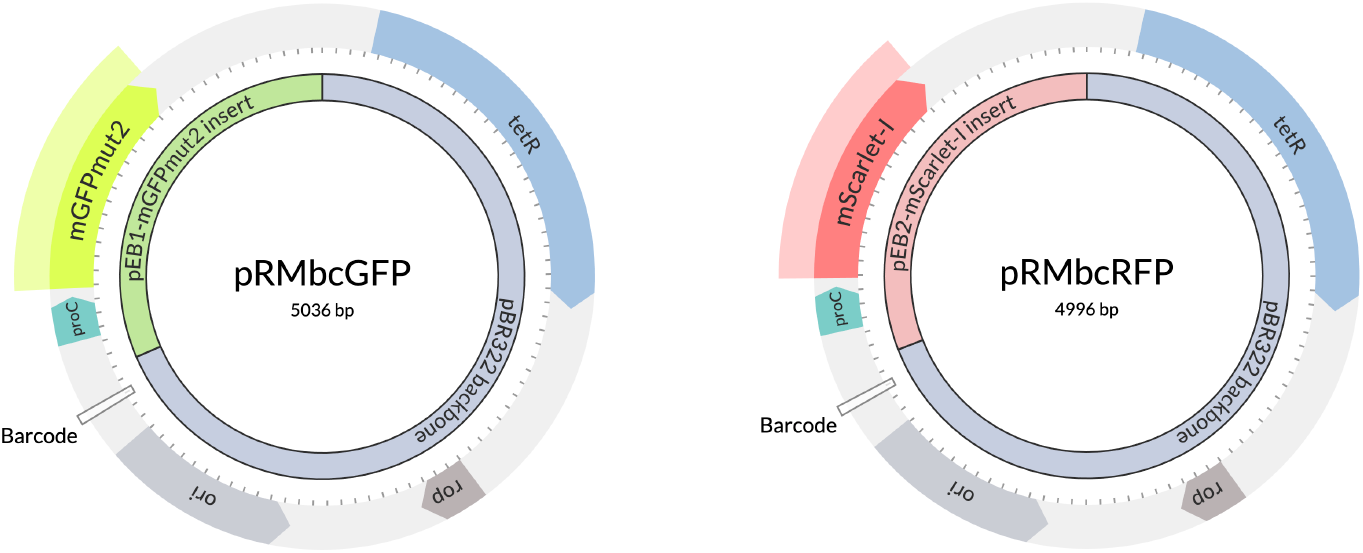
Engineered plasmid maps. Each strain was transformed with an engineered marker plasmid. The pBR322 plasmid, which carries a *bla* gene conferring ampicillin resistance (Amp^*R*^) and a *tetA* gene conferring tetracycline resistance (Tet^*R*^), served as a vector backbone. Using Gibson assembly, the *bla* gene and corresponding promoter region was removed and replaced with an insert carrying a fluorescent protein gene under a constitutive *proC* promoter. **Left:** The pRMbcGFP plasmid carries a region encoding the green fluorescent protein gene *mGFPmut2* under the *proC* promoter, which was taken from the pEB1-mGFPmut2 plasmid (Addgene plasmid #103980). **Right:** The pRMbcRFP plasmid carries a region encoding the red fluorescent protein gene *mScarlet-I* under the *proC* promoter, which was taken from the pEB2-mScarlet-I plasmid (Addgene plasmid #104007).

#### E.2 Supplemental methods

Selection competition experiments were conducted between three pairs of strains: WT vs. M1, WT vs. M2, and WT vs. M3. Each pair of strains participated in three competitions with different initial strain frequency compositions: WT at approximately 12.5%, 25%, and 50% of the initial population, respectively.

Strains were cultured in standard Luria-Bertani (LB) broth with with 15*μ*g/mL tetracycline for plasmid retention (hereafter referred to as “media”). Cultures were incubated in 25 mL of media in 125 mL capacity flasks at 37^°^C with shaking. A single large batch of media was prepared and used for all cultures across all stages of all experiments, and all competitions were conducted simultaneously using the same incubator in order to mitigate potential day or block effects.

Strains were revived and incubated overnight to saturation density (∼1×10^9^ cfus/mL). Prior to the start of competitions, individual strain cultures were diluted 1000-fold into 25 mL of fresh media (∼1 × 10^6^ cfus/mL) and incubated for 1 hour, which was sufficient time for cells to begin resuming exponential growth. At the end of this “reanimation” period, the density of each strain culture was spot-checked using flow cytometry. Selection experiments were initiated by combining the strains to be competed in the designated ratios at a total density of ∼1 × 10^6^ cfus/mL in 25 mL of fresh media. Because the strains had entered exponential growth phase prior to initiating competitions, some growth occurred during the initiation process, which caused minor imprecisions in the initial strain frequencies.

Every 4 hours, a 1 mL sample was taken from each competition culture to measure strain densities and frequencies using flow cytometry (see Appendix E.2.2 for more information). At the same time, a sample of the culture was transferred to 25 mL of fresh media. An adaptive protocol was used to determine the transfer volume for each competition culture at each transfer in order to ensure that the culture would remain in exponential growth while maintaining a measurable density for flow cytometry measurements in the next growth interval (see Appendix E.2.1 for more information). Competitions proceeded in this fashion for 36 hours.

##### E.2.1 Transfer protocol

Selection experiments involved incubating mixed batch cultures and tracking the densities and frequencies of the competing strains for a total of 36 hours, which was the time necessary to observe the optimal WT strain approaching fixation in all competitions. These strains enter a stationary phase as the culture approaches a saturation density of approximately 1 × 10^9^ cfus/mL, so regular transfers to fresh media were necessary to maintain active growth and selection over this duration. Basic transfer protocols often involve repeatedly growing cultures to saturation and transfering a sample of the saturated culture to fresh media at low density (e.g., ∼1 × 10^5^ cfus/mL). However, we were interested in measuring the outcomes of selection due to differential exponential growth rates, and thus we wished to maintain cultures in exponential phase to avoid the potential confounding effects of cells entering and exiting lag phase. In addition, highly accurate density estimation using flow cytometry required a culture density of at least ∼1 × 10^7^ cfus/mL at the time of sampling. Therefore, in order to ensure that the culture would remain in exponential growth while maintaining a measurable density for the flow cytometer we required a transfer protocol that would reliably maintain culture densities between ∼1×10^7^ −1×10^8^ cfus/mL at the end of each growth interval. The following transfer protocol was developed to dynamically calculate the transfer volume for each competition culture in order to achieve this (note that all competitions were conducted simultaneously).

**Fig. E3.**
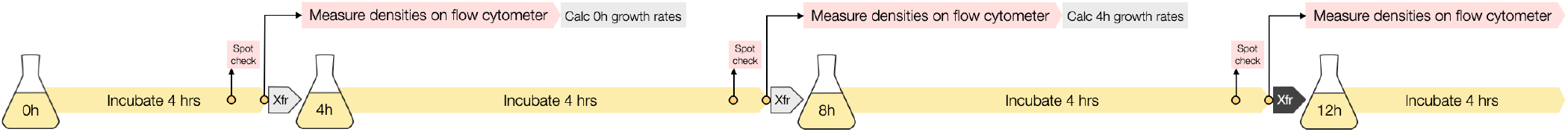
Illustration of transfer protocol.

To make things concrete, we outline the transfer protocol from the perspective of an experimenter determining transfer volumes at the 12 hour mark of the overall experiment (black-shaded block arrow in Figure E3), although the same process was used at all transfer points throughout the 36 hour experiment. Flasks are labeled by the time point at which they were inoculated (e.g., the ‘0h’ culture flask was inoculated at the beginning of the experiment, the ‘4h’ culture flask was inoculated via transfer at the 4 hour mark, and so on).

- **4 hours prior to transfer:** Data collection samples are taken from each competition, and competition cultures are transferred to fresh media.
  1. Samples are taken from the previous set of culture flasks (e.g., the ‘4h’ cultures) for density estimation on the flow cytometer.
  2. The current set of flasks (e.g., the ‘8h’ cultures) are inoculated via transfer and their incubation period begins.
  3. Cytometry samples (e.g., of the ‘4h’ cultures) are prepared and measured over the following 1-2 hours.
  4. The newly obtained cytometry measurements provide endpoint density estimates for the the previous set of culture flasks (e.g., the ‘4h’ cultures). The initial density of these same cultures is estimated from the endpoint density of the prior set of cultures (e.g., obtained from cytometry measurements of the ‘0h’ cultures) and the transfer volumes used at that time. The.growth rates of each of the previous cultures (e.g., the ‘4h’ cultures) is estimated using the initial and endpoint densities for this interval (e.g., the 4h-8h interval).
- **30 minutes prior to transfer:** A “spot check” is performed on a subset of the current flasks in order to obtain a rough estimate of the densities and growth rates of the currently incubating cultures.
  1. Samples are quickly taken from the competition cultures with the lowest growth rate and the highest growth rate observed in the latest set of cytometry-based growth rate estimates (e.g., from the ‘4h’ cultures). These flasks are immediately returned to the incubator to finish the rest of the incubation period.
  2. These samples are enumerated on the flow cytometer and used to estimate the endpoint densities and growth rates of the presumed fastest and slowest growing competition cultures in the current incubation period. This provides a presumed minimum and maximum growth rate for all competitions in the current growth interval.
  3. The endpoint densities for all competition cultures in the current growth interval are projected based on the initial densities of these cultures (using the previous cultures’ endpoint densities and the transfer volumes involved in their inoculations) and the average of the growth rates estimated from the spot check measurement described above.
- **10 minutes prior to transfer:** The transfer volume to be used for each competition culture is calculated as follows (a spreadsheet was programmed to perform these calculations automatically using flow cytometry data collected previously in the experiment):
  – For each competition culture, working in order of highest projected endpoint density (from #3 above) to lowest:
    1. If this is the first competition considered, let the “trial” transfer volume for this competition culture be a default volume of 250 *μ*L (i.e., a 100-fold dilution into 25 mL of media); else, let the “trial” transfer volume be the actual transfer volume of the previously considered competition.
    2. Calculate the projected initial density of the post-transfer culture based on the projected final density of the current culture and the “trial” transfer volume.
    3. Calculate the projected endpoint density of this competition culture in the next incubation period based on the projected initial density and the *minimum* growth rate that has been observed for any culture so far. Likewise, calculate the projected endpoint density of this competition culture in the next incubation period using the *maximum* growth rate that has been observed for any culture so far. This gives the range of plausible endpoint densities the post-transfer culture may realize if the “trial” transfer volume is used.
    4. If the range of plausible endpoint densities using the “trial” transfer volume falls within the acceptable range of endpoint densities (i.e., 1 × 10^7^ − 1 × 10^8^ cfus/mL), then use the “trial” transfer volume to transfer this competition culture; else, do the following:
      * Backcalculate the initial density that would lead this culture to just reach the *minimum* acceptable endpoint density (i.e., 1×107 cfus/mL) if the culture were to grow at the *minimum* growth rate that has been observed for any culture so far. This is the smallest post-transfer initial density that is ensured to reach the minimum acceptable endpoint density in a worst-case, slow growth scenario (based on the range of growth rates observed to this point). (This heuristic is used because we deemed it more important to guarantee that cultures reach accurately measurable densities than to guarantee that cultures do not approach saturation.)
      * Calculate the transfer volume that would give this initial density based on the projected endpoint density for this culture in the current incubation period.
      * Use this “backcalculated” transfer volume to transfer this competition culture.
- **At time of transfer:** Data collection samples are taken from each competition, and competition cultures are transferred to fresh media using the calculated transfer volumes. The cycle repeats for the following incubation period and transfer.

This transfer protocol successfully maintained endpoint densities for all competition cultures within the range 8.3×10^6^ −2.7×10^8^ cfus/mL throughout the entire 36 hour experiment, with only a few culture endpoints falling outside of the tight target range of 1 × 10^7^ − 1 × 10^8^ cfus/mL. All culture samples had densitites that were accurately measurable on the flow cytometer, and no competition culture showed signs of cells exiting exponential phase to any appreciable degree.

##### E.2.2 Flow cytometry

Flow cytometry was used to enumerate cells in culture samples and estimate culture densities. Fixed fluorescence gates were established to separately enumerate cells expressing GFP and RFP markers. Culture samples were immediately centrifuged at 13,000 rpm for 5 minutes to remove growth media supernatant. Pelleted cells were then resuspended in 1 mL of flow buffer (1X PBST with EDTA added at a 500-fold dilution). Samples were then transferred to a round-bottom 96-well plate where a 10-fold dilution series was performed. The accuracy of cell enumeration was maximized when the sample density was in the range of ∼1 × 10^4^ − 1 × 10^6^ cfus/mL, and samples were measured at 10-fold, 100-fold, and 1000-fold dilutions to ensure each sample had measurements in this range. Three replicate flow cytometer enumerations were performed for each sample at each dilution.

**Fig. E4.**
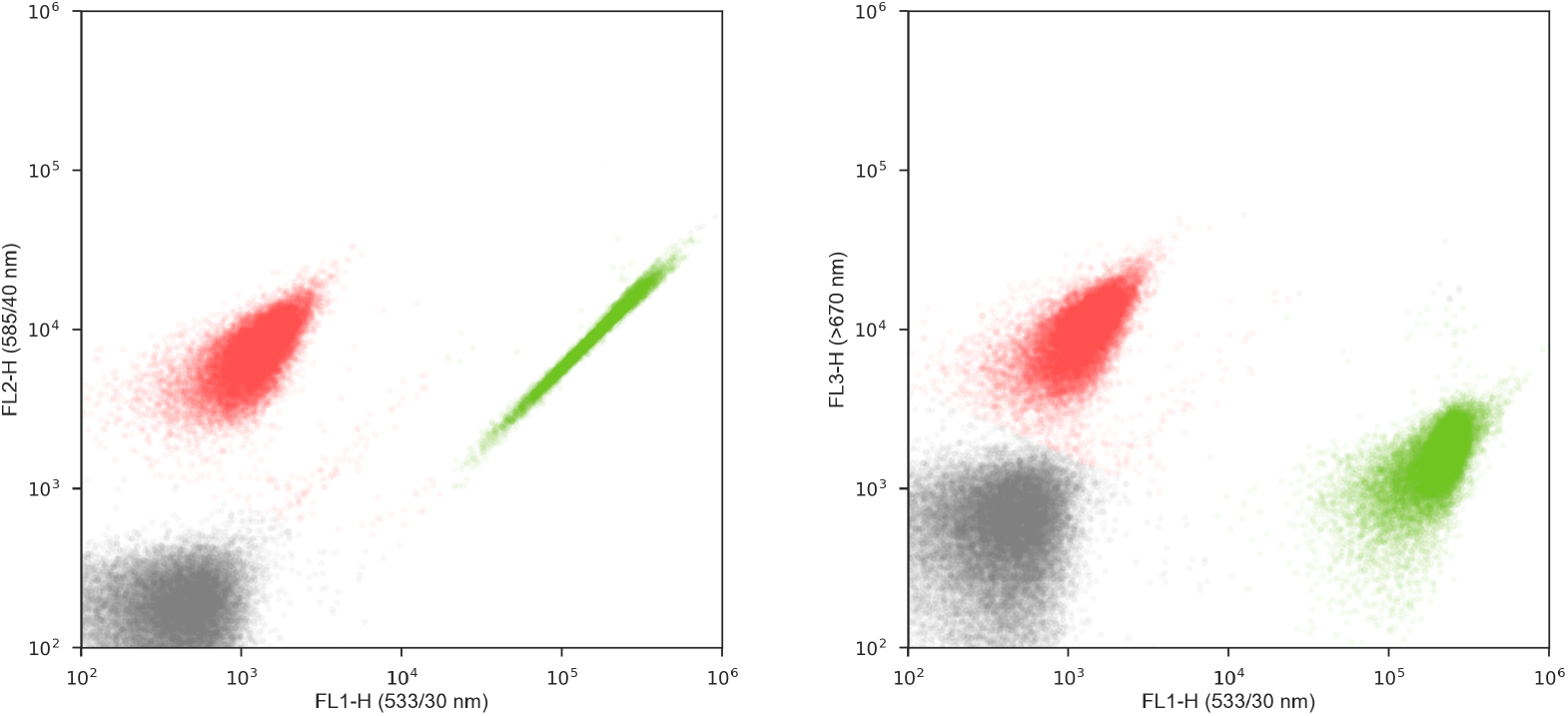
Example flow cytometry data. Culture densities were estimated with cell enumeration using a BD Accuri C6 Flow Cytometer. These plots present example data collected from a single sample taken from a mixed culture of GFP-marked WT cells and RFP-marked M1 cells. The x- and y-axes refer to fluorescence intensities in the wavelength bands indicated in parentheses. Each point represents the fluorescence intensities of a particle drawn from the culture sample. Three distinct clusters of points are immediately apparent, and fixed fluorescence gates were established to automatically differentiate GFP-marked cells, RFP-marked cells, and other particles. Cells that fall within the RFP gate are colored red above, cells that fall within the GFP gate are colored green, and other particles that fall outside of both FP gate regions are colored gray. Such data provides counts of GFP-marked cells, RFP-marked cells, and total cells, which can be used to calculate the strain densities and frequencies in the culture from which the cytometry sample was taken.

**Fig. E5.**
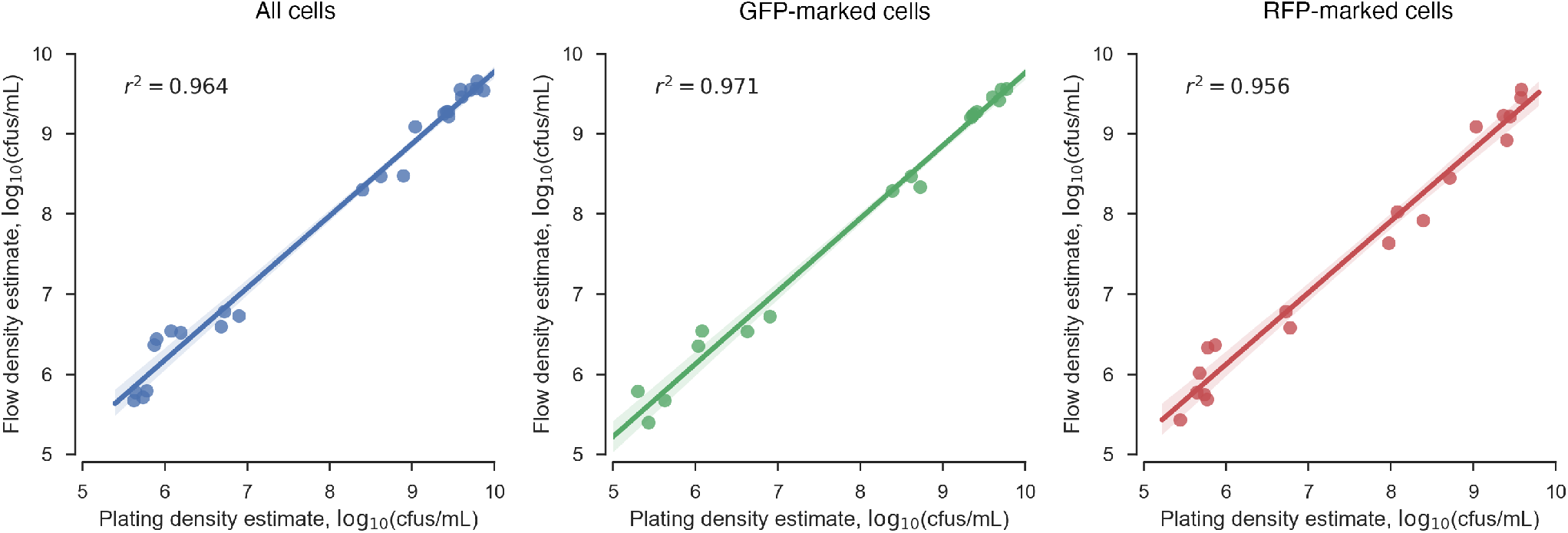
Correspondence of cell enumeration using flow cytometry vs. agar plating. Mixed cultures of GFP-marked cells and RFP-marked cells were grown from low density (∼1 *×* 10^6^ cfus/mL) to saturation (∼2 *×* 10^9^ cfus/mL). Samples of these cultures were taken throughout the growth process with densities spanning several orders of magnitude. Culture densities were estimated both with cell enumeration using a BD Accuri C6 Flow Cytometer and with traditional agar plating and colony counting (colonies of GFP-marked cells appear green, and colonies of RFP-marked cells appear pink). These plots show how density estimates from flow cytometry compare to those from plating. Each point depicts the respective density estimates for the same culture sample. The leftmost plot shows the correspondence of density estimates for all cells, the middle plot shows the correspondence of density estimates for GFP-marked cells alone, and the rightmost plot shows the correspondence of density estimates for RFP-marked cells alone. In all cases, the correlation of cytometry- and plating-based estimates of culture density is high.

#### E.3 Supplemental results of selection experiments

**Fig. E6.**
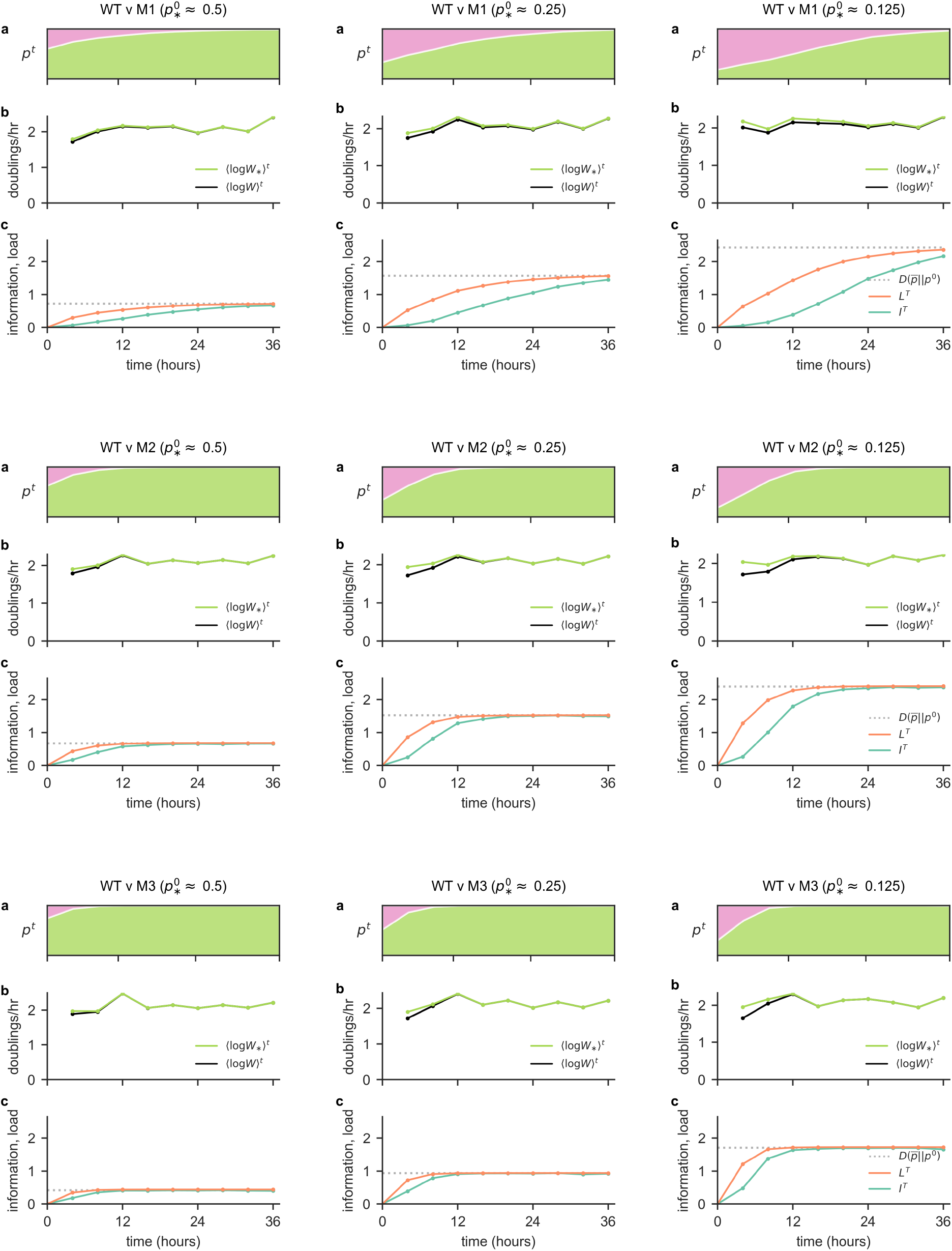
Supplemental results from selection experiments. Here we show the changes in strain frequencies (**a** plots) and growth rates (**b** plots), and information and load measures (**c** plots) over time for each of the 9 selection competitions presented in main text Figure 3. The title of each sub-panel indicates the strain combination and approximate initial frequency of the optimal WT strain in the respective competition.

## Footnotes

* The main text presents bounds for the limit of weak selection. Appendix D presents bounds in full generality, which differ from those in the main text by only a small gap term that vanishes with decreasing selection strength.

* Different environmental *histories* (*E*) are not to be confused with different *conditions* (*j*) within an environment (***x***^*t*^). In terms of the framework studied here, environmental histories can be interpreted as different sequences of environments {***x***^0^, ***x***^1^, …, ***x***^*T*^ }

* The Population versus Environment game model is wholly consistent with the models of replicator dynamics in variable environments outlined in Appendix A.2.

† Note that this differs from standard evolutionary game theory where each individual within the population is a separate player.

* Replicator dynamics expressed here in terms of the expected relative fitness ⟨*w*_*i*_⟩^*t*^ of each type *i* over all environmental conditions.

* Haldane (1957) considered a 2 allele model, but it is straightforward to extend this to *n* types.

* We extend the 2 allele model from Kimura (1960, 1961) to *n* types.

* Note that any loss function can be scaled to the range [0, 1] without loss of generality.

